# Protocol: Genome-scale CRISPR-Cas9 Knockout and Transcriptional Activation Screening

**DOI:** 10.1101/059626

**Authors:** Julia Joung, Silvana Konermann, Jonathan S. Gootenberg, Omar O. Abudayyeh, Randall J. Platt, Mark D. Brigham, Neville E. Sanjana, Feng Zhang

**Author notes:** These authors contributed equally to this work. Correspondence should be addressed to (F.Z.).

## Abstract

Forward genetic screens are powerful tools for the unbiased discovery and functional characterization of specific genetic elements associated with a phenotype of interest. Recently, the RNA-guided endonuclease Cas9 from the microbial immune system CRISPR (clustered regularly interspaced short palindromic repeats) has been adapted for genome-scale screening by combining Cas9 with guide RNA libraries. Here we describe a protocol for genome-scale knockout and transcriptional activation screening using the CRISPR-Cas9 system. Custom-or ready-made guide RNA libraries are constructed and packaged into lentivirus for delivery into cells for screening. As each screen is unique, we provide guidelines for determining screening parameters and maintaining sufficient coverage. To validate candidate genes identified from the screen, we further describe strategies for confirming the screening phenotype as well as genetic perturbation through analysis of indel rate and transcriptional activation. Beginning with library design, a genome-scale screen can be completed in 6-10 weeks followed by 3-4 weeks of validation.

## INTRODUCTION

Systematic and high-throughput genetic perturbation technologies within live model organisms are necessary for fully understanding gene function and epigenetic regulation^1–3^. Forward genetic screens allow for a “phenotype-to-genotype” approach to mapping specific genetic perturbations to a phenotype of interest. Generally, this involves perturbing many genes at once, selecting cells or organisms for a desired phenotype, and then sequencing to identify the genetic features involved in the phenotypic changes. Initial screening approaches relied on chemical DNA mutagens to induce genetic changes, but this process was inefficient and mutations were costly to identify. More recently, tools that utilize the RNA interference (RNAi) pathway, specifically through short hairpin RNAs (shRNAs)^4–7^, to perturb transcript levels have revolutionized screening approaches^8–13^. ShRNAs exploit the endogenous RNAi machinery to knock down sequence-complementary mRNA (**Fig. 1**). Despite the contribution of RNAi screens to many biological advances, this approach is hampered by incomplete knockdown of transcripts and high off-target activity, resulting in low signal-to-noise and limited interpretations^14–16^.

**Figure 1.**
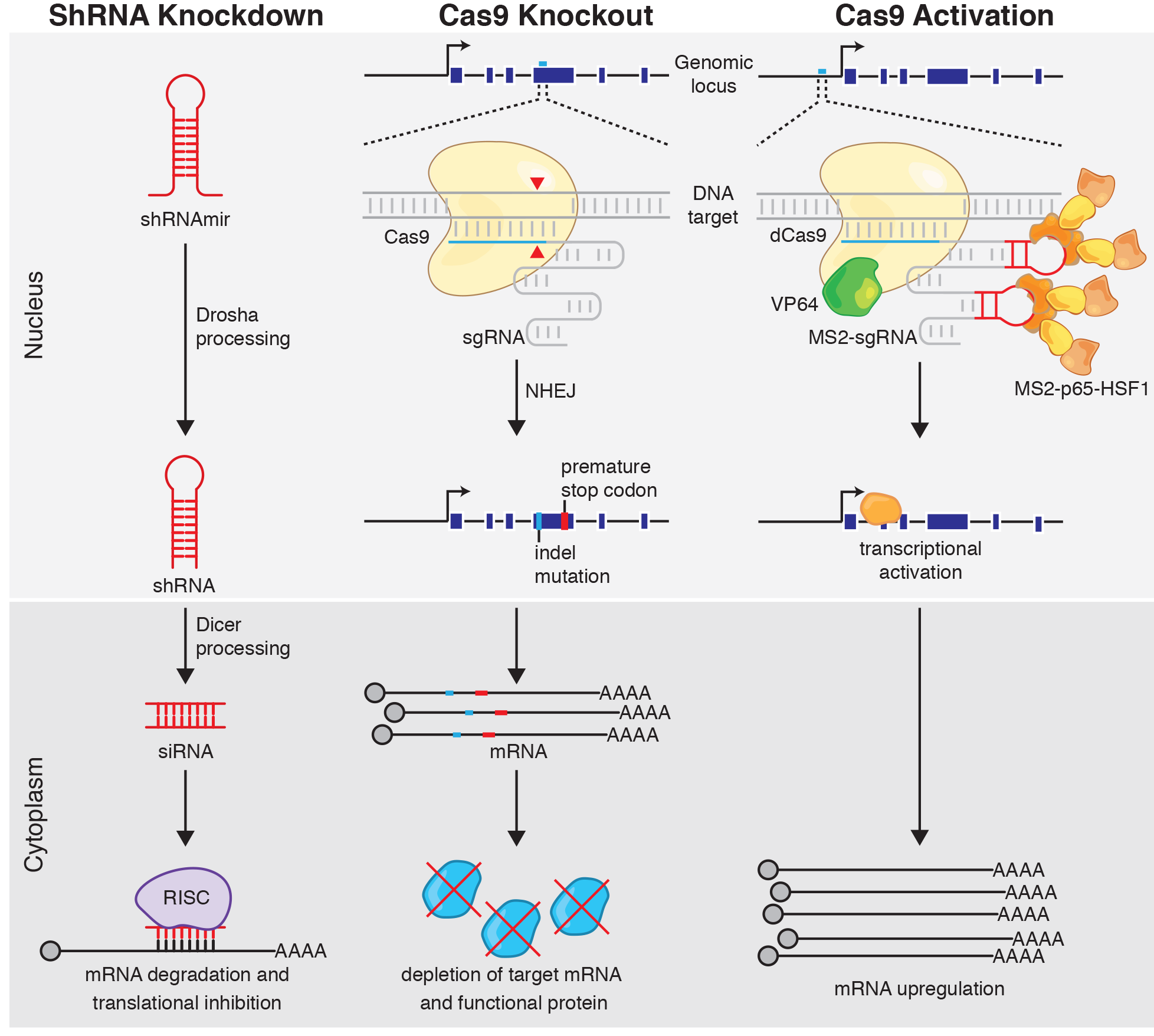
Approaches to genetic perturbation: shRNA knockdown, Cas9 knockout, and Cas9 transcriptional activation. Schematic of the mechanisms behind shRNA knockdown, Cas9 knockout, and Cas9 transcriptional activation. ShRNA knockdown begins with processing of the shRNA by Drosha/Dicer machinery and results in degradation of an RNA transcript with a complementary target site by the RISC complex. Cas9 knockout is accomplished by targeted indel formation at a genomic site complementary to the sgRNA. An indel can result in a frameshift, causing early termination, and either production of non-functional protein or non-sense mediated decay of the mRNA transcript. Programmable transcriptional activation can be achieved using dCas9 and activation domains (e.g. VP64/p65/HSF1) to recruit transcriptional machinery to the transcriptional start site of the desired gene target, resulting in upregulation of the target transcript.

### Cas9 as a tool for precise genome editing

Programmable nucleases have emerged as a promising new genetic perturbation technology capable of precisely recognizing and cleaving target DNA^17–19^. Particularly, the RNA-guided endonuclease Cas9 from the microbial immune system CRISPR (clustered regularly interspaced short palindromic repeat) has proved powerful for precise DNA modifications^20–23^. Cas9 is guided to specific genomic targets by guide RNAs through Watson-Crick base pairing. As with RNAi, Cas9 is thus easily retargetable; in contrast, however, extensive characterization has shown that Cas9 is much more robust and specific than RNAi^24–28^.

Cas9 generates precise double-strand breaks (DSBs) at target loci that are repaired through either homology-directed repair (HDR) or more often, non-homologous end-joining (NHEJ)^29^. HDR precisely repairs the DSB using a homologous DNA template, whereas NHEJ is error-prone and introduces indels. When Cas9 is targeted to a coding region, loss-of-function (LOF) mutations can occur as a result of frameshifting indels that produce a premature stop codon and subsequent nonsense-mediated decay of the transcript or generate a non-functional protein (**Fig. 1**)^22, 23^. These features make Cas9 ideal for genome editing applications.

### High-throughput loss-of-function screening using Cas9

Together with large pooled single guide RNA (sgRNA) libraries, Cas9 can mediate high-throughput LOF dissection of many selectable phenotypes. Indeed, Cas9 LOF screens have provided insight into the molecular basis of gene essentiality, drug and toxin resistance, as well as the hypoxia response^27, 30–42^. Previously, we constructed a genome-scale CRISPR-Cas9 knockout (GeCKO) library to probe 18,080 genes in the human genome for roles in BRAF-inhibitor vemurafenib resistance in a melanoma cell line^27^. A comparison of GeCKO with shRNA screening indicated that GeCKO had higher levels of consistency amongst guides targeting the same gene and higher validation rates.

Similarly, Cas9 knockout screening has been shown to be more consistent and effective than shRNA screening for the identification of essential genes^28^. The Cas9 system is also effective for screening *in vivo*. For instance, Chen et al. identified key factors in cancer metastasis by infecting a non-metastatic mouse lung cancer cell line with a mouse GeCKO library, transplanting it into mice, and sequencing metastases in the lung^34^.

### Genome-scale transcriptional activation with Cas9

In addition to Cas9-based knockout screens, catalytically inactive Cas9 (dCas9) fused to transcriptional activation and repression domains can be used to modulate transcription without modifying the genomic sequence^41–48^. CRISPR activation (CRISPRa) and CRISPR inhibition (CRISPRi) can be achieved by direct fusion or recruitment of activation and repression domains, such as VP64 and KRAB respectively^44, 49^. Of these alternative CRISPR screening approaches, CRISPRa is perhaps the most robust and reliable. Up until CRISPRa, gain-of-function (GOF) screens were primarily limited to cDNA overexpression libraries, which suffered from incomplete representation, overexpression beyond physiological levels and endogenous regulation, lack of isoform diversity, and high cost of construction. CRISPRa overcomes these limitations because it activates gene transcription at the endogenous locus and simply requires the synthesis and cloning of small sgRNAs.

The first generation of CRISPRa fused dCas9 to a VP64 or p65 activation domain to produce modest transcriptional upregulation, but was not suitable for genome-scale screening^44–47, 49^. Second generation CRISPRa designs produced more robust upregulation by recruiting multiple activation domains to the dCas9 complex. For instance, SunTag recruits multiple VP64 activation domains via a repeating peptide array of epitopes paired with single-chain variable fragment antibodies^41^. Another activation method, VPR, uses a tandem fusion of three activation domains, VP64, p65, and Rta to dCas9 to enhance transcriptional activation^43^. We devised an alternative approach to CRISPRa that involved incorporating MS2 binding loops in the sgRNA backbone to recruit several different activation domains, p65 and HSF1, to a dCas9-VP64 fusion (**Fig. 1**)^42^. By recruiting three distinct transcriptional effectors, this synergistic activation mediator (SAM) complex could robustly and reliably drive transcriptional upregulation.

As a proof of principle to demonstrate SAM's utility for GOF screening, similar to GeCKO we performed a genome-scale GOF screen to identify genes that, upon activation, confer vemurafenib resistance in a melanoma cell line. A comparison of SunTag, VPR, and SAM across various cell types and species suggested that SAM is the most robust and effective transcriptional activator^50^.

### Screening strategies

In general, there are two formats for conducting a screen: arrayed and pooled. For arrayed screens, individual reagents are aliquoted into separate wells in multi-well plates. This format allows for a diverse range of measured phenotypes such as fluorescence, luminescence, or even direct imaging of cellular phenotypes^2, 51–53^, but at the same time it is costly and time-consuming. An alternative format, and one that has been widely used for Cas9-based screens, is pooled screening in which pooled lentiviral libraries are transduced at a low multiplicity of infection (MOI) to ensure that most cells receive only one stably-integrated RNA guide. After the screen is complete, deep sequencing of the sgRNAs in the bulk genomic DNA identifies changes in the sgRNA distribution due to the applied screening selection pressure. While pooled screens are generally limited to growth phenotypes or to fluorescence-activated cell sorting (FACS)-selectable phenotypes, they require less cost and effort than arrayed screens. Additional design considerations for screening include the type of selection pressure applied. This can be categorized as positive (e.g. resistance to a drug, toxin, or pathogen), negative (e.g. essential genes, toxicity), or marker gene selection (e.g. reporter gene expression) (**Box 1**).

Here we explain in detail how to set up and perform a pooled genome-scale knockout and transcriptional activation screen using Cas9. We describe protocols for designing and cloning an sgRNA library, packaging lentivirus for transduction, analyzing screening results, and validating candidate genes identified from the screen (**Fig. 2**). Although we specifically focus on knockout and activation screening using the GeCKO and SAM systems, the protocol can be applied to other types of screens (e.g. other CRISPRa systems, Cas9 knockdown, and saturated mutagenesis). For reference, we have compiled a table of previously published screens (**Table 1**).

**Figure 2.**
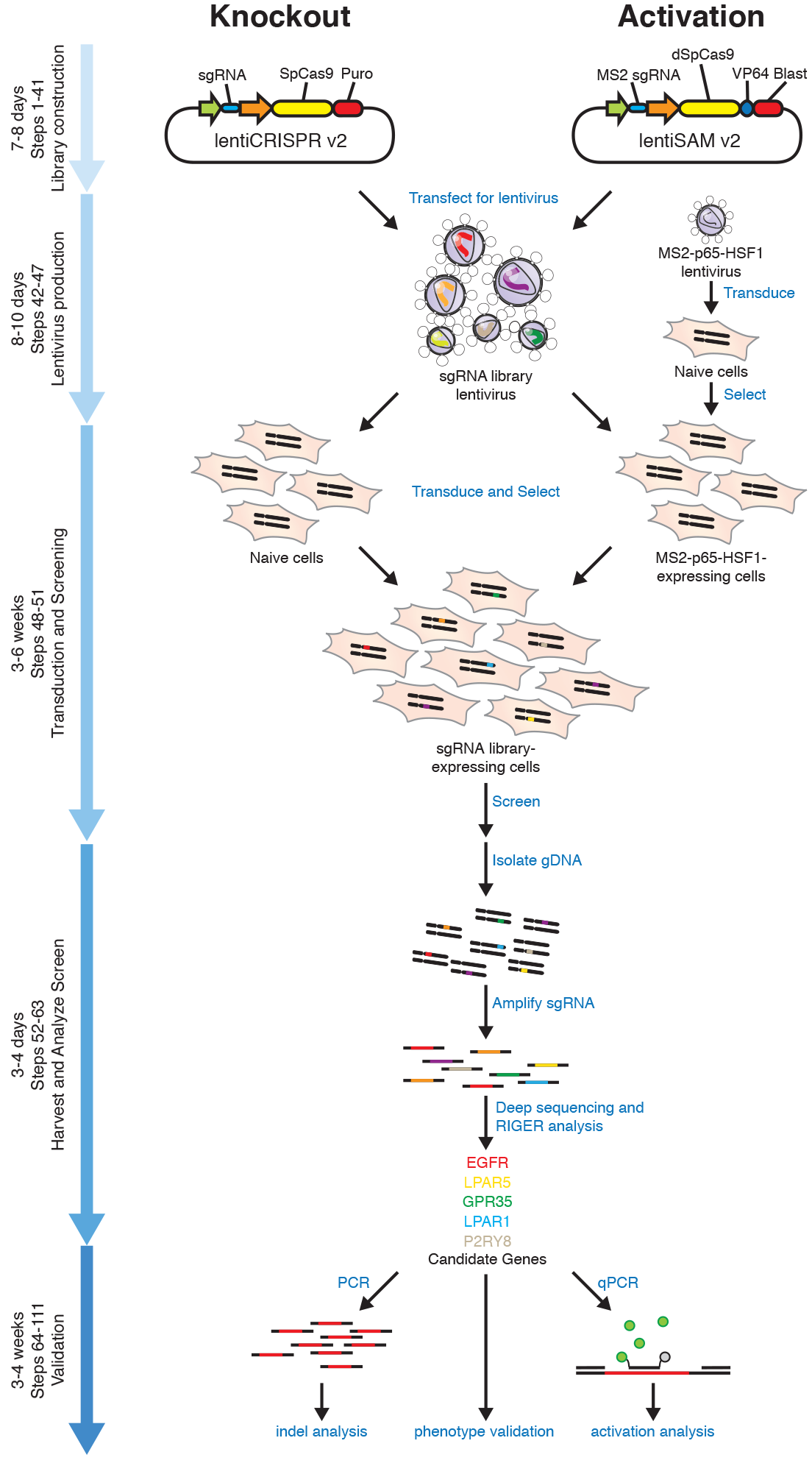
Timeline and overview of experiments. Genome-scale Cas9 knockout and transcriptional activation screens begin with the construction of a plasmid library encoding the effector protein and sgRNAs. These plasmid libraries are packaged into lentivirus and then transduced into the cell type of interest to generate stably expressing lines for the screen, along with an accessory transcriptional activator complex (MS2-p65-HSF1) lentivirus for the case of activation screening. A selection pressure is applied depending on the nature of the screen and at given timepoints, genomic DNA is harvested. The sgRNA regions are amplified from genomic DNA and then analyzed by next generation sequencing followed by statistical analyses (e.g. RIGER) to identify candidate genes. Candidate genes are then validated by various forms of analysis, including testing individual sgRNAs for the screening phenotype, indel formation by targeted sequencing, or transcript upregulation by qPCR.

**Table 1.**
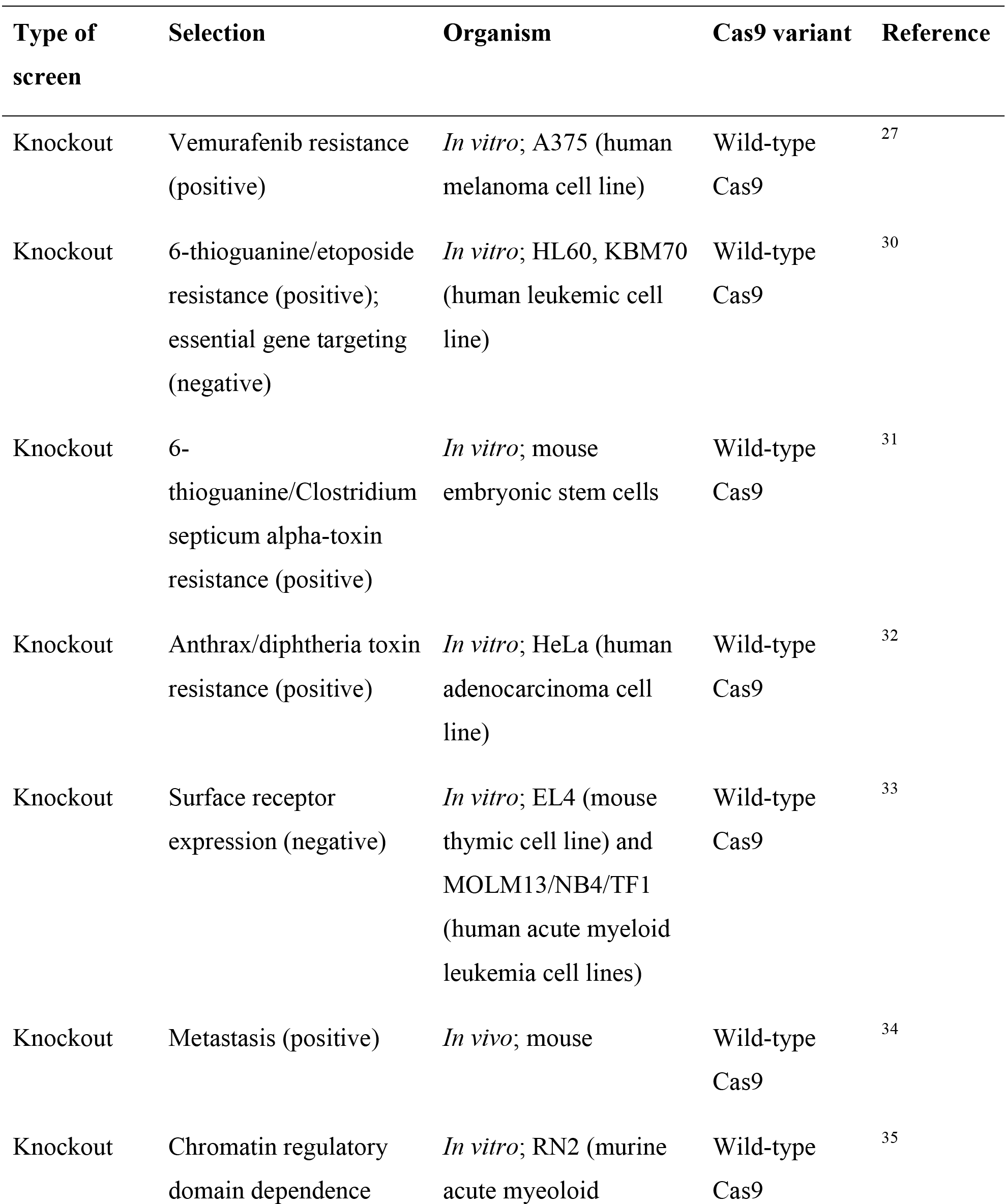
Previously published screens using Cas9.

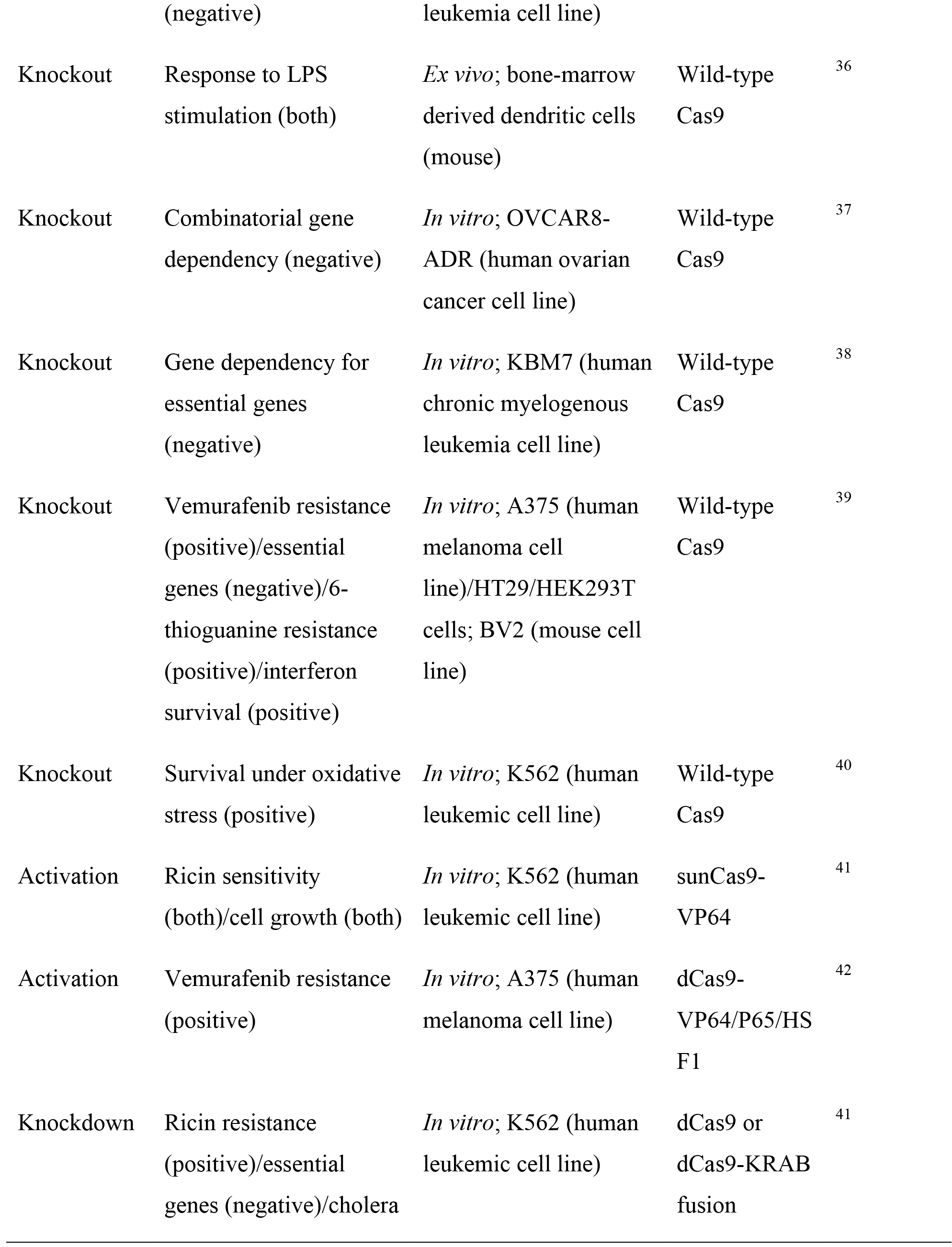

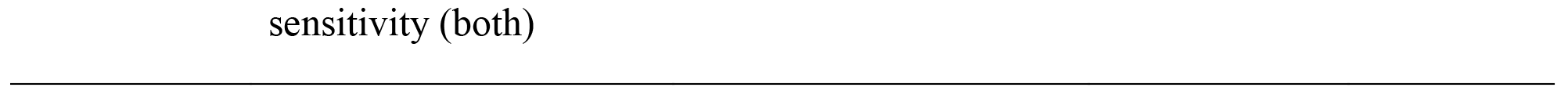

### Experimental Design

**Design and selection of the sgRNA library**. Although each sgRNA library is computationally designed for a specific purpose, the basic design process is consistent across libraries. First, the genomic regions of interest for targeting the sgRNA library are identified based on known sgRNA targeting rules (e.g. 5’ conserved exons for gene knockout, upstream or downstream of the transcriptional start site for transcriptional activation or repression respectively). Second, all possible sgRNA targets with the Cas9 ortholog-specific protospacer adjacent motif (PAM) are identified and selected based on four criteria: (i) minimization of off-target activity, (ii) maximization of on-target activity, (iii) avoidance of homopolymer stretches (e.g. AAAA, GGGG) and (iv) GC content. Recent work has begun to elucidate the features that govern sgRNA specificity and efficiency^33, 39^. Although specificity and efficiency will likely vary across experimental settings, false positive sgRNAs in screens can still be mitigated by including redundant sgRNAs in the library and requiring multiple distinct sgRNAs targeting the same gene to display the same phenotype when identifying screening hits. Once the targeting sgRNAs have been chosen, additional non-targeting guides that do not target the genome should be included as negative controls.

We provide several genome-scale libraries for knockout and activation screening through Addgene. For knockout screening, the GeCKO v2 libraries target the 5’ conserved coding exons of 19,050 human or 20,611 mouse coding genes with 6 sgRNAs per gene (Fig. 3)^54^. In addition to targeting coding genes, the GeCKO v2 libraries also target 1,864 human miRNAs or 1,175 mouse miRNAs with 4 sgRNAs per miRNA. Each species-specific library contains 1,000 non-targeting control sgRNAs. The GeCKO library is available in a 1 vector (lentiCRISPR v2) or 2 vector (lentiCas9-Blast and lentiGuide-Puro) format. For activation screening, the SAM libraries target the 200bp region upstream of the transcriptional start site of 23,430 human or 23,439 mouse RefSeq coding isoforms with 3 sgRNAs per isoform (Fig. 3)^42^. The library has to be combined with additional SAM effectors in a 2 vector (lentiSAM v2 and MS2-P65-HSF1) or 3 vector (dCas9-VP64, sgRNA(MS2), and MS2-P65-HSF1) format. Both GeCKO v2 and SAM libraries prioritize sgRNAs with minimal off-target activity.

**Figure 3.**
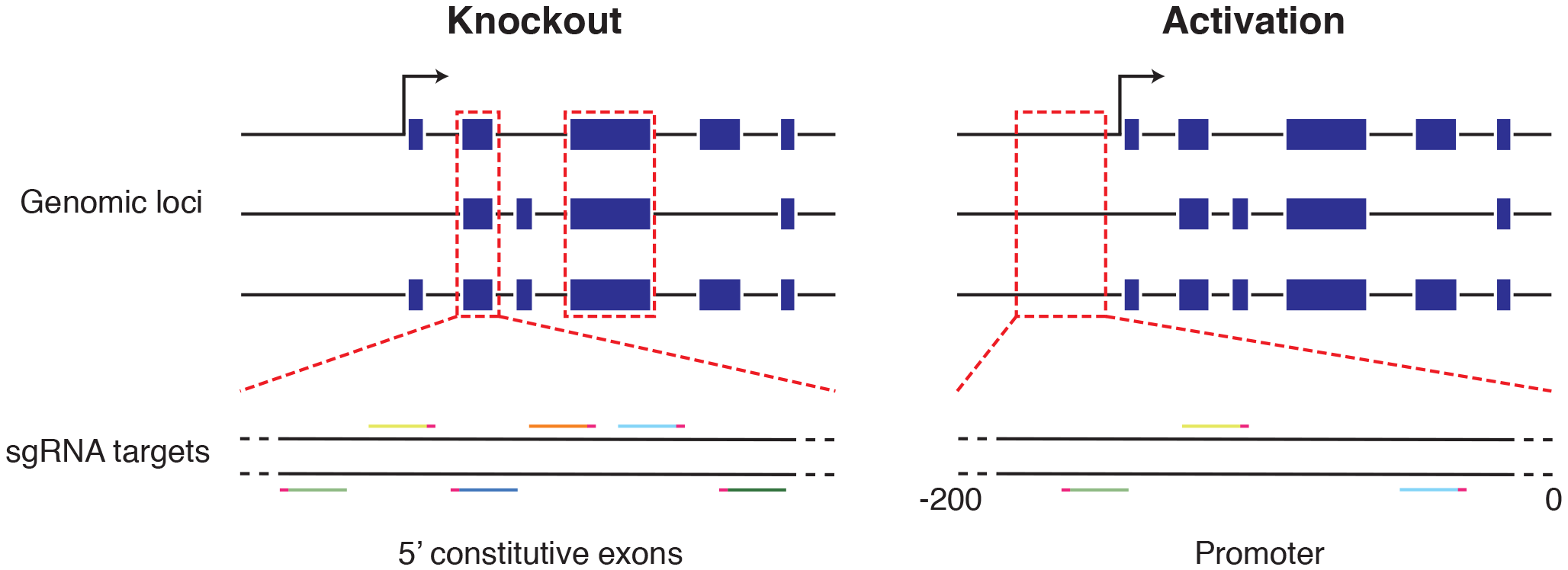
SgRNA library design for genome-scale knockout or activation screens. For knockout screening, the GeCKO v2 libraries target the 5’ conserved coding exons of 19,050 human or 20,611 mouse coding genes with 6 sgRNAs per gene. For activation screening, the SAM libraries target the 200bp region upstream of the transcriptional start site of 23,430 human or 23,439 mouse RefSeq coding isoforms with 3 sgRNAs per isoform. Both libraries select sgRNAs with minimal off-target activity.

In cases when a subset of genes is known to be involved in the screening phenotype and/or when the cell number is limited, one can consider performing a targeted screen that captures a subset of the genes in the genome-scale screens provided. We have included a python script for isolating the sgRNA target sequences corresponding to the genes in the targeted screen and adding flanking sequences for cloning. Additionally, one can consider adapting the sgRNA library plasmid backbone to the needs of the screen.
For instance, when screening *in vivo* in complex tissues, one can use a cell-type specific promoter to ensure that only the cell type of interest is perturbed. To select for successful transduction by FACS, one can replace the antibiotic selection marker with a fluorescent marker. For these situations, we provide a protocol for cloning a custom sgRNA library.

**Approaches for sgRNA library construction and delivery**. Throughout the sgRNA library cloning and amplification process, it is important to minimize any potential bias that may affect screening results. For example, the number of PCR cycles in the initial amplification of the pooled oligo library synthesis should be limited to prevent introducing bias during amplification. Scale each step of the cloning procedure provided according to the size of the library to reduce loss of sgRNA representation. After sgRNA library transformation, limit the growth time to avoid intercolony competition which can result in plasmid amplification bias. We provide a protocol and accompanying python script for assessing sgRNA library distribution by next-generation sequencing (NGS) prior to screening.

Depending on the desired application, the sgRNA library can be delivered with lentivirus, retrovirus, or adeno-associated virus (AAV). Lentivirus and retrovirus integrate into the genome, whereas AAV does not integrate and thus for screening, AAV delivery is limited to non-dividing cells. In contrast, retrovirus only transduces dividing cells. In addition, AAV has a smaller insert size capacity compared to lentivirus and retrovirus. As a result, to date most of the screens have relied on lentiviral delivery and we have provided two methods for lentivirus production and transduction.

**Screening**. Since the parameters for each screen differs according to the screening phenotype, in lieu of providing a protocol for screening we have outlined general considerations for setting the relevant screening parameters as well as technical advice for carrying out a screen in **Box 2**. Additional *in vivo* screening considerations are described in **Box 3**. We also provide guidelines for saturated mutagenesis screening design and analysis in **Box 4**.

**Analysis of screening results**. For examples of anticipated results, we provide data from genome-scale knockout and transcriptional activation screening for genes that confer BRAF inhibitor vemurafenib (PLX) resistance in a BRAF^V600E^ (A375) cell line^27, 42^. As a result of the screening selection pressure, at the end of the screen the sgRNA library distribution in the experimental condition should be significantly skewed compared to the baseline and control conditions, with some sgRNAs enriched and others depleted (**Fig. 4a,b**). The sgRNA representation, which is measured by NGS, in the experimental relative to the control condition determines the enrichment or depletion of the sgRNA. Then, depending on the type of screen (positive, negative, or marker gene selection), the enrichment or depletion of sgRNAs will be used to identify candidate genes that confer the screening phenotype.

**Figure 4.**
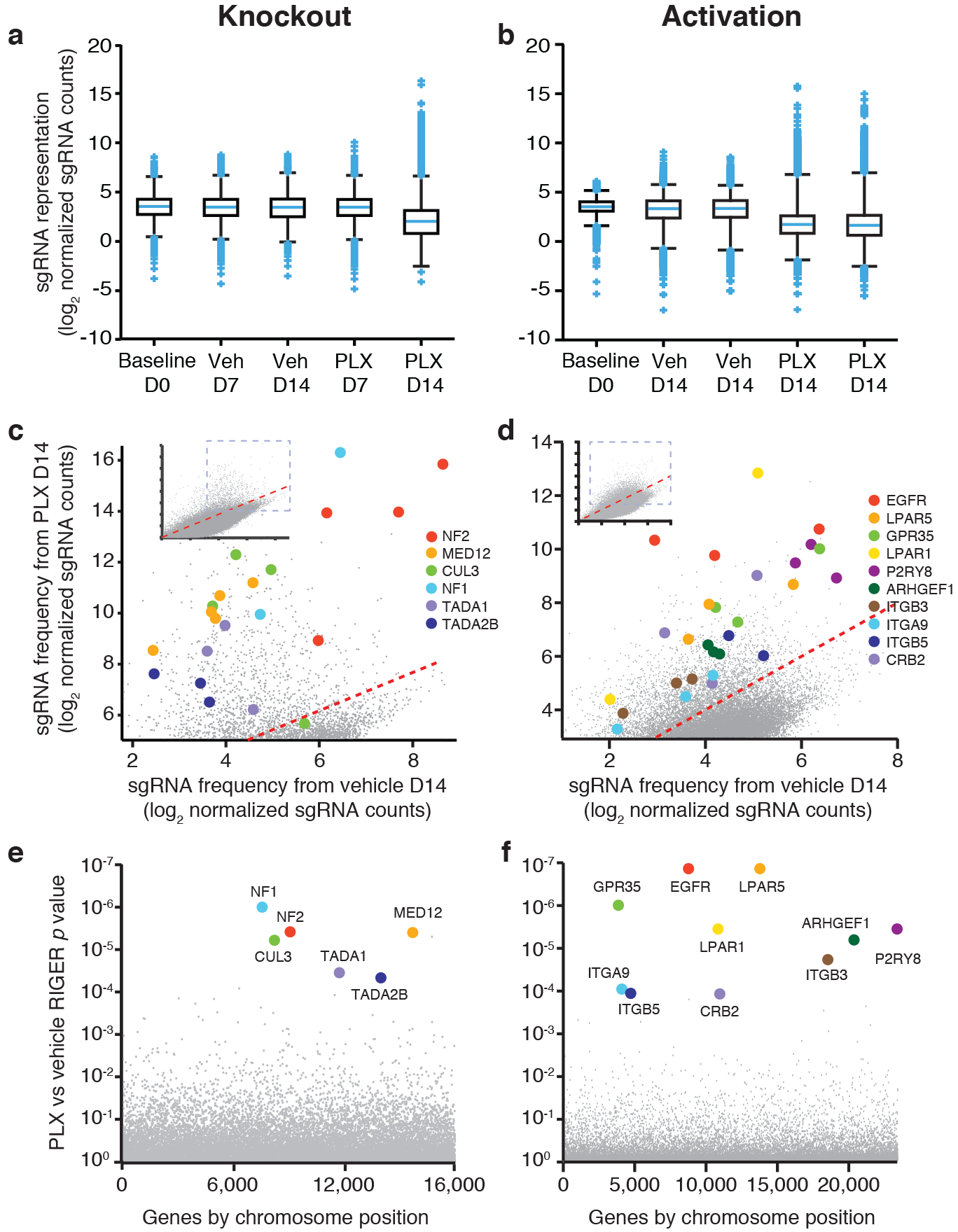
Anticipated results for genome-scale knockout and activation screens. Genome wide knockout and activation screens are performed to identify drivers of resistance to the BRAF inhibitor vemurafenib (PLX) in a BRAF^V600E^ (A375) melanoma cell line. A significant number of guides are seen enriched and depleted in the PLX day 14 condition, revealing depletion of guides essential for cell growth and enrichment of guides that promote resistance to BRAF inhibitor (**a,b**). RIGER identification of candidate enriched genes from the screens are highlighted. Each gene has multiple sgRNAs that are enriched. Many of these genes are known tumor suppressors or oncogenes that play a role in PLX4720 resistance (**c,d**). The top hits of the screen are seen as distributed across the genome, revealing the necessity of genome-scale screens for identifying drivers of resistance. RIGER p-values for candidate enriched genes are significantly lower (**e,f**).

Screening analysis methods such as RNAi gene enrichment ranking (RIGER) and redundant siRNA activity (RSA) typically select candidate genes with multiple enriched or depleted sgRNAs to reduce the possibility that the observed change in sgRNA distribution was due to off-target activity of a single sgRNA^55, 56^. RIGER ranks sgRNAs according to their enrichment or depletion and for each gene, examines the positions of the sgRNAs targeting that gene in the ranked sgRNA list^55^. The algorithm then assesses whether the set of positions is biased towards the top of the list using a Kolmogorov-Smirnov statistic and calculates an enrichment score and gene ranking based on a permutation test. RSA is similar to RIGER, except that it assigns statistical significance based on an iterative hypergeometric distribution formula^56^. In this protocol we describe in detail how to identify candidate genes using RIGER.

Each candidate gene identified from the screening analysis should have multiple significantly enriched or depleted sgRNAs in the experimental condition relative to the control (**Fig. 4c,d**). The RIGER *P* values of the candidate genes should also be significantly lower than the rest of the genes (**Fig. 4e,f**).

**Validation of candidate genes**. Given that the screening process can be noisy and the analysis produces a ranked list of candidate genes, it is necessary to verify that perturbation of the identified candidate genes confers the phenotype of interest. For validation, each of the sgRNAs that target the candidate gene can be individually cloned into the plasmid backbone of the sgRNA library and validated for the screening phenotype. In addition, the perturbation induced by each sgRNA, indel rate and transcriptional activation for knockout and activation screening respectively, will be quantified to establish a phenotype-to-genotype relationship.

Indel rates can be detected either by the SURVEYOR nuclease assay or by NGS. Compared to SURVEYOR, which we have described previously^57^, NGS is more suitable for sampling a large number of sgRNA target sites and therefore described here. For measuring indel rates, it is important to design primers situated at least 50 bp from the target cleavage site to allow for the detection of longer indels. Our protocol for targeted NGS outlines a two-step PCR in which the first step uses custom primers to amplify the genomic region of interest and the second step uses universal, barcoded primers for multiplex deep sequencing on the Illumina platform. Relative to the one-step PCR method recommended for preparing sgRNA libraries for NGS, the two-step PCR method is more versatile and less costly for assessing many different target sites because custom primers for each target site can be readily combined with different universal, barcoded primers.

After NGS, indel rates can be calculated by running the provided python script that implements two different algorithms. The first aligns reads using the Ratcliff-Obershelp algorithm and then finds regions of insertion or deletion from this alignment^58^. The second method, adapted from the Geneious aligner scans k-mers across the read and maps the alignment to detect indels^59^. In practice, Ratcliff-Obershelp alignment algorithm is more accurate, while k-mer based alignment algorithm is faster. These indel rates are then adjusted to account for background indel rates via a maximum likelihood estimation (MLE) correction^25^. The MLE correction models the observed indel rate as a combination of the true indel rate resulting from Cas9 cleavage and a separately measured background indel rate. The true indel rate is that which maximizes the probability of the observed read counts under the assumption that they obey a binomial distribution with the background rate.

Measurement of transcriptional activation usually entails isolation of RNA, reverse transcription of the RNA to cDNA, and quantitative PCR (qPCR). Various different methods have been described for each step of the process. In this protocol, we provide a method for reverse transcription followed by qPCR that is rapid, high-throughput, and cost-effective and thus ideal for quantifying fold upregulation for validation. Our method involves direct lysis of cells grown on a 96-well plate followed by reverse transcription and TaqMan qPCR. TaqMan-based detection is more specific and reproducible than SYBR-based detection because it relies on a fluorogenic probe specific to the target gene, whereas SYBR depends on a dsDNA-binding dye. TaqMan also allows for multiplexing with control probes that measure housekeeping gene expression as a proxy for total RNA concentration.

After validation of the screening phenotype and perturbation, we recommend verifying down-or up-regulation of protein expression for knockout or transcriptional activation screening respectively. Immunohistochemistry and Western blot are two of the most common methods for verifying protein expression. Immunohistochemistry requires fixing the validation cell lines and detecting the target protein using a specific antibody, whereas Western blot involves harvesting protein and separating by electrophoresis before staining with the specific antibody. Although immunohistochemistry provides additional information on protein localization, it often requires a more specific antibody than Western blot because proteins are not separated by size. Thus, Western blot is preferable for verifying protein expression of candidate genes.

## MATERIALS

### REAGENTS

#### SgRNA libraries and backbones

- lentiCRISPR v2 (Addgene, cat. no. 52961)
- lentiCas9-Blast (Addgene, cat. no. 52962)
- lentiGuide-Puro (Addgene, cat. no. 52963)
- lenti dCAS-VP64_Blast (Addgene, cat. no. 61425)
- lenti MS2-P65-HSF1_Hygro (Addgene, cat. no. 61426)
- lenti sgRNA(MS2)_zeo backbone (human; Addgene, cat. no. 61427)
- lenti sgRNA(MS2)_puro backbone (human; Addgene, cat. no. 73795)
- lenti sgRNA(MS2)_puro optimized backbone (mouse; Addgene, cat. no. 73797)
- lentiSAMv2 backbone (human; Addgene, cat. no. 75112)
- Human GeCKO v2 Library, 1 plasmid system (Addgene, cat. no. 1000000048)
- Human GeCKO v2 Library, 2 plasmid system (Addgene, cat. no. 1000000049)
- Mouse GeCKO v2 Library, 1 plasmid system (Addgene, cat. no. 1000000052)
- Mouse GeCKO v2 Library, 2 plasmid system (Addgene, cat. no. 1000000053)
- Human SAM Library, Zeo, 3 plasmid system (Addgene, cat. no. 1000000057)
- Human SAM Library, Puro, 3 plasmid system (Addgene, cat. no. 1000000074)
- Mouse SAM Library, Puro optimized, 3 plasmid system (Addgene, cat. no. 1000000075)
- Human SAM library, lentiSAMv2, 2 plasmid system (Addgene, cat. no. 75117)

#### Custom sgRNA library cloning

- Pooled oligo library (Twist Bioscience or CustomArray)
- PCR primers for amplifying oligo library for cloning are listed in **Table 2**. Primers longer than 60bp can be ordered as 4-nmol ultramers (Integrated DNA technologies)
- NEBNext High Fidelity PCR Master Mix, 2× (New England BioLabs, cat. no. M0541L)
- UltraPure DNase/RNase-free distilled water (Thermo Fisher, cat. no. 10977023)
- QIAquick PCR Purification Kit (Qiagen, cat. no. 28104)
- QIAquick gel extraction kit (Qiagen, cat. no. 28704)
- UltraPure TBE buffer, 10x (Thermo Fisher, cat. no. 15581028)
- SeaKem LE agarose (Lonza, cat. no. 50004)
- SYBR Safe DNA stain, 10,000x (Thermo Fisher, cat. no. S33102)
- 1-kb Plus DNA ladder (Thermo Fisher, cat. no. 10787018)
- 50bp DNA ladder (Thermo Fisher, cat. no. 10416014)
- TrackIt Cyan/Orange Loading Buffer (Thermo Fisher, cat. no. 10482028)
- FastDigest Esp3I (BsmBI; Thermo Fisher, cat. no. FD0454)
- FastAP Thermosensitive Alkaline Phosphatase (Thermo Fisher, cat. no. EF0651)
- DTT, Molecular Grade (Promega, cat. no. P1171)
- Gibson Assembly Master Mix (New England BioLabs, cat. no. E2611L)
- GlycoBlue Coprecipitant (Thermo Fisher, cat. no. AM9515)
- 2-Propanol (Sigma-Aldrich, cat. no. I9516-25ML)
- Sodium chloride solution (Sigma-Aldrich, cat. no. 71386-1L)
- Ethyl alcohol, Pure (Sigma-Aldrich, cat. no. 459844-500ML)
- Tris-EDTA buffer solution (Sigma-Aldrich, cat. no. 93283-100ML)

**Table 2.**
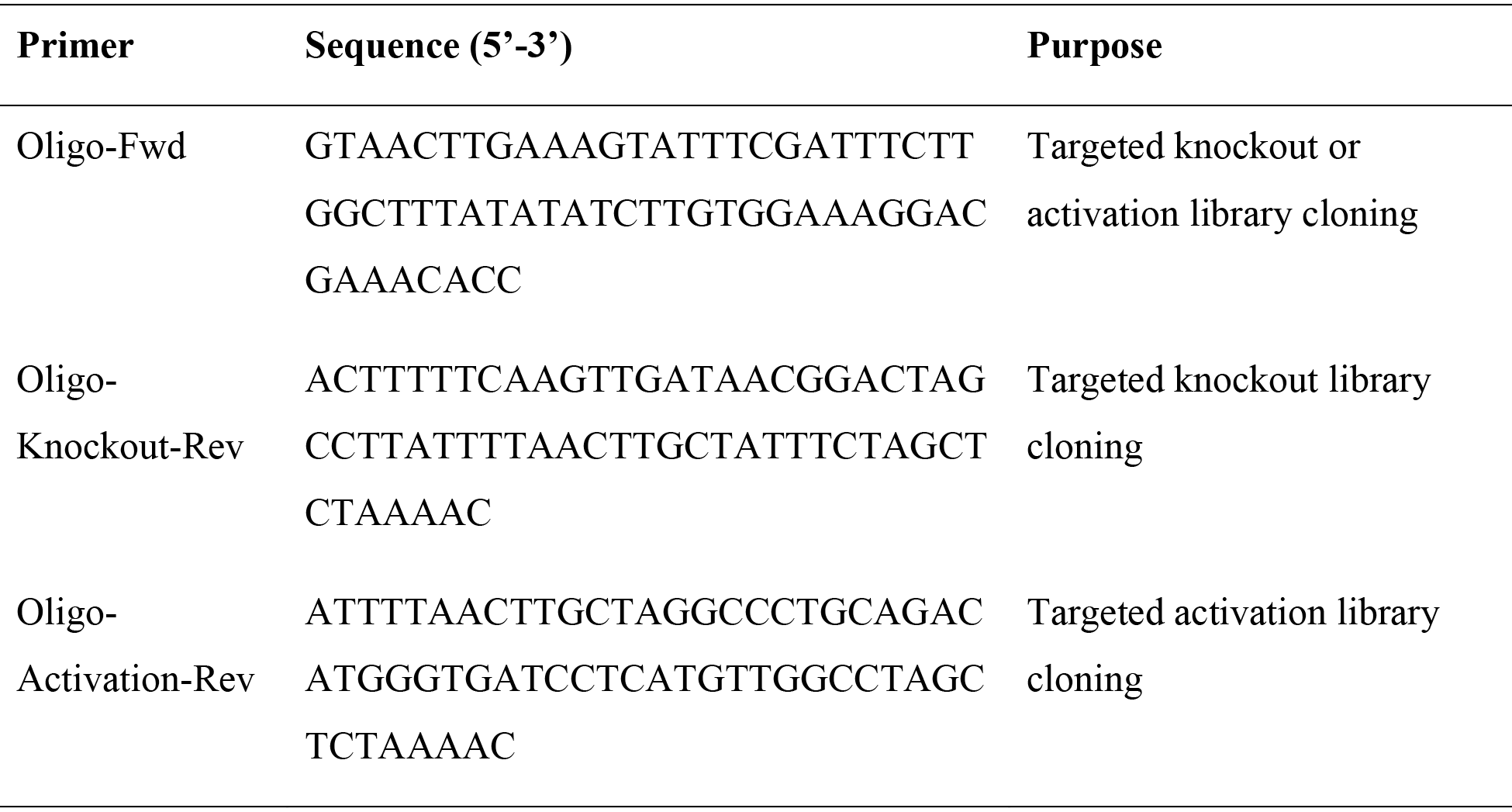
Primers for amplifying the sgRNA oligo library.

#### SgRNA plasmid amplification

- LB Agar, Ready-Made Powder (Affymetrix, cat. no. 75851)
- LB Broth, Ready-Made Powder (Affymetrix, cat. no. 75852)
- Ampicillin, 100 mg ml^-1^, sterile filtered (Sigma-Aldrich, cat. no. A5354)
- Endura ElectroCompetent Cells (Lucigen, cat. no. 60242-1)
- NucleoBond Xtra Maxi EF (Macherey-Nagel, cat. no. 740424.10)
- One Shot Stbl3 chemically competent *E. coli* (Thermo Fisher, cat. no. C737303)
- SOC outgrowth medium (New England BioLabs, cat. no. B9020S)
- NucleoBond Xtra Midi EF (Macherey-Nagel, cat. no. 740420.50)

#### Next-generation sequencing

- Primers for amplifying the library for NGS are listed in **Table 3**. Primers for amplifying indels for NGS are listed in **Table 5**. Primers longer than 60bp can be ordered as 4 nmol ultramers (Integrated DNA technologies)
- NEBNext High Fidelity PCR Master Mix, 2× (New England BioLabs, cat. no. M0541L)
- KAPA HiFi, HotStart ReadyMix, 2× (Kapa Biosystems, cat. no. KK2602)
- UltraPure DNase/RNase-free distilled water (Thermo Fisher, cat. no. 10977023)
- QIAquick PCR Purification Kit (Qiagen, cat. no. 28104)
- QIAquick gel extraction kit (Qiagen, cat. no. 28704)
- UltraPure TBE buffer, 10× (Thermo Fisher, cat. no. 15581028)
- SeaKem LE agarose (Lonza, cat. no. 50004)
- SYBR Safe DNA stain, 10,000x (Thermo Fisher, cat. no. S33102)
- 1-kb Plus DNA ladder (Thermo Fisher, cat. no. 10787018)
- TrackIt CyanOrange loading buffer (Thermo Fisher, cat. no. 10482028)
- Qubit dsDNA HS Assay Kit (Thermo Fisher, cat. no. Q32851)
- NextSeq 500/550 High Output Kit v2 (150 cycle; Illumina, cat. no. FC-404-2002)
- MiSeq Reagent Kit v3 (150 cycle; Illumina, cat. no. MS-102-3001)
- MiSeq Reagent Kit v2 (300 cycle; Illumina, cat. no. MS-102-2002)
- PhiX Control Kit v3 (Illumina, cat. no. FC-110-3001)
- Sodium hydroxide solution, 10 N (Sigma-Aldrich, cat. no. 72068-100ML)
- Tris, pH 7.0 (Thermo Fisher, cat. no. AM9850G)

**Table 3.**
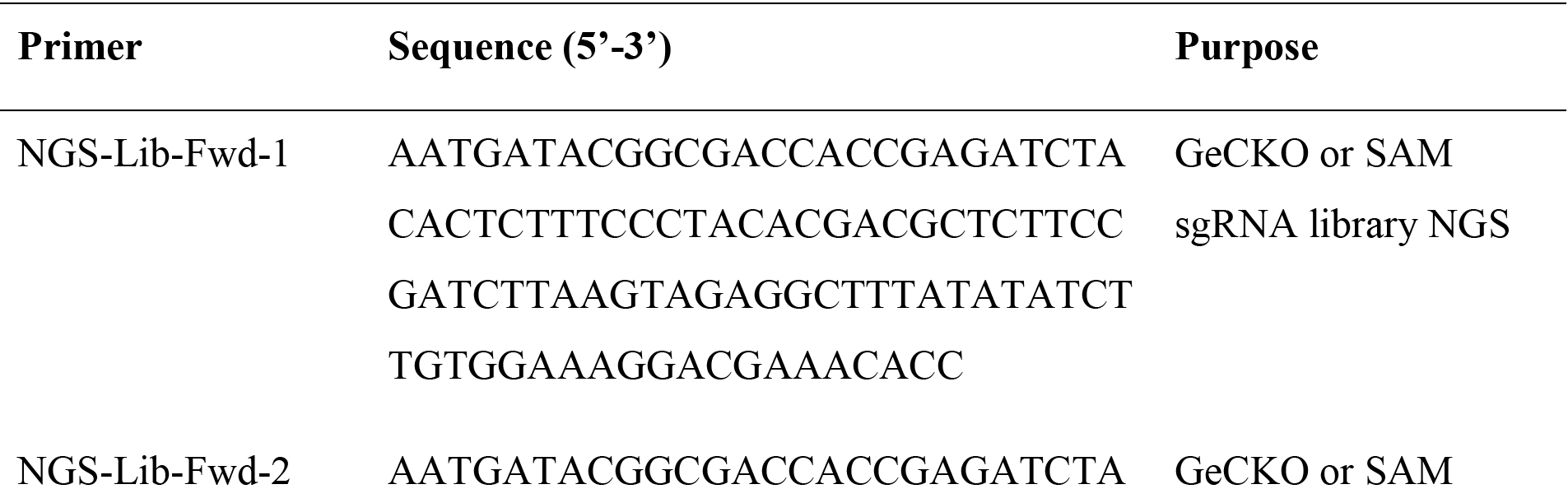
Primer sequences for amplifying sgRNA library and NGS. The NGS-Lib-Fwd primers contain 1-10bp staggered nucleotides designed to increase the diversity of the NGS library, and the NGS-Lib-Rev primers provide unique barcodes for distinguishing different sgRNA libraries (i.e. from different screening conditions) in a pooled sequencing run. Since the sgRNA backbone is different between the GeCKO and SAM libraries, separate NGS-Lib-KO-Rev and NGS-Lib-SAM-Rev primers have been provided for each library.

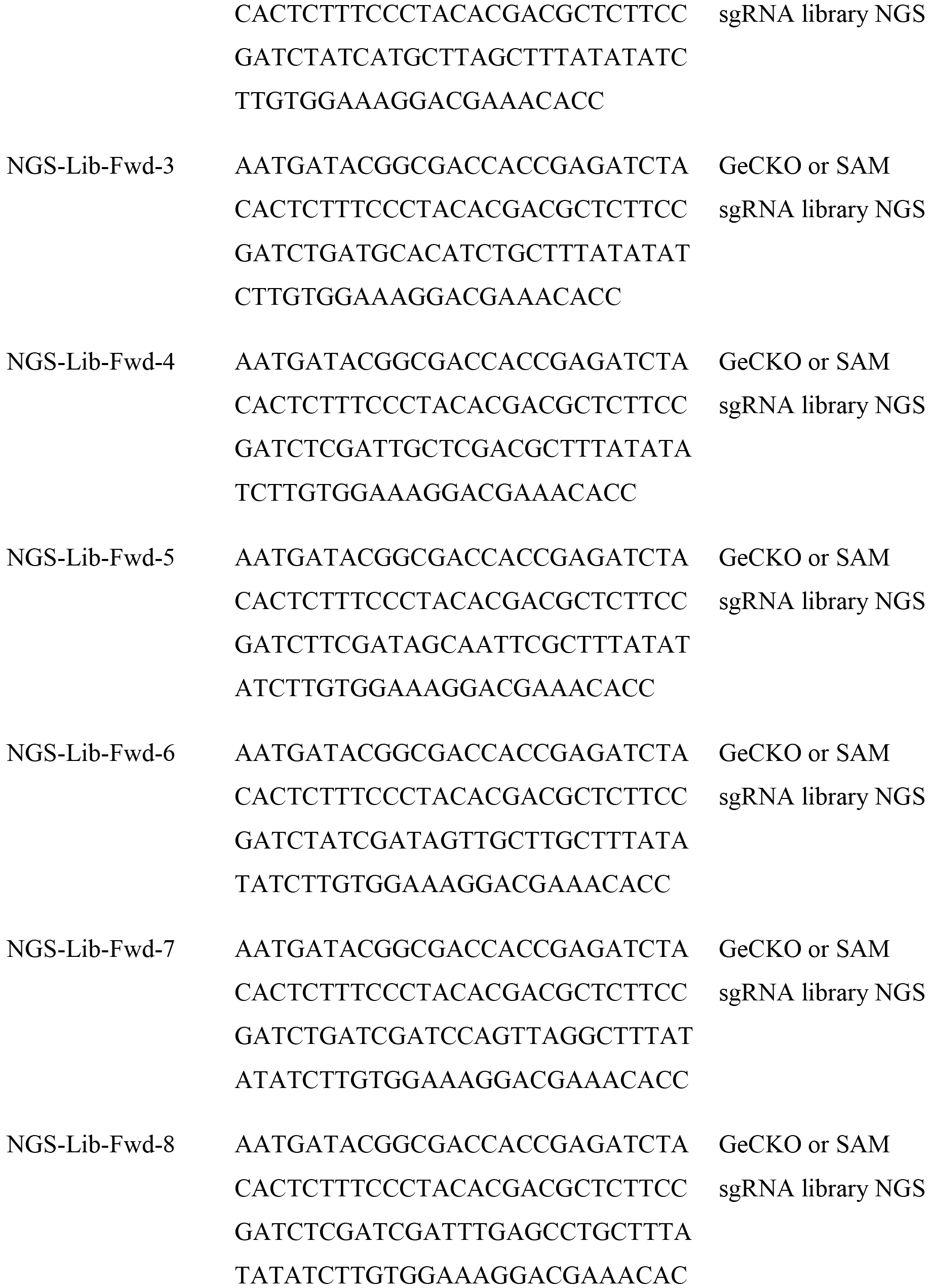

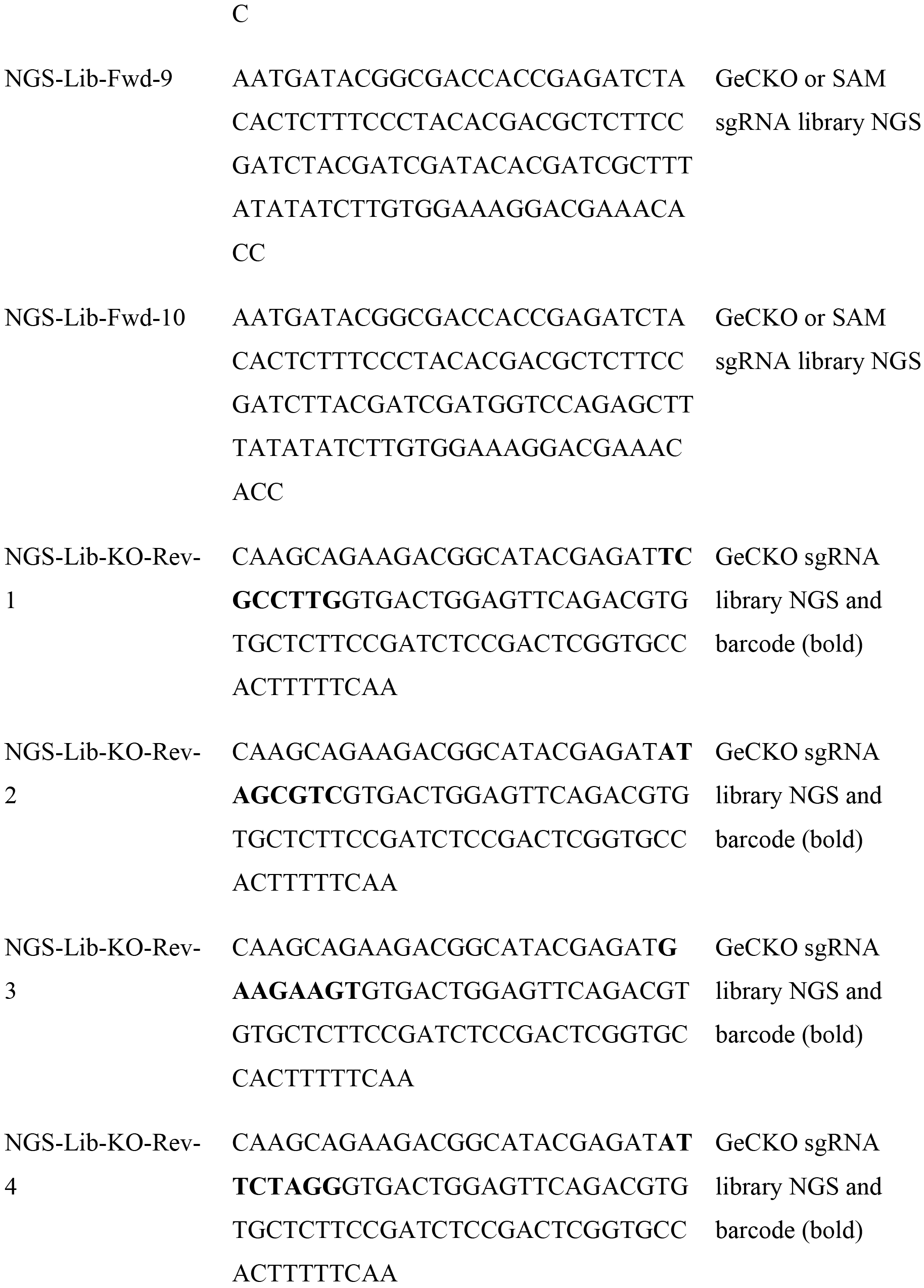

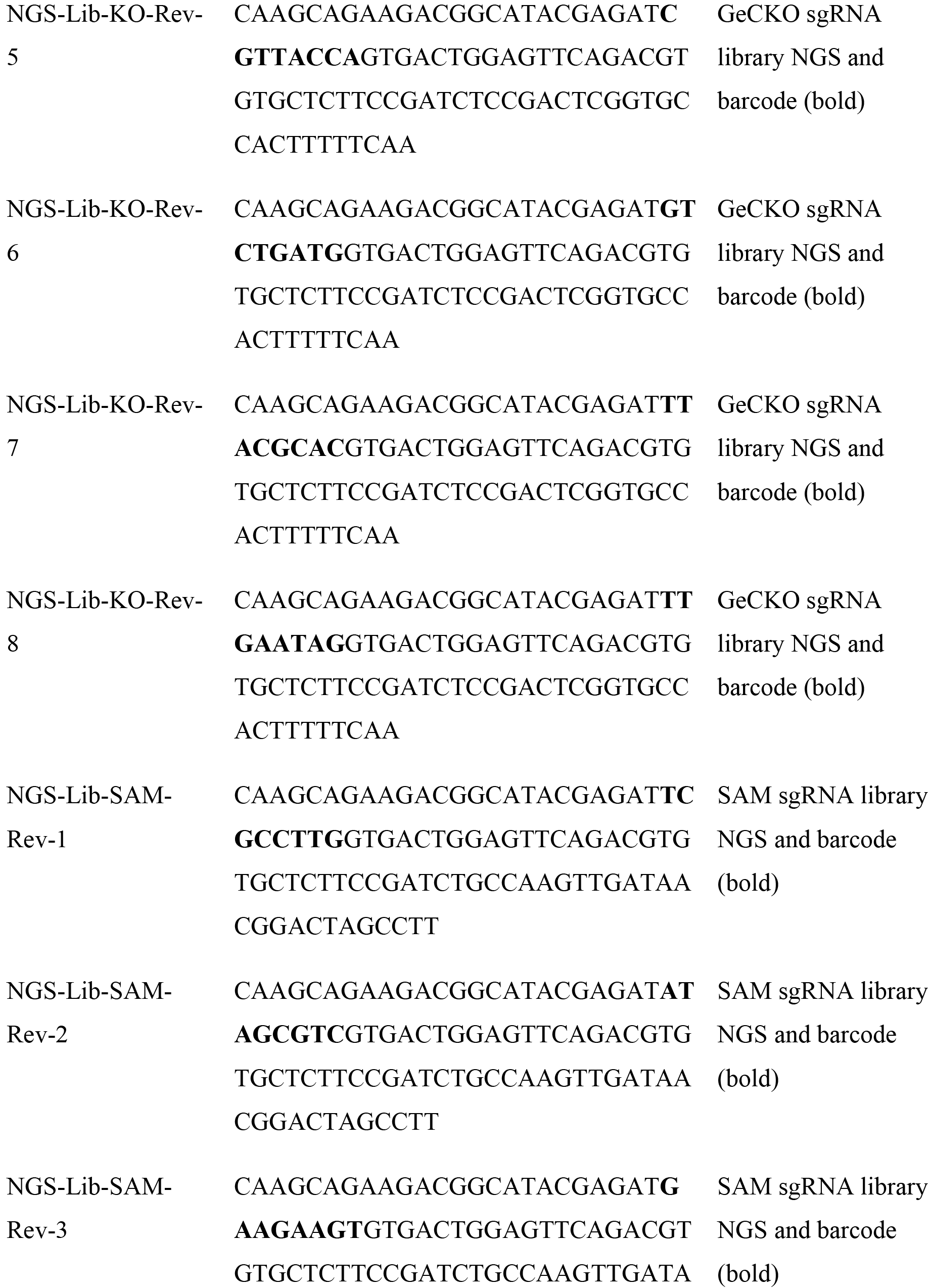

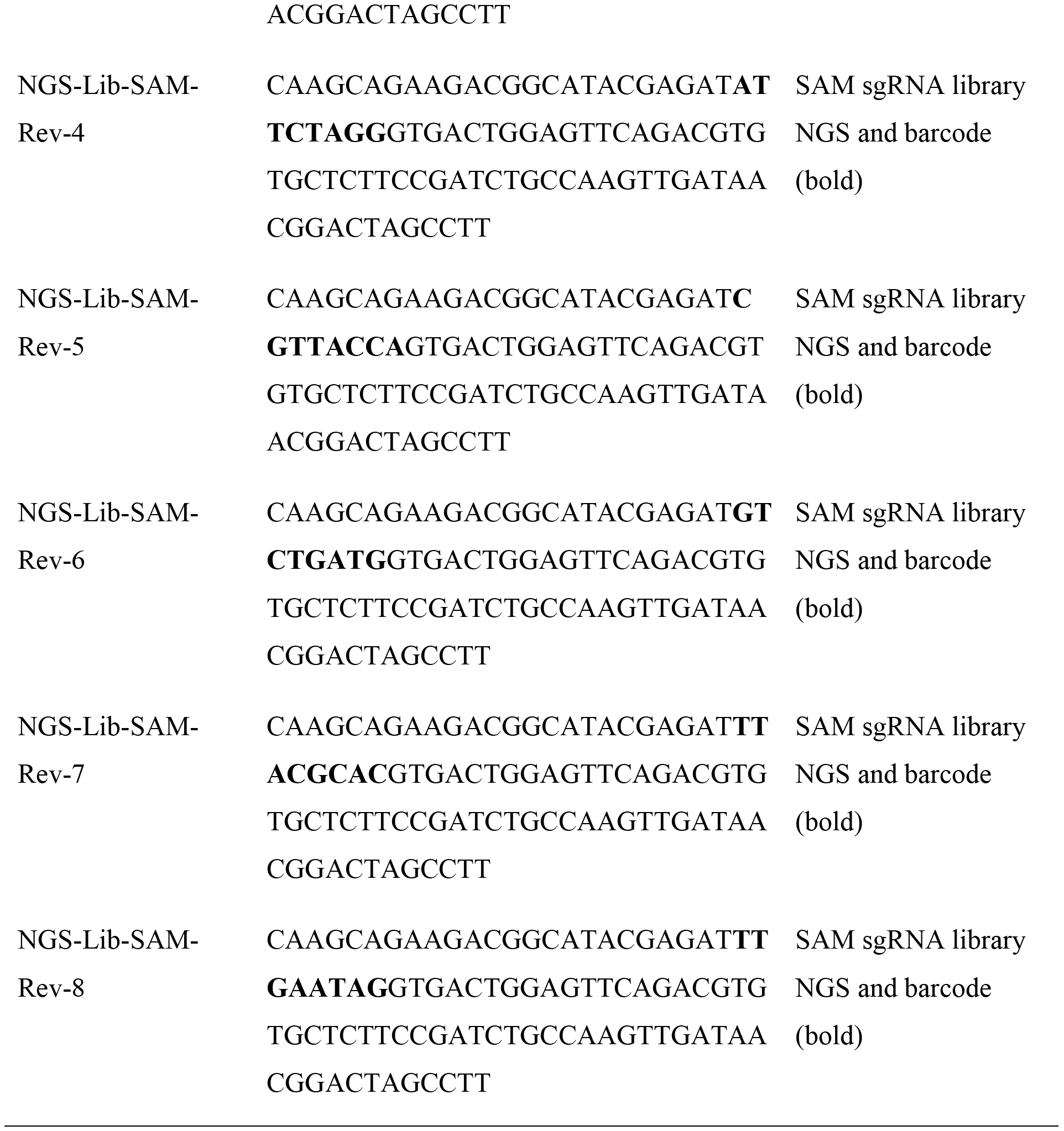

#### Mammalian cell culture

- HEK293FT (Thermo Fisher, cat. no. R70007)
- DMEM, high glucose, GlutaMAX Supplement, pyruvate (Thermo Fisher, cat. no. 10569010)
- Penicillin-streptomycin, 100× (Thermo Fisher, cat. no. 15140122)
- Fetal bovine serum, premium grade (VWR, cat. no. 97068-085)
- TrypLE Express, no phenol red (Thermo Fisher, cat. no. 12604021)
- HUES 66 cell line (Harvard Stem Cell Science)
- Geltrex LDEV-free reduced growth factor basement membrane matrix (Thermo Fisher, cat. no. A1413202)
- mTeSR1 medium (Stemcell Technologies, cat. no. 05850)
- Normocin (InvivoGen, cat. no. ant-nr-1)
- Rho-associated protein kinase (ROCK) inhibitor (Y-27632; Millipore, cat. no. SCM075)
- Accutase (Stemcell Technologies, cat. no. 07920)
- Dulbecco's PBS (DPBS; Thermo Fisher, cat. no. 14190250)

#### Lentivirus production and titer

- Opti-MEM I reduced serum medium (Thermo Fisher, cat. no. 31985062)
- pMD2.G (Addgene, cat. no. 12259)
- psPAX2 (Addgene, cat. no. 12260)
- pcDNA3-EGFP transfection control plasmid (Addgene, cat. no. 13031)
- Lipofectamine 2000 transfection reagent (Thermo Fisher, cat. no. 11668019)
- PLUS Reagent (Thermo Fisher, cat. no. 11514015)
- Polyethylenimine HCl MAX, Linear, Mw 40,000 (PEI Max; Polysciences, cat. no. 24765-1)
- Sodium hydroxide solution, 10 N (Sigma-Aldrich, cat. no. 72068-100ML)
- Polybrene (Hexadimethrine bromide; Sigma-Aldrich, cat. no. 107689-10G)
- Blasticidin S HCl (Thermo Fisher, cat. no. A1113903)
- Puromycin dihydrochloride (Thermo Fisher, cat. no. A1113803)
- Hygromycin B (Thermo Fisher, cat. no. 10687010)
- Zeocin (Thermo Fisher, cat. no. R25001)
- CellTiter-Glo Luminescent Cell Viability Assay (Promega, cat. no. G7571)

#### Screening and Validation

- Quick-gDNA MidiPrep (Zymo Research, cat. no. D3100)
- DNA Binding Buffer (Zymo Research, cat. no. D4004-1-L)
- DNA Wash Buffer (Zymo Research, cat. no. D4003-2-24)
- DNA Elution Buffer (Zymo Research, cat. no. D3004-4-4)
- Ethyl alcohol, Pure (Sigma-Aldrich, cat. no. 459844-500ML)
- Primers for cloning the validation sgRNAs are listed in **Table 4**. (Integrated DNA technologies)
- T4 polynucleotide kinase (New England BioLabs, cat. no. M0201S)
- T4 DNA ligase reaction buffer, 10× (New England BioLabs, cat. no. B0202S)
- UltraPure DNase/RNase-free distilled water (Thermo Fisher, cat. no. 10977023)
- T7 DNA ligase with 2× rapid ligation buffer (Enzymatics, cat. no. L6020L)
- BSA, Molecular Biology Grade (New England BioLabs, cat. no. B9000S)
- FastDigest Esp3I (BsmBI; Thermo Fisher, cat. no. FD0454)
- DTT, Molecular Grade (Promega, cat. no. P1171)
- QuickExtract DNA extraction solution (Epicentre, cat. no. QE09050)
- RNase AWAY (VWR, cat. no. 53225-514)
- Proteinase K (Sigma-Aldrich, cat. no. P2308-25MG)
- Tris, 1 M, pH 8.0 (Thermo Fisher, cat. no. AM9855G)
- Deoxyribonuclease I bovine (Sigma-Aldrich, cat. no. D2821-50KU)
- UltraPure 1 M Tris-HCI Buffer, pH 7.5 (Thermo Fisher, cat. no. 15567027)
- Calcium chloride solution (Sigma-Aldrich, cat. no. 21115-1ML)
- Glycerol (Sigma-Aldrich, cat. no. G5516-100ML)
- MgCl_2_, 1 M (Thermo Fisher, cat. no. AM9530G)
- Triton X-114 (Sigma-Aldrich, cat. no. X114-100ML)
- DTT, Molecular Grade (Promega, cat. no. P1171)
- Proteinase K Inhibitor (EMD Millipore, cat. no. 539470-10MG)
- Dimethyl sulfoxide (Sigma-Aldrich, cat. no. D8418-50ML)
- Ethylene glycol-bis(2-aminoethylether)-N,N,N',N'-tetraacetic acid (Sigma-Aldrich, cat. no. E3889-10G)
- Sodium hydroxide solution, 10 N (Sigma-Aldrich, cat. no. 72068-100ML)
- RevertAid RT Reverse Transcription kit (Thermo Fisher, cat. no. K1691)
- Oligo dT (TTTTTTTTTTTTTTTTTTTTNN; Integrated DNA Technologies)
- TaqMan target probes, FAM dye (Thermo Fisher)
- Taqman endogenous control probe, VIC dye (e.g. Human GAPD, GAPDH, Endogenous Control VIC^®^/MGB probe, primer limited; Thermo Fisher, cat. no. 4326317E)
- TaqMan Fast Advanced Master Mix, 2x (Thermo Fisher, cat. no. 4444557)

**Table 4.**
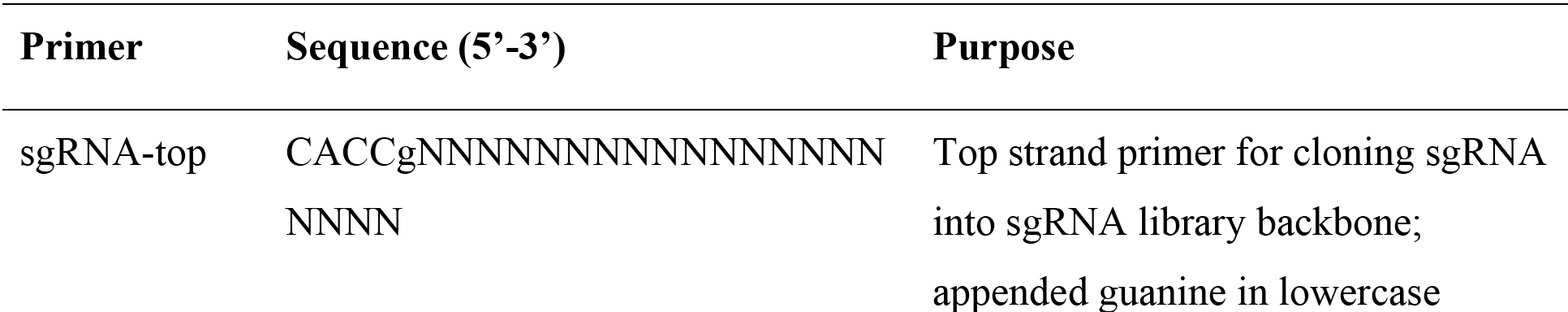
Primers for sgRNA cloning and validation.

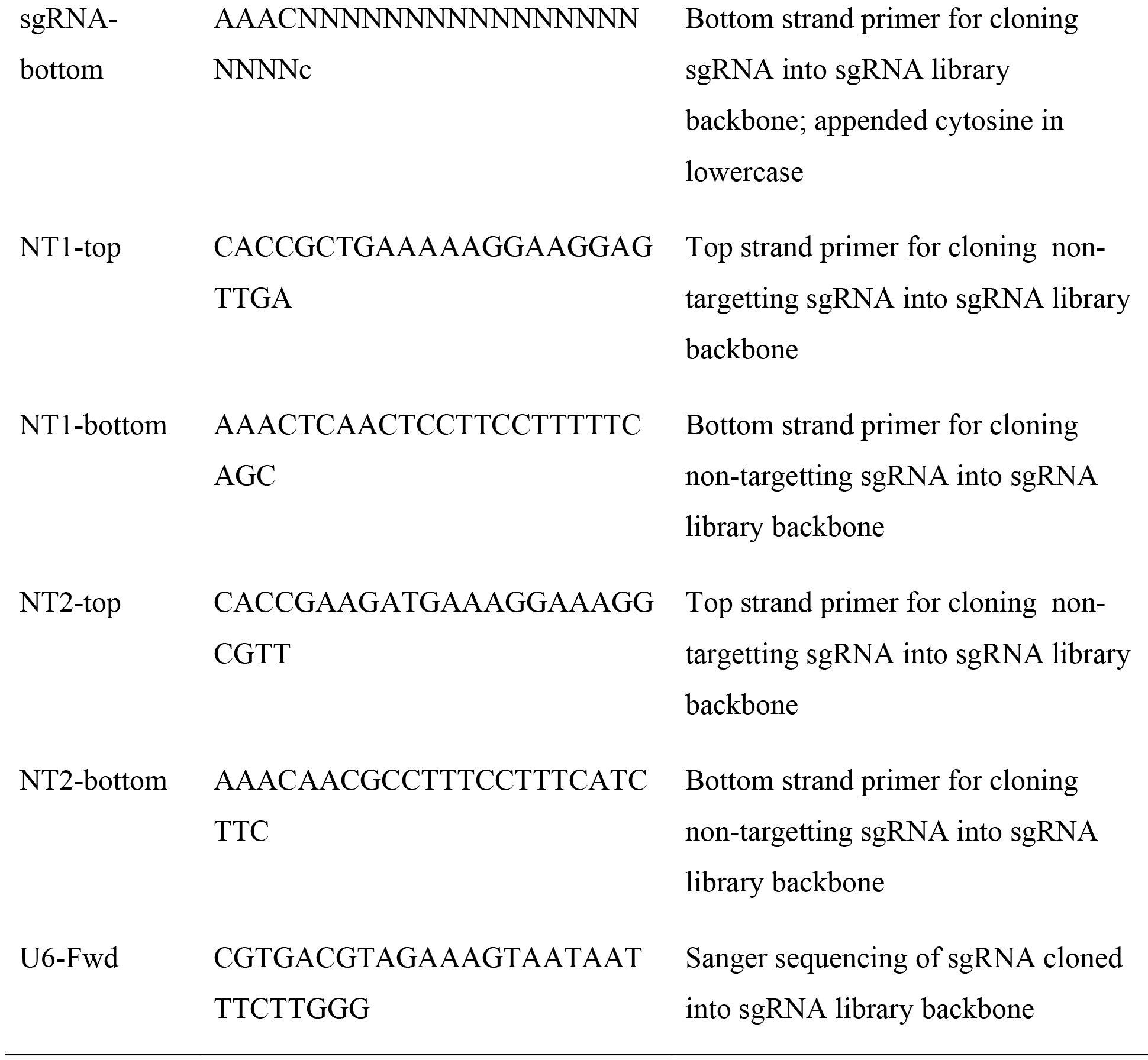

### EQUIPMENT

- Axygen 8-Strip PCR tubes (Fisher Scientific, cat. no. 14-222-250)
- Axygen PCR plates, 96 well (VWR, cat. no. PCR-96M2-HS-C)
- 384-well optical plate (e.g. Roche, LightCycler 480 Multiwell plate 384, cat. no. 5102430001)
- Axygen 1.5 ml Boil-Proof Microcentrifuge Tubes, (VWR, cat. no. 10011-702)
- Falcon tubes, polypropylene, 15 ml (Corning cat. no. 352097)
- Falcon tubes, polypropylene, 50 ml (Corning, cat. no. 352070)
- Filtered sterile pipette tips (e.g. Rainin)
- 100 mm × 15 mm Not TC-Treated Bacteriological Petri Dish (Corning, cat. no. 351029)
- 245mm Square BioAssay Dish without Handles, not TC-Treated Culture (Corning, cat. no. 431111)
- VWR Bacti Cell Spreaders (VWR, cat. no. 60828-688)
- AirPore Tape Sheets (Qiagen, cat. no. 19571)
- Nunc EasYFlask 25cm^2^, Filter Cap, 7 ml working volume (T25 flask; Thermo Scientific, cat. no. 156367)
- Nunc EasYFlask 75cm^2^, Filter Cap, 25 ml working volume, (T75 flask; Thermo Scientific, cat. no. 156499)
- Nunc EasYFlask 225 cm^2^, filter cap, 70 ml working volume (T225 flask; Thermo Scientific, cat. no. 159934)
- Stericup filter unit, 0.45 μM (Millipore, cat. no. SCHVU02RE)
- Syringe filter unit, 0.45 μM (Millipore, cat. no. SLHV013SL)
- Disposable Syringes with Luer-Lok Tip (Fisher Scientific, cat. no. 14-829-45)
- Falcon tissue culture plate, 6 wells (Corning, cat. no. 353224)
- Falcon tissue culture plate, 12 wells (Corning, cat. no. 353043)
- Falcon tissue culture dish, 100mm (Corning, cat. no. 353003)
- 96 well flat clear bottom black polystyrene TC-treated microplates (Corning, cat. no. 3904)
- BD BioCoat clear Poly-D-Lysine 96 well clear flat bottom TC-treated microplate (Corning, cat. no. 356461)
- Cellometer SD100 Counting Chambers (Nexcelom Bioscience, cat. no. CHT4-SD100-002)
- Zymo-Spin V with Reservoir (Zymo Research, cat. no. C1016-25)
- Collection Tubes, 2 ml (Zymo Research, cat. no. C1001-25)
- Amicon Ultra-15 Centrifugal Filter Unit with Ultracel-100 membrane (Millipore, cat. no. UFC910008)
- Thermocycler with programmable temperature stepping functionality, 96 well (e.g. Applied Biosystems Veriti, cat. no. 4375786)
- Real-time PCR system, 384 well (e.g. Roche Lightcycler 480 cat. no. 05015243001)
- Desktop microcentrifuges (e.g. Eppendorf, cat. nos. 5424 and 5804)
- Eppendorf ThermoStat C (Eppendorf, cat. no. 5383000019)
- Gene Pulser Xcell Microbial System (Bio-Rad, cat. no. 1652662)
- Digital gel imaging system (GelDoc EZ, Bio-Rad, cat. no. 170-8270), and blue sample tray (Bio-Rad, cat. no. 170-8273)
- Blue-light transilluminator and orange filter goggles (SafeImager 2.0; Invitrogen, cat. no. G6600)
- Gel quantification software (Bio-Rad, ImageLab or open-source ImageJ frmo the National Institutes of Health (NIH), USA, available at http://rsbweb.nih.gov/ij/)
- UV spectrophotometer (e.g. NanoDrop 2000c, Thermo Scientific)
- Plate spectrophotometer (e.g. Synergy H4 Hybrid Multi-Mode Microplate Reader, BioTek)
- Qubit Assay Tubes (Thermo Fisher, cat. no. Q32856)
- Qubit Fluorometer (Thermo Fisher, cat. no. Q33216)
- MiSeq System (Illumina, cat. no. SY-410-1003)
- NextSeq 500/550 System (Illumina, cat. nos. SY-415-1001 and SY-415-1002)
- Cell counter (e.g. Cellometer Image Cytometer, Nexcelom Bioscience)
- Sorvall Legend XTR Centrifuge (Thermo Fisher, cat. no. 75004520)

## REAGENT SETUP

**TBE electrophoresis solution** Dilute TBE buffer in distilled water to a 1 × working condition, and store it at room temperature (18-22 °C) for up to 6 months.

**Ethanol, 80% (vol/vol)** Prepare 80% (vol/vol) ethanol in UltraPure water right before use.

**D10 medium** For culture of HEK 293FT cells, prepare D10 medium by supplementing DMEM with GlutaMAX and 10% (vol/vol) FBS. For routine cell line culture and maintenance, D10 can be further supplemented with 1 × penicillin-streptomycin. Store the medium at 4 °C for up to 1 month.

**mTeSR1 medium** For culture of human embryonic stem cells (hESCs), prepare mTeSR1 medium by supplementing it with the supplement supplied with the medium and 100 μg ml^-1^ Normocin. Prepared medium can be stored at 4 °C for up to 2 months.

**Proteinase K, 300 U ml^-1^** Resuspend 25 mg of Proteinase K in 2.5 ml of 10 mM Tris, pH 8.0 for 10 mg ml^-1^ (300 U ml^-1^) of Proteinase K. Store at 4 °C for up to 1 year.

**Deoxyribonuclease I, 50 KU ml^-1^** Resuspend 50 KU of Deoxyribonuclease I in a solution containing 50% (vol/vol) Glycerol, 10 mM CaCl_2_, and 50 mM Tris-HCl (pH 7.5) for 50 KU ml^-1^ of Deoxyribonuclease I. See **Table S1** for a detailed setup of the Deoxyribonuclease I storage solution. Store at −20 °C for up to 1 year.

**RNA lysis buffer** Prepare an RNAse-free solution of 9.6 mM Tris-HCl (pH 7.8), 0.5 mM MgCl_2_, 0.44 mM CaCl_2_, 10 μM DTT, 0.1% (wt/vol) Triton X-114, and 3 U ml^-1^ Proteinase K in UltraPure water. The final pH of the solution should be approximately 7.8. Store at 4 °C for up to 1 year. See **Table S2** for a detailed setup.

**RNA lysis stop solution** Prepare solution under RNAse-free conditions. Resuspend 10 mg of Proteinase K Inhibitor in 150 μl of DMSO for a final concentration of 100 mM. Prepare 0.5 M EGTA (pH 8.3) in UltraPure water. **Critical** EGTA is light sensitive, and can be stored at 4 °C protected from light for up to 2 years. Combine for a final solution with 1 mM Proteinase K inhibitor, 90 mM EGTA, and 113 μM DTT in UltraPure water. Aliquot into 8-strip PCR tubes to avoid freeze-thaw and facilitate sample processing with multichannel pipettes. Store at −20 °C for up to 1 year. See **Table S3** and **Table S4** for a detailed setup. **Oligo dT, 100 μM** Resuspend oligo dT to 100 μM in UltraPure water. Aliquot and store at −20 °C for up to 2 years.

## EQUIPMENT SETUP

**Large LB agar plates (245 mm square bioassay dish, ampicillin)** Reconstitute the LB Broth with agar at a concentration of 35 g L^-1^ in deionized water and swirl to mix. Autoclave to sterilize. Allow the LB agar to cool to 55 °C before adding ampicillin to a final concentration of 100 μg ml^-1^ and swirl to mix. On a sterile bench area, pour ~300 ml of LB agar per 245 mm square bioassay dish. Place the lids on the plates and allow them to cool for 30-60 min until solidified. Invert the plates and let sit for several more hours or overnight. Agar plates can be stored in plastic bags or sealed with parafilm at 4 °C for up to 3 months.

**Standard LB agar plates (100 mm Petri dish, ampicillin)** Preparation of standard LB agar plates is similar to large LB agar plates, except pour ~20 ml of LB agar per 100 mm Petri dish. Store at 4 °C for up to 3 months.

## PROCEDURE

Prior to performing the screen, construct a pooled sgRNA library by designing and cloning a custom sgRNA library (Steps 1-20) or amplifying a ready-made library from Addgene (Skip to Step 21).

### Designing a targeted sgRNA library o TIMING 1 d

*1. Input target genes for library design*. We provide a python script design_oligos.py that extracts the sgRNA spacers that target an input set of genes from the genome-scale screen. Once a set of genes for the targeted screen has been identified, prepare a csv file containing the names of the target genes with each line corresponding to one gene. Prepare another csv file for the annotated genome-scale library with the names of each gene in the first column and respective spacer sequences in the second column. Each line contains a different spacer sequence. The gene names in the target genes file should be in the same format as the names of the annotated library file.

2. Isolate the subset of spacers from the genome-scale library that correspond to the target genes by running python design_oligos.py with the following optional parameters:

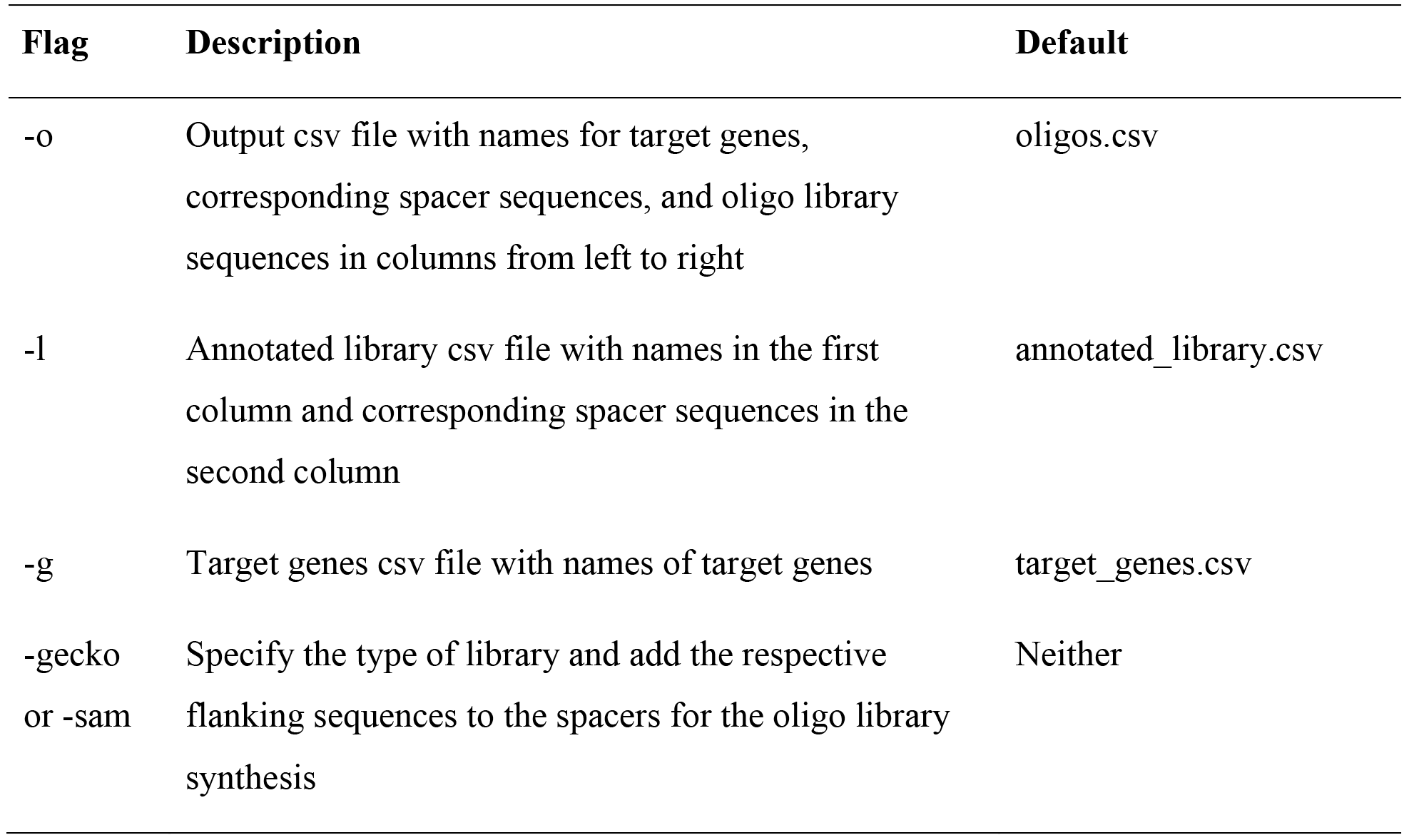

3. After running design_oligos.py, the subset of spacers for the target genes will be written to an output csv file. If-gecko or-sam is specified, the full oligo library sequence containing the spacers and respective flanking sequences for synthesis will be in the last column.

4. Synthesize the oligo library as a pool on an array through a DNA synthesis platform such as Twist Bioscience or CustomArray. Parafilm and store pooled oligos at −20 °C.

### Cloning a custom sgRNA library o TIMING 2 d

5. Throughout the sgRNA library cloning process, refer to the table below for the number of reactions recommended at each cloning step for a library size of 100,000 sgRNAs and scale the number of reactions according to the size of the targeted sgRNA library.

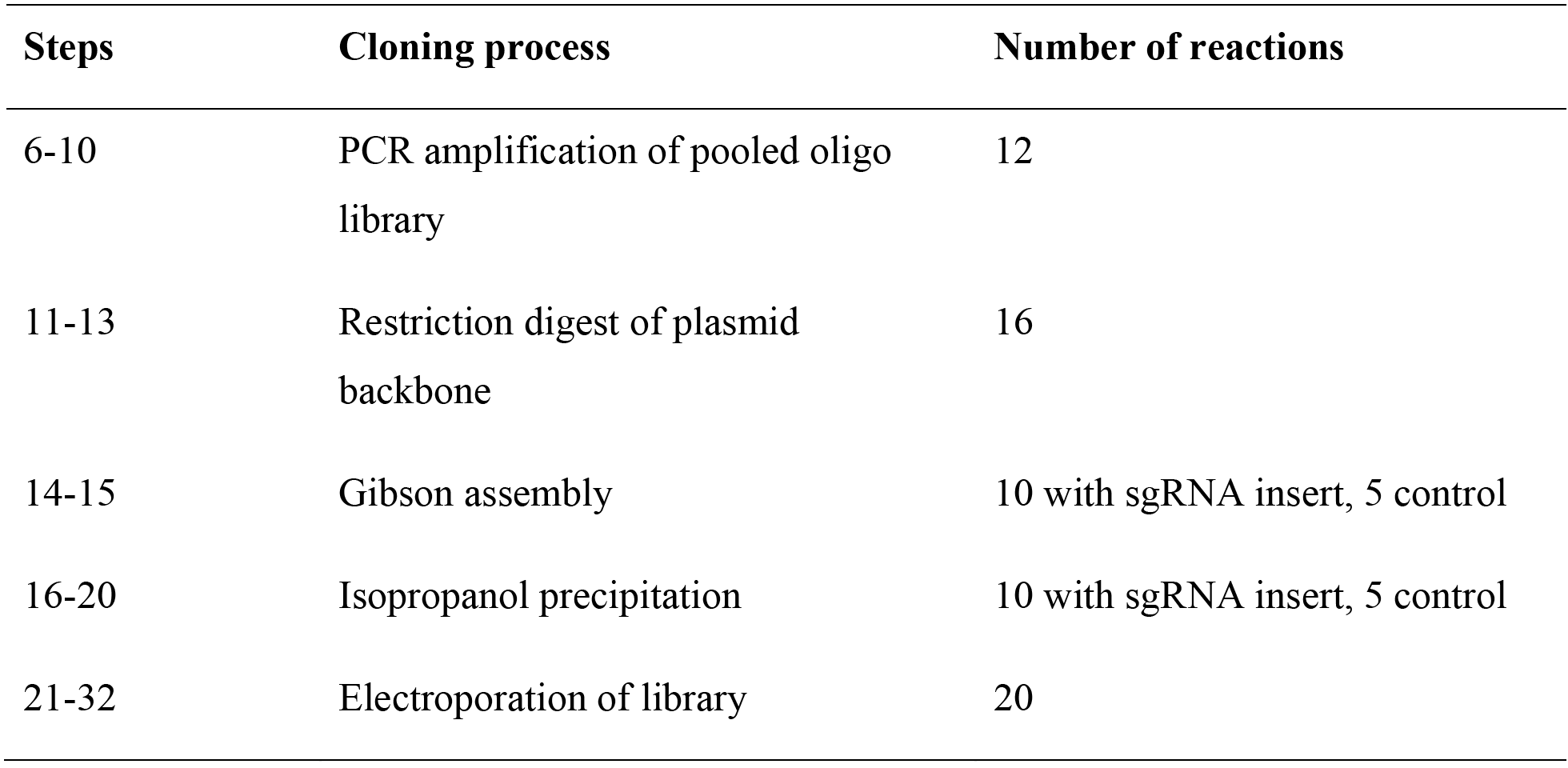

*6. PCR amplification of pooled oligo library*. Amplify the pooled oligo library using the Oligo-Fwd and Oligo-Rev primers (**Table 2**). Prepare a master mix using the reaction ratios outlined below:

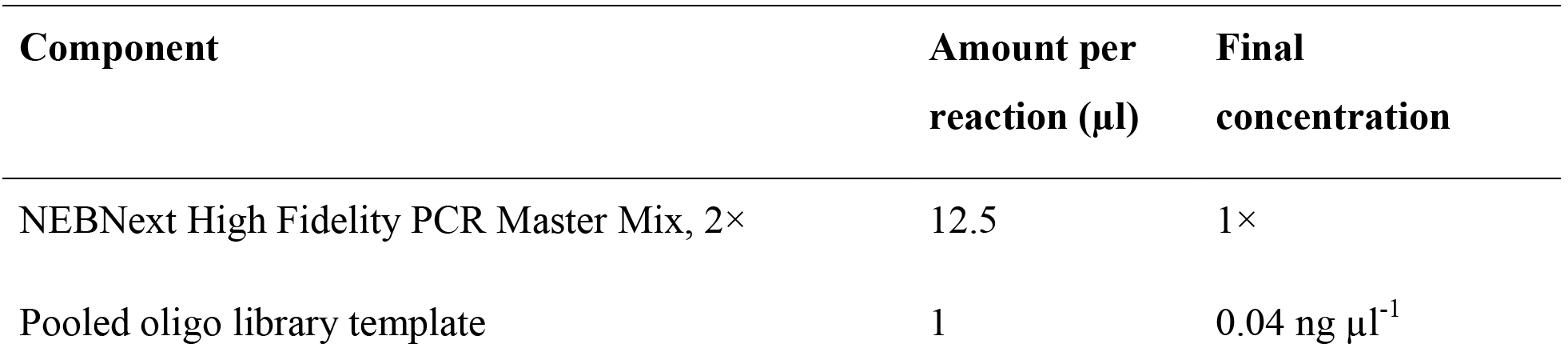

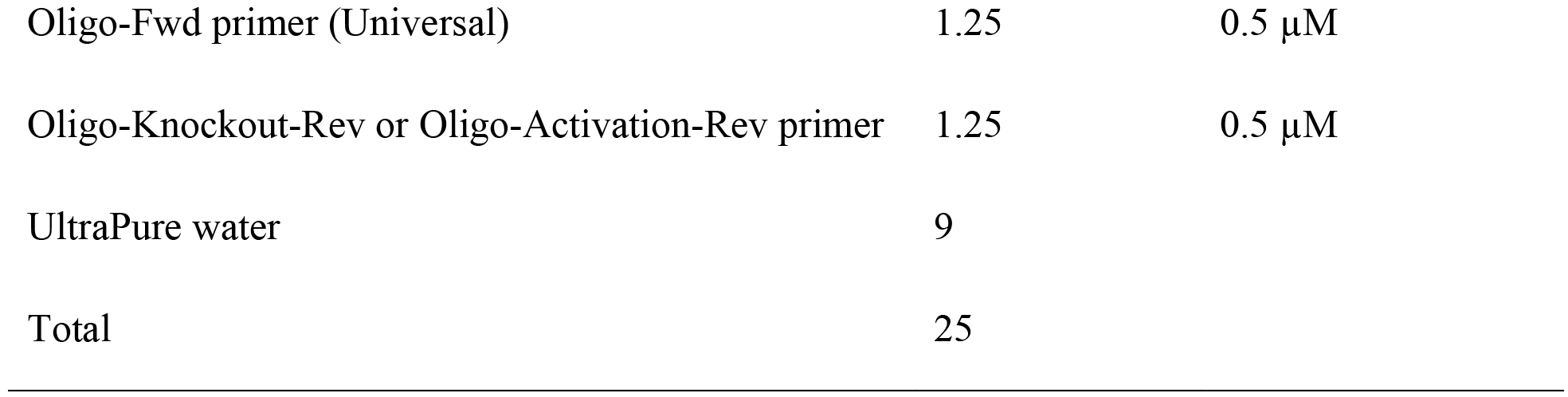

**Critical Step** To minimize error in amplifying oligos, it is important to use a high-fidelity polymerase. Other high-fidelity polymerases, such as PfuUltra II (Agilent) or Kapa HiFi(Kapa Biosystems), may be used as a substitute.

7. Aliquot the PCR master mix into 25 μl reactions and perform a PCR by using the following cycling conditions:

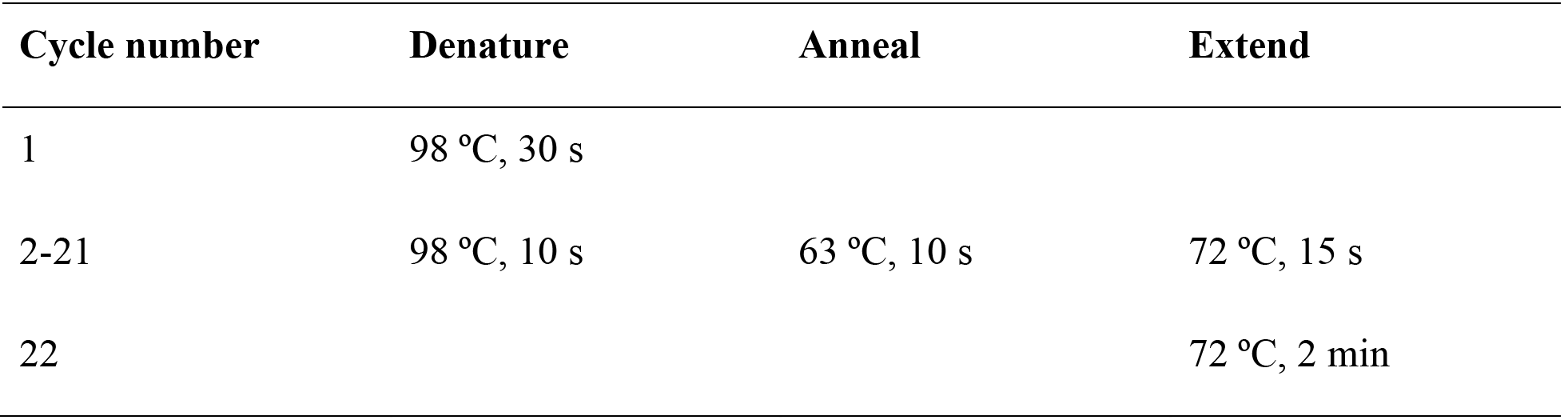

**Critical Step** Limit the number of PCR cycles to 20 cycles during amplification to reduce potential biases introduced during amplification.

8. After the reaction is complete, pool the PCR reactions and purify the PCR product by using the QIAquick PCR purification kit according to the manufacturer’s directions.

9. Run PCR purified oligo library on a 2% (wt/vol) agarose gel along with a 50bp ladder.

**Critical Step** Run on a 2% (wt/vol) agarose gel for long enough to separate the target library (140bp) from a possible primer dimer of ~120bp. Under the optimized PCR conditions suggested above the presence of primer dimers should be minimal.

10. Gel extract the purified PCR product using the QIAquick gel extraction kit according to the manufacturer’s directions and quantify.

11. *Restriction digest of plasmid backbone*. Digest the desired library plasmid backbone with the restriction enzyme Esp3I (BsmBI) that cuts around the sgRNA target region. Refer to the master mix set up below for the reaction ratios:

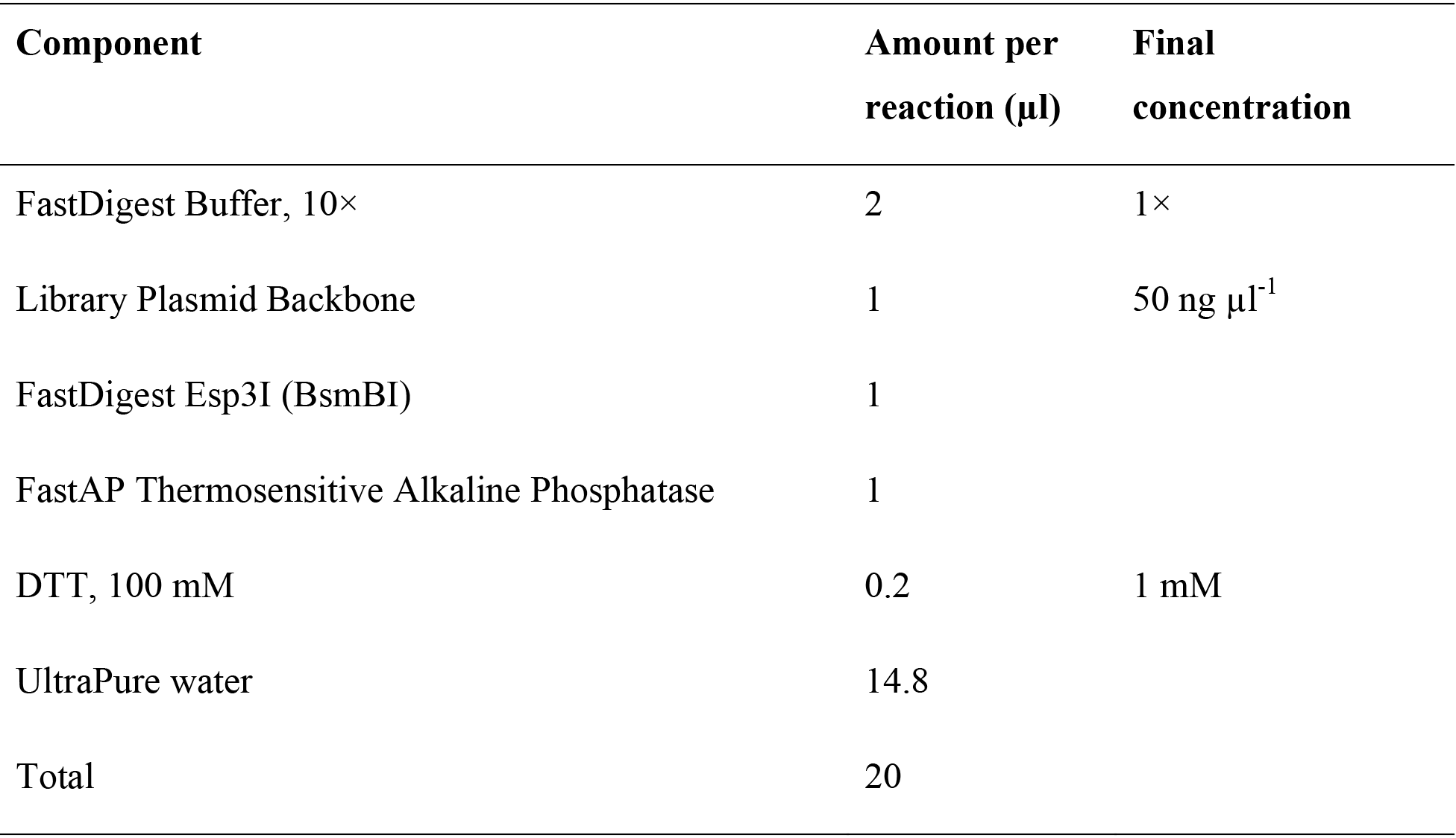

12. Aliquot 20 μl reactions from the master mix and incubate the restriction digest reaction at 37 °C for 1 h.

13. After the reaction has completed, run the restriction digest on a 2% (wt/vol) agarose gel and gel extract the library plasmid backbone using the QIAquick gel extraction kit according to the manufacturer’s protocol and quantify. Note that the Gecko library backbones contain a 1880bp filler sequence which should be visible as a dropout. The SAM library backbones do not contain a filler sequence and the expected dropout of 20bp is usually not readily visible.

14. *Gibson Assembly*. Set up a master mix for the Gibson reactions on ice according to the reaction ratios below. Be sure to include reactions without the sgRNA library insert as a control.

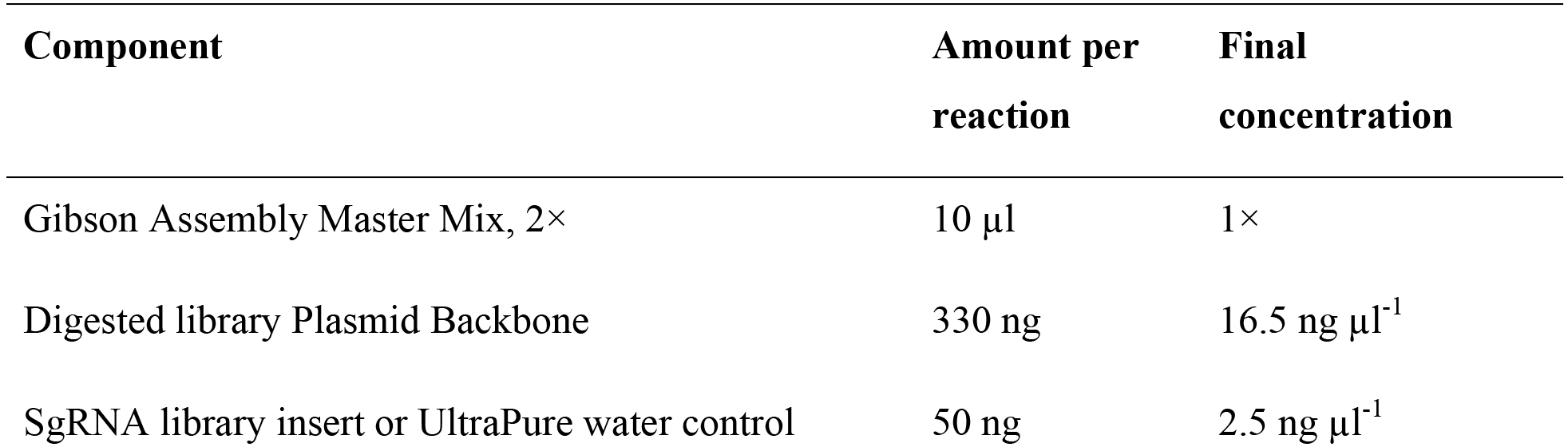

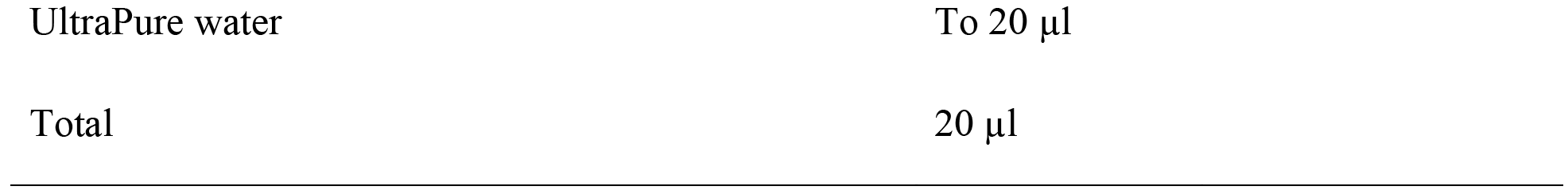

15. Aliquot 20 μl reactions from the master mix and incubate the Gibson reaction at 50 °C for 1h.

**Pause Point** Completed Gibson reactions can be stored at −20 °C for at least 1 week.

16. *Isopropanol precipitation*. Pool cloning and control reactions separately. Purify and concentrate the sgRNA library by mixing the following:

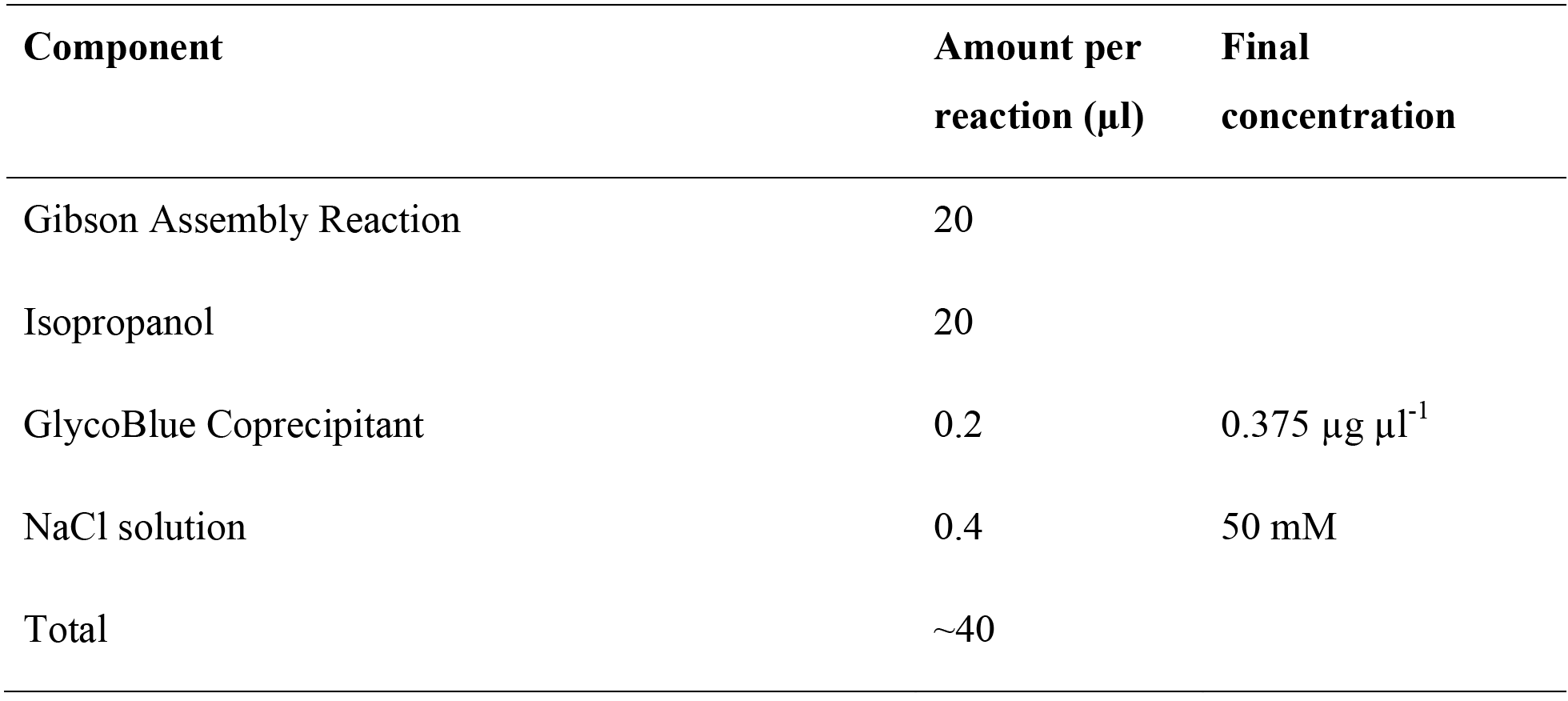

**Critical Step** In addition to concentrating the library, purification by isopropanol precipitation removes salts from the Gibson reaction that can interfere with electroporation.

17. Vortex and incubate at room temperature for 15 min and centrifuge at >15,000 × g for 15 min to precipitate the plasmid DNA. The precipitated plasmid DNA should appear as a small light blue pellet at the bottom of the microcentrifuge tube.

18. Aspirate the supernatant and gently wash the pellet twice without disturbing it using 1 ml of ice-cold (−20 °C) 80% (vol/vol) ethanol in UltraPure water.

19. Carefully remove any residual ethanol and air dry for for 1 min.

20. Resuspend the plasmid DNA pellet in 5 μl of TE per reaction by incubating at 55 °C for 10 min and quantify the targeted sgRNA library by nanodrop. Isopropanol-purified sgRNA libraries can be stored at −20 °C for several months.

### Amplification of pooled sgRNA library o TIMING 2 d

21. *Pooled sgRNA library transformation*. Electroporate the library at 50-100 ng μl^-1^ using Endura ElectroCompetent cells according to the manufacturer's directions. If amplifying a ready-made genome-scale library from Addgene, repeat for a total of 8 electroporations. If amplifying a targeted sgRNA library refer to Step 5 to scale the amplification steps appropriately and include an additional electroporation for the control Gibson reaction without sgRNA library insert.

22. Pre-warm 1 standard LB agar plate (100 mm Petri dish, ampicillin) and large LB agar plates (245 mm square bioassay dish, ampicillin). Each large LB agar plate can be substituted with 10 standard LB agar plates. For amplification of a targeted sgRNA library, include an additional standard LB agar plate for the control Gibson reaction.

23. After the recovery period, pool electroporated cells and mix well by inverting.

24. Plate a dilution for calculating transformation efficiency. To prepare the dilution mix, add 10 μl of the pooled electroporated cells to 990 μl of LB medium for a 100-fold dilution and mix well. Then add 100 μl of the 100-fold dilution to 900 μl of LB medium for a 1,000-fold dilution and mix well.

25. Plate 100 μl of the 1,000-fold dilution onto a pre-warmed standard LB agar plate (10 cm Petri dish, ampicillin). This is a 10,000-fold dilution of the full transformation that will be used to estimate the transformation efficiency. If amplifying a targeted sgRNA library, repeat Steps 24-25 for the control Gibson reaction.

26. Plate pooled electroporated cells. Add 1 volume of LB medium to the pooled electroporated cells, mix well, and plate on large LB agar plates (option A) or standard LB agar plates (option B).

a. Plate 2 ml of electroporated cells on each of the pre-warmed large LB agar plates using a cell spreader. Spread the liquid culture until it is largely absorbed into the agar and does not drip when the plate is inverted. At the same time, make sure the liquid culture does not completely dry out as this will lead to poor survival.
b. Alternatively, plate 200 μl of electroporated cells on each of the prewarmed standard LB agar plates using the same technique as described in Step 26a.

**Critical Step** Plating the electroporated cells evenly is important for preventing intercolony competition that may skew the sgRNA library distribution.

27. Incubate all LB agar plates overnight at 37 °C for 12-14 h.

**Critical Step** Limiting the bacterial growth time to 12-14 h ensures that there is sufficient growth for sgRNA library amplification without potentially biasing the sgRNA library distribution through intercolony competition or differences in colony growth rates.

28. Calculate electroporation efficiency.

a. Count the number of colonies on the dilution plate.
b. Multiply the number of colonies by 10,000 and the number of electroporations to obtain the total number of colonies on all plates. If amplifying a ready-made sgRNA library from Addgene, proceed if the total number of colonies is greater than 100 colonies per sgRNA in the library. If amplifying a custom targeted sgRNA library, proceed if there are more than 500 colonies per sgRNA in the library. **Critical Step** Obtaining a sufficient number of colonies per sgRNA is crucial for ensuring that the full library representation is preserved and that sgRNAs do not drop out during amplification.
c. In addition, for amplification of a targeted sgRNA library, calculate the electroporation efficiency for the control Gibson reaction and proceed if there are at least 20 times more colonies per electroporation in the sgRNA library condition compared to the control Gibson reaction.

### Troubleshooting

29. *Harvest colonies from the LB agar plates*. Pipette 10 ml of LB medium onto each large LB agar plate or 1 ml of LB medium onto each standard LB agar plate. Gently scrape the colonies off with a cell spreader and transfer the liquid with scraped colonies into a 50-ml Falcon tube.

30. For each LB agar plate, repeat Step 29 for a total of 2 LB medium washes to capture any remaining bacteria.

31. Maxiprep the amplified sgRNA library by using the Macherey-Nagel NucleoBond Xtra Maxi EF according to the manufacturer’s directions. Calculate the number of maxipreps needed by measuring the 0D600 of the harvested bacterial suspension as follows: Number of maxipreps = 0D600*(total volume of suspension)/1200.

**Critical Step** Using an endotoxin-free plasmid purification kit is important for avoiding endotoxicity in virus production and mammalian cell culture.

32. Pool the resulting plasmid DNA and quantify. Maxiprepped sgRNA library can be aliquoted and stored at −20 °C.

### Next-generation sequencing of the amplified sgRNA library to determine sgRNA distribution o TIMING 2-3 d

33. *Library PCR for NGS*. We have provided NGS primers that amplify the sgRNA target region with Illumina adapter sequences (**Table 3**). To prepare the sgRNA library for NGS, set up a reaction for each of the 10 NGS-Lib-Fwd primers and 1 NGS-Lib-KO-Rev or NGS-Lib-SAM-Rev barcode primer as follows:

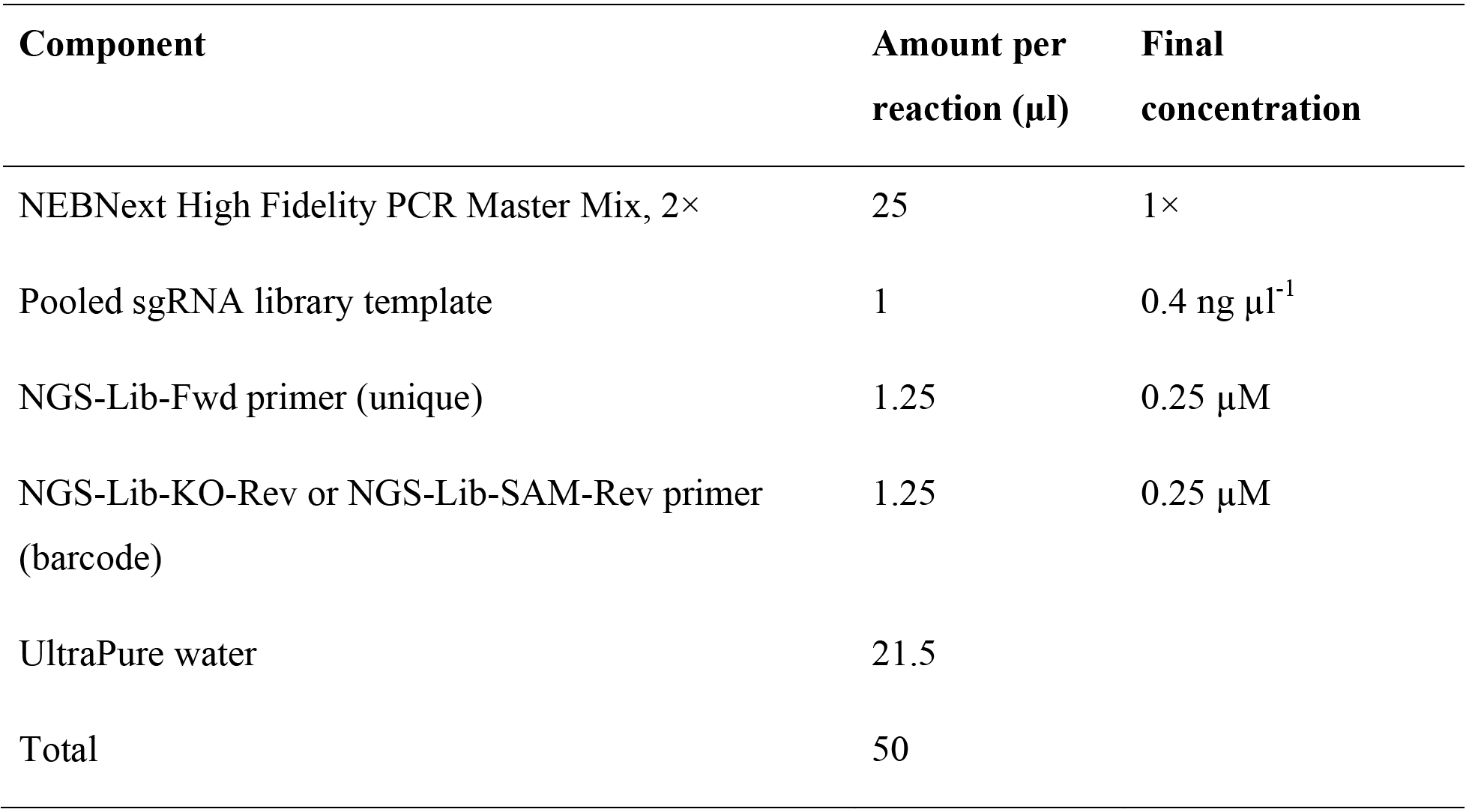

**Critical step** Using a different reverse primer with a unique barcode for each library allows for pooling and sequencing of different libraries in a single NextSeq or HiSeq run. This is more efficient and cost-effective than running the same number of libraries on multiple Miseq runs.

**Critical Step** To minimize error in amplifying sgRNAs, it is important to use a high-fidelity polymerase. Other high-fidelity polymerases, such as PfuUltra II (Agilent) or Kapa HiFi(Kapa Biosystems), may be used as a substitute.

34. Perform a PCR by using the following cycling conditions:

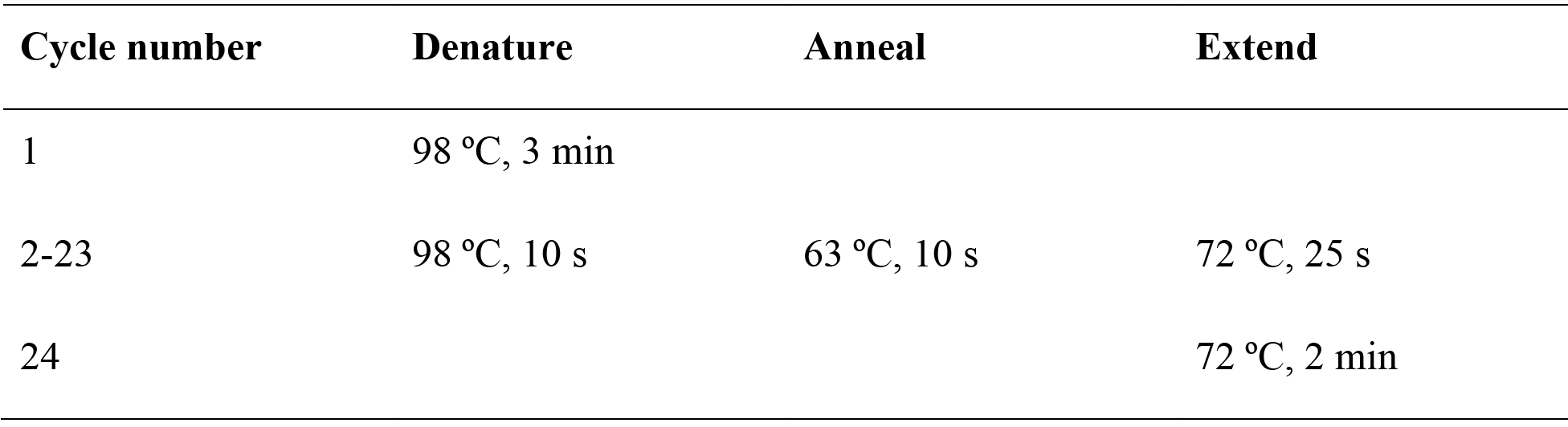

35. After the reaction is complete, pool the PCR reactions and purify the PCR product by using the QIAquick PCR purification kit according to the manufacturer’s directions.

36. Quantify the purified PCR product and run 2 μg of the product on a 2% (wt/vol) agarose gel. Successful reactions should yield a ~260-270bp product for the knockout library and a ~270-280bp product for the activation library. Gel extract using the QIAquick gel extraction kit according to the manufacturer’s directions.

**Pause Point** Gel-extracted samples can be stored at −20 °C for several months.

37. Quantify the gel-extracted samples using the Qubit dsDNA HS Assay Kit according to the manufacturer’s instructions.

38. Sequence the samples on the Illumina MiSeq or NextSeq according to the Illumina user manual with 80 cycles of read 1 (forward) and 8 cycles of index 1. We recommend sequencing with 5% PhiX control on the MiSeq or 20% PhiX on the NextSeq to improve library diversity and aiming for a coverage of >100 reads per sgRNA in the library.

39. *Analyze sequencing data with count spacers.py*. Install biopython (http://biopython.org/DIST/docs/install/Installation.html). Prepare a csv file containing the guide spacer sequences with each line corresponding to one sequence.

40. To determine the spacer distribution, run python count_spacers.py with the following optional parameters:

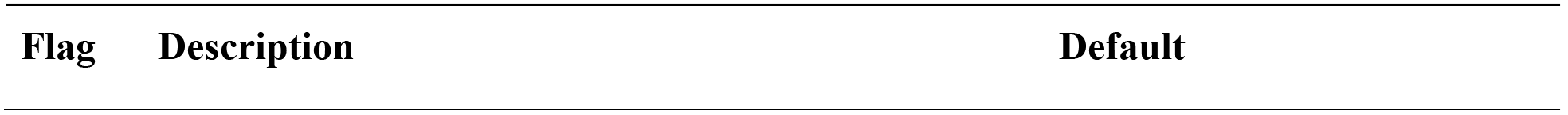

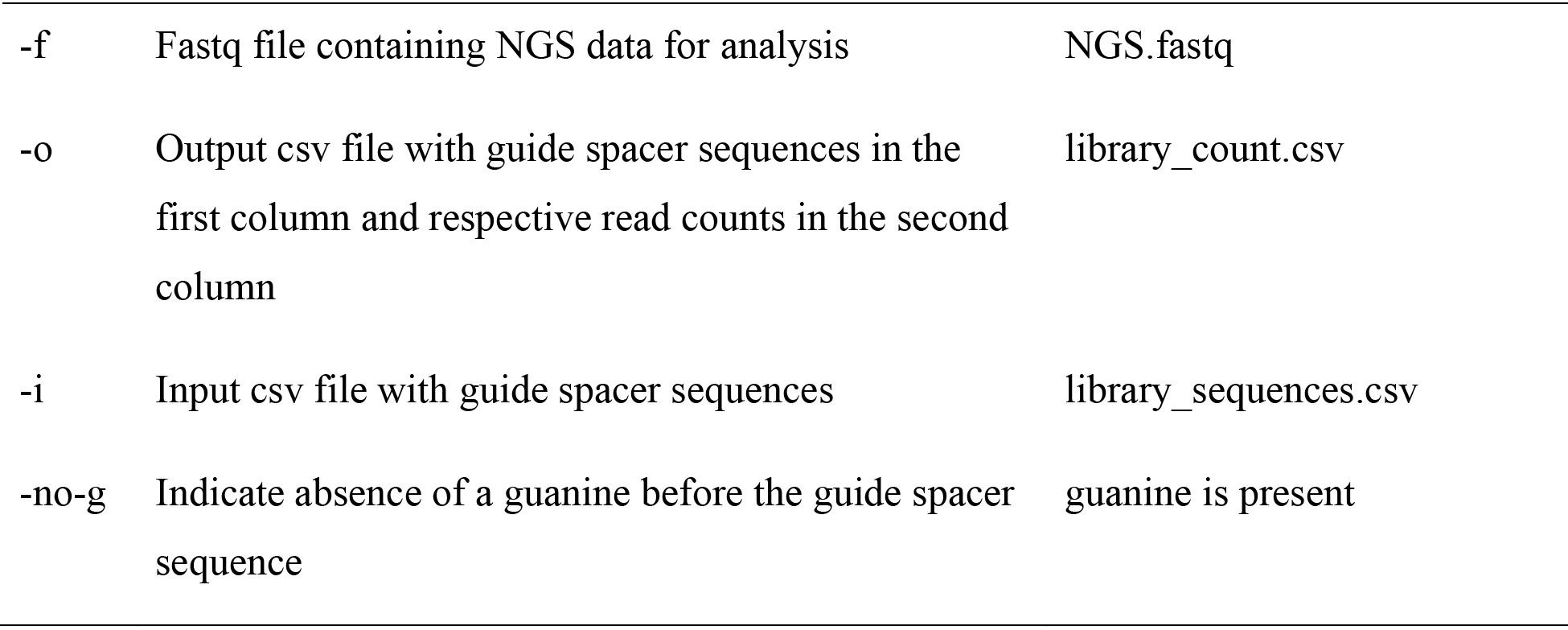

Place all relevant files are in the same folder before running python count_spacers.py. The human SAM libraries do not have a guanine before the guide spacer sequence, so make sure to run the script with the parameter-no-g when analyzing those libraries.

41. After running count_spacers.py, spacer read counts will be written to an output csv file. Relevant statistics including the number of perfect guide matches, non-perfect guide matches, sequencing reads without key, the number of reads processed, percentage of perfectly matching guides, percentage of undetected guides, and skew ratio will be written to statistics.txt. An ideal sgRNA library should have more than 70% perfectly matching guides, less than 0.5% undetected guides, and a skew ratio of less than 10.

### Troubleshooting

#### Lentivirus production and titer o TIMING 8-10 d

42. *HEK 293FT maintenance*. Cells are cultured in D10 medium at 37 °C with 5% CO_2_ and maintained according to the manufacturer’s recommendation.

43. To passage, aspirate the medium and rinse the cells by gently adding 5 ml of TrypLE to the side of the T225 flask, so as not to dislodge the cells. Remove the TrypLE and incubate the flask for 4-5 min at 37 °C until the cells have begun to detach. Add 10 ml of warm D10 to the flask and dissociate the cells by pipetting them up and down gently, and transfer the cells to a 50-ml Falcon tube.

**Critical Step** We typically passage cells every 1-2 d at a split ratio of 1:2 or 1:4, never allowing cells to reach more than 70% confluency. For lentivirus production, we recommend using HEK 293FT cells with a passage number less than 10.

44. *Preparation of cells for transfection*. Seed the well-dissociated cells into T225 flasks 20-24 h before transfection at a density of 1.8 × 10^7^ cells per flask in a total volume of 45 ml D10 medium. Plate 5 T225 flasks for knockout screening and 9 T225 flasks for activation screening.

**Critical Step** Do not plate more cells than the recommended density, as doing so may reduce transfection efficiency.

45. *Lentivirus plasmid transfection*. Cells are optimal for transfection at 80-90% confluency. Transfect 4 T225 flasks with the sgRNA library and 1 T225 flask with a constitutive GFP expression plasmid as a transfection control. If transfecting for activation screening, transfect an additional 4 T225 flasks with the Cas9 activator components that are not in the sgRNA library backbone, i.e. MS2-p65-HSF1. We outline below a transfection method using Lipofectamine 2000 and PLUS reagent. Alternatively, we describe a cost-effective method for lentivirus transfection with Polyethylenimine (PEI) in **Box 5**.

**Critical Step** Transfecting at the recommended cell density is crucial for maximizing transfection efficiency. Lower densities can result in Lipofectamine 2000 toxicity for cells, while higher densities can reduce transfection efficiency.

a. For each lentiviral target, combine the following lentiviral target mix in a 15-ml or 50-ml Falcon tube and scale up accordingly:

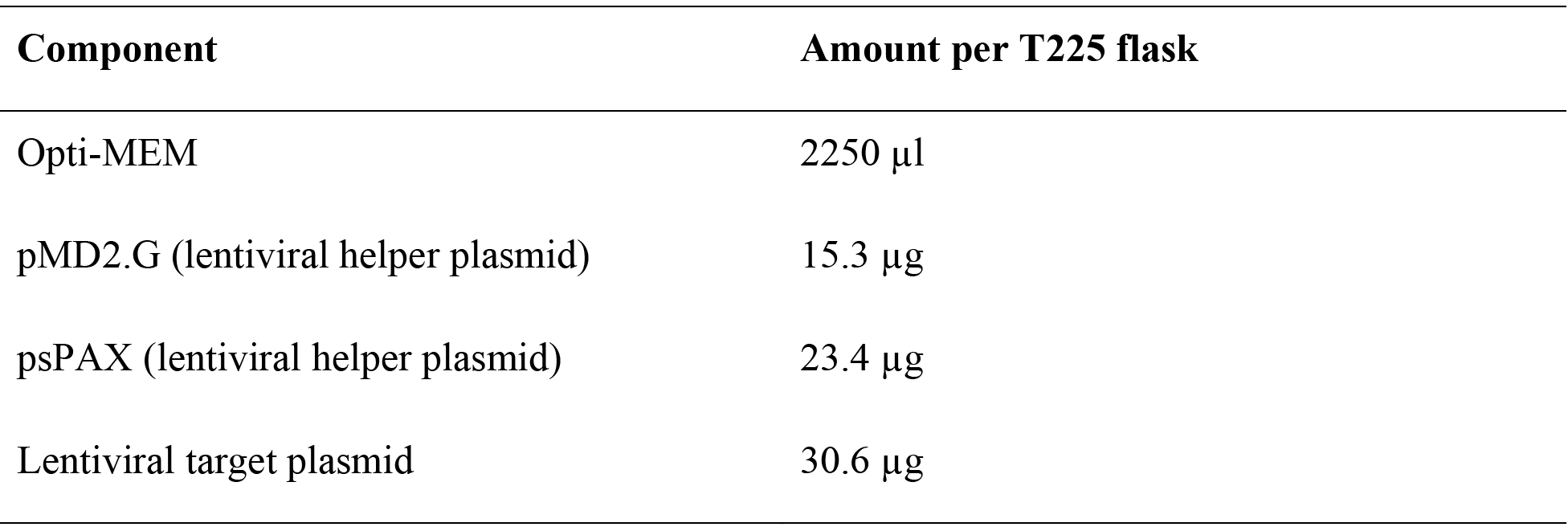
b. Prepare the PLUS reagent mix as follows and invert to mix:

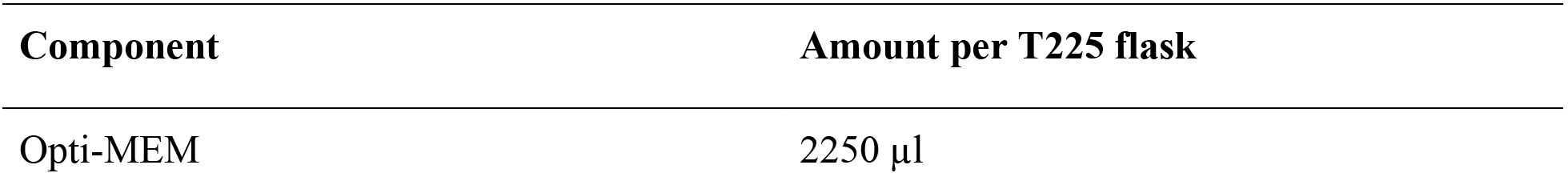

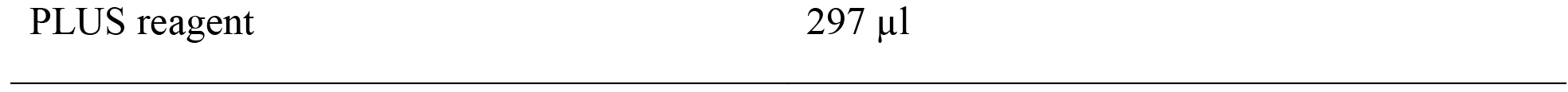
c. Add the PLUS reagent mix to the lentiviral target mix, invert, and incubate at room temperature for 5 min.
d. Prepare the Lipofectamine reagent mix as follows and invert to mix:

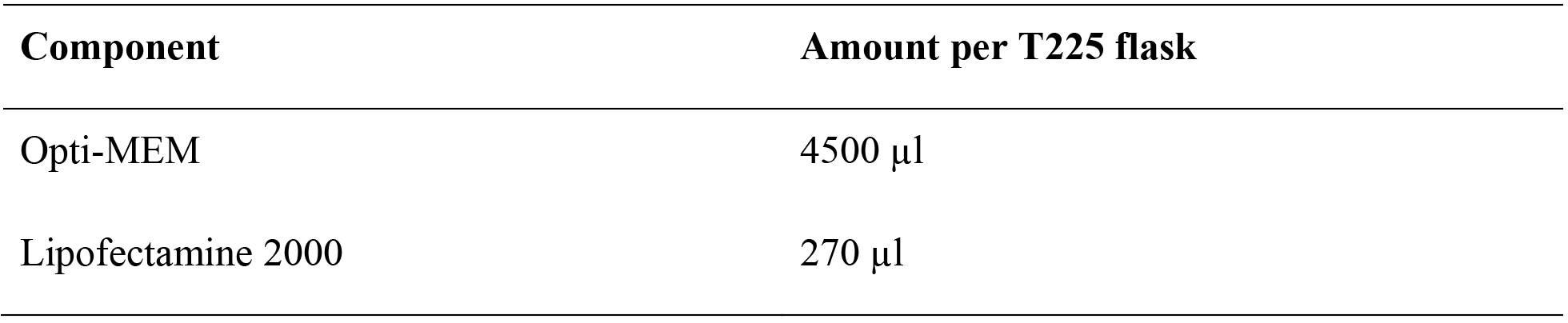
e. Add the lentiviral target and PLUS reagent mix to the Lipofectamine reagent mix, invert, and incubate at room temperature for 5 min.
f. Pipette 9 ml of the lentiviral transfection mix into each T225 flask and shake gently to mix. Return the T225 flasks to the incubator.
g. After 4 h, replace the medium with 45 ml of pre-warmed D10 medium. The constitutive GFP expression plasmid transfection control indicates transfection efficiency.

### Troubleshooting

46. *Harvest and store lentivirus*. 2 d after the start of lentiviral transfection, pool the lentivirus supernatant from the T225 flasks with the same lentivirus and filter out cellular debris using Millipore’s 0.45 μm Stericup filter unit.

**Pause Point** The filtered lentivirus supernatant can be aliquoted and stored at -80 °C. Avoid freeze-thawing lentivirus supernatant.

47. *Determine the lentiviral titer through transduction*. The CRISPR-Cas9 system has been used in a number of mammalian cell lines. Conditions may vary for each cell line. Lentiviral titer should be determined using the relevant cell line for the screen. Below we detail transduction conditions and calculation of viral titer for HEK 293FT cells (option A) and hESC HUES66 cells (option B).

**Critical Step** We recommend performing a kill curve for the antibiotic used to select the sgRNA library on your cells prior to determination of lentiviral titer. It is important to use the lowest concentration of antibiotic sufficient to to kill the negative control within 4-7 days to avoid excessively stringent selection that biases selection for cells transduced with multiple sgRNAs.

a. Lentiviral transduction and titer for HEK 293FT cells by spinfection

i. *HEK 293FT maintenance and passaging*. Refer to steps 42-43.
ii. *Preparation of cells for spinfection*. For each lentivirus, seed 6 wells of a 12-well plate at a density of 3 × 10^6^ cells in 2 ml D10 medium per well with 8 μg ml^-1^ of polybrene. In each well, add 400 μl, 200 μl, 100 μl, 50 μl, 25 μl, or 0 μl of lentivirus supernatant. Mix each well thoroughly by pipetting up and down.
iii. Spinfect the cells by centrifuging the plates at 1000 × g for 2 h at 33 °C. Return the plates to the incubator after spinfection.
iv. *Replating spinfection for calculation of viral titer*. 24 h after the end of spinfection, remove the medium, gently wash with 400 μl TrypLE per well, add 100 μl of TrypLE, and incubate at 37 °C for 5 min to dissociate the cells. Add 2 ml of D10 medium per well and resuspend the cells by pipetting up and down.
v. Determine the cell concentration for the 0 μl lentivirus supernatant condition.
vi. For each virus condition, seed 4 wells of a 96-well clear bottom black tissue culture plate at a density of 4 × 10^3^ cells based on the cell count determined in the previous step in 100 μl of D10 medium. Add an additional 100 μl of D10 medium with the corresponding selection antibiotic for the virus at an appropriate final concentration to 2 wells and 100 μl of regular D10 medium to the other 2 wells for a total of 2 bioreps per virus condition.
vii. 72-96 h after replating, when the no virus conditions contain no viable cells and the no antibiotic selection conditions are at 80-90% confluency, quantify the cell viability for each condition using CellTiter Glo according to the manufacturer’s protocol. We have found that Cell Titer Glo can be diluted 1:4 in PBS to reduce cost while still achieving optimal results.
viii. For each virus condition, the multiplicity of infection (MOI) is calculated as the luminescence, or viability, of the condition with antibiotic selection divided by the condition without antibiotic selection. A linear relationship between lentivirus supernatant volume and MOI is expected at lower volumes, with saturation achieved at higher volumes.
b. Lentiviral transduction and titer for hESC HUES66 cells by mixing

i. *HUES66 maintenance*. We routinely maintain HUES66 cells (a hESC cell line) in feeder-free conditions with mTeSR1 medium on GelTrex-coated tissue culture plates. To coat a 100-mm tissue culture dish, dilute cold GelTrex 1:100 in 5 ml of cold DMEM, cover the entire surface of the culture dish, and place the dish in the incubator for at least 30 min at 37 °C. Aspirate the GelTrex mix prior to plating. During passaging and plating, mTeSR1 medium is supplemented further with 10 μM ROCK inhibitor. The mTeSR1 medium is refreshed daily.
ii. *Passaging HUES66*. Aspirate the medium and rinse the cells once by gently adding 10 ml of DPBS to the side of the 100-mm tissue culture dish, so as not to dislodge the cells. Dissociate the cells by adding 2 ml of Accutase and incubate at 37 °C for 3-5 min until the cells have detached. Add 10 ml of DMEM, resuspend the dissociated cells, and pellet the cells at 200 × g for 5 min. Remove the supernatant, resuspend the cells in mTeSR1 medium with 10 μM ROCK inhibitor and replate the cells onto GelTrex-coated plates. Replace with normal mTeSR1 medium 24 h after plating. **Critical Step** We typically passage cells every 4-5 d at a split ratio of 1:5 or 1:10, never allowing cells to reach more than 70% confluency.
iii. *Preparation of cells for lentiviral transduction*. For each lentivirus, plate 6 wells of a Geltrex-coated 6-well plate at a density of 5 × 10^5^ cells in 2 ml mTeSR1 medium per well. In each well, add 400 μl, 200 μl, 100 μl, 50 μl, 25 μl, or 0 μl of lentivirus supernatant, fill to a total volume to 3 ml with DPBS, and supplement with 10 μM ROCK inhibitor. Plate an additional no antibiotic selection control well at the same seeding density without virus. Mix each well thoroughly by pipetting up and down.
iv. 24 h after lentiviral transduction, replace the medium with mTeSR1 containing the relevant antibiotic selection. For the no antibiotic selection control well, replace the medium with normal mTeSR1 medium. Refresh the mTeSR1 medium with and without antibiotic selection every day until the plate is ready for the next step.
v. *Calculation of viral titer*. 72-96 h after starting the antibiotic selection, when the no virus condition contains no viable cells and the no antibiotic selection control condition is 80-90% confluent, rinse the cells with 2 ml of DPBS, add 500 μl of Accutase, and incubate at 37 °C for 3-5 min to dissociate the cells. Add 2 ml of DMEM and mix well.
vi. Count and record the number of cells in each well.
vii. For each virus condition, the MOI is calculated as the number of cells in the antibiotic selection condition divided by the number of cells in the no antibiotic selection control. A linear relationship between lentivirus supernatant volume and MOI is expected at lower volumes, with saturation achieved at higher volumes.

### Troubleshooting

#### Lentiviral transduction and screening o TIMING 3-6 w

Skip to Step 50 if performing a knockout screen.

48. *Generation of a cell line with stably expressed Cas9 activation components*. Prior to performing an activation screen, transduce the relevant cell line with the additional Cas9 activation components that are not present in the sgRNA library backbone at an MOI < 0.7. If two additional Cas9 activation components are required, both components can be transduced at the same time. Scale up as necessary to generate sufficient cells for maintaining sgRNA representation after sgRNA library transduction and selection. We have found that generation of a clonal line with Cas9 or SAM components is not necessary for successful screening. The cells can therefore be transduced and selected as a bulk population at the desired scale. Below we describe lentiviral transduction and cell line generation methods for HEK 293FT cells (option A) and hESC HUES66 cells (option B).

a. Generation of HEK 293FT cell lines

i. Seed cells in 12-well plates at a density of 3 × 10^6^ cells in 2 ml D10 medium per well with 8 μg ml^-1^ of polybrene. Add the appropriate volume of lentivirus supernatant to each well, and make sure to include a no virus control. Mix each well thoroughly by pipetting up and down.
ii. Spinfect the cells by spinning the plates at 1000 × g for 2 h at 33 °C. Return the plates to the incubator after spinfection.
iii. 24 h after the end of spinfection, remove the medium, gently wash with 400 μl TrypLE per well, add 100 μl of TrypLE, and incubate at 37 °C for 5 min to dissociate the cells. To each well, add 2 ml of D10 medium with the appropriate selection antibiotic for the lentivirus and resuspend the cells by pipetting up and down.
iv. Pool the resuspended cells from the wells with virus and seed the cells into T225 flasks at a density of 9 × 10^6^ cells per flask in 45 ml of D10 medium with selection antibiotic.
v. Transfer the resuspended cells from the no virus control into a T75 and add 13 ml of D10 medium with selection antibiotic.
vi. Refresh the selection antibiotic every 3 d and passage as necessary for 4-7 d and until there are no viable cells in the no virus control.
b. Generation of HUES66 cell lines

i. Seed cells in Geltrex-coated 6-well plates at a density of 5 × 10^5^ cells in of 2 ml mTeSR1 medium per well. Add the appropriate volume of lentivirus supernatant to each well, and make sure to include a no virus control. Fill up the total volume to 3 ml with DPBS, and supplement with 10 μM ROCK inhibitor. Mix each well thoroughly by pipetting up and down.
ii. 24 h after lentiviral transduction, replace the medium with mTeSR1 containing the relevant selection antibiotic. Refresh the mTeSR1 medium with selection antibiotic every day and passage as necessary for 4-7 d and until there are no viable cells in the no virus control.

**Critical Step** The lentiviral transduction method for generating a cell line for screening should be consistent with the method for titering the virus to ensure that cells are transduced at the appropriate MOI.

49. After selecting for successfully transduced cells, allow the cells to recover from selection by culturing in normal medium for 2-7 d before transducing with the sgRNA library. If culturing for more than 7 d after selection or after freezing cells, re-select the Cas9 activation cell line with the appropriate selection antibiotic to ensure expression of the Cas9 activation components and allow the cells to recover before sgRNA library transduction.

**Pause Point** Cells can be frozen down according to the manufacturer’s protocol.

50. *Transduction of cells with the sgRNA library*. Refer to Steps 48-49 for lentiviral transduction at the appropriate MOI and selection of transduced cell lines. To ensure that most cells receive only one genetic perturbation, transduce the sgRNA library at an MOI < 0.3. Scale up the transduction such that the sgRNA library has a coverage of >500 cells expressing each sgRNA. For example, for a library size of 100,000 unique sgRNAs, transduce 1.67 × 10^8^ cells at an MOI of 0.3. After the appropriate selection for 4-7 days, the cells are ready for screening. For knockout screening, we have found that maximal knockout efficiency is achieved 7 days after sgRNA transduction and therefore recommend selecting for 7 days before starting the screen selection. In contrast, maximal SAM activation is achieved as early as 4 days after sgRNA transduction. If selection is complete based on the no virus control, gain-of-function screening can be started 5 days after transduction. We generally recommend to perform 4 independent screening replicates (i.e. 4 separate sgRNA library infections followed by separate screening selection). Multiple bioreps are critical for determining screening hits with a high rate of validation.

**Critical Step** It is important to aim for a coverage of >500 cells per sgRNA to guarantee that each perturbation will be sufficiently represented in the final screening readout. Increase the coverage as necessary if the screening selection pressure is not very strong or if performing a negative selection screen. Transducing the sgRNA library at an MOI < 0.3 ensures that most cells receive at most one genetic perturbation. Transducing at higher MOI’s may confound screening results.

51. Since the parameters of each screen depends on the screening phenotype of interest, we provide guidelines and technical considerations for the screen (**Box 2-4**).

### Harvest genomic DNA for screening analysis o TIMING 3-4 d

52. *Harvest genomic DNA*. At the end of the screen, harvest genomic DNA (gDNA) from a sufficient number of cells to maintain a coverage of >500. For a library size of 100,000 unique sgRNAs, harvest gDNA from at least 5 × 10^7^ cells for downstream sgRNA analysis using the Zymo Research Quick-gDNA MidiPrep according to the manufacturer’s protocol. Make sure to tighten the connection between the reservoir and the column and centrifuge at a sufficient speed and time to remove any residual buffer. Addition of a final dry spin is recommended to remove residual wash buffer. Elution should be performed twice with 150-200 μl each for maximum recovery of gDNA.

**Pause Point** Frozen cell pellets or isolated gDNA can be stored at −20 °C for several months.

53. *Preparation of the gDNA for NGS analysis*. Refer to Steps 33-34 for how to amplify the sgRNA for NGS. Scale up the number of reactions such that all of the gDNA harvested from the screen is amplified. Each 50 μl reaction can hold up to 2.5 μg of gDNA. Barcoded NGS-Lib-Rev primers enable sequencing of different screening conditions and bioreps on the same sequencing run.

### Troubleshooting

54. *Purification of amplified screening NGS library*. For large-scale PCR purification, we recommend using the Zymo-Spin V with Reservoir. Add 5 volumes of DNA Binding Buffer to the PCR reaction, mix well, and transfer to Zymo-Spin V with Reservoir in a 50 ml Falcon tube. Each Zymo-Spin V column can hold up to 12 ml. Make sure to tighten the connection between the reservoir and the Zymo-Spin V column.

55. Centrifuge at 500 × g for 5 min. Discard the flow-through.

56. Add 2 ml of DNA Wash Buffer and centrifuge at 500 × g for 5 min. Discard the flow-through and repeat for an additional wash.

57. Remove the reservoir from the Zymo-Spin V column and transfer the column to a 1 ml collection tube. In a microcentrifuge, spin at the maximum speed (>12,000 × g) for 1 min to remove residual wash buffer.

58. Transfer the Zymo-Spin V column to a 1.5 ml microcentrifuge tube. Add 150 μl of elution buffer, wait for 1 min, and spin at the maximum speed (>12,000 × g) for 1.5 min to elute the purified PCR reaction.

59. Pool the purified PCR reactions and quantify. Refer to Steps 36-41 for NGS analysis of the sgRNA distribution.

60. *Analysis of screening results with RNAi gene enrichment ranking (RIGER)*. Prior to RIGER analysis, determine the sgRNA fold change due to screening selection. For each biorep of screening experimental or control condition, add a pseudocount of 1 to the NGS read count of each sgRNA and normalize by the total number of NGS read counts for that condition. To obtain the sgRNA fold change, divide the experimental normalized sgRNA count by the control and take the base 2 logarithm.

61. Prepare a RIGER input csv file with the column headers WELL_ID, GENE_ID, and biorep 1, biorep 2, etc from left to right. WELL_ID is a list of sgRNA identification numbers, GENE_ID the genes that the sgRNAs target, and biorep columns the sgRNA fold change. Each row contains a different sgRNA in the library.

62. RIGER is launched through GENE-E from the Broad Institute (http://www.broadinstitute.org/cancer/software/GENE-E/download.html). Start GENE-E and import the input csv file by navigating to File > Import > Ranked Lists. Click the table cell containing the first data row and column as instructed. Launch RIGER by going to Tools > RIGER. Adjust the RIGER settings to the following recommended values:

- Number of permutations: 1,000,000
- Method to convert hairpins to genes: Kolmogorov-Smirnov
- Gene rank order: Positive to negative for positive selection screens; negative to positive for negative selection screens
- Select adjust gene scores to accommodate variation in hairpin set size
- Select hairpins are pre-scored
- Hairpin Id: WELL_ID
- Convert hairpins to: GENE_ID

63. Once the RIGER analysis has completed, export the gene rank dataset. Determine the top candidates genes based on either the overlap or the average ranking between the screening bioreps.

### Validation of candidate genes for screening phenotype o TIMING 3-4 w

64. *Cloning validation sgRNAs into the plasmid backbone of the sgRNA library*. Design top and bottom strand primers for cloning the top 3 sgRNAs for each candidate gene individually into the plasmid backbone of the sgRNA library used for screening according to **Table 4** as we have previously described^57^. Primers for cloning 2 nontargeting sgRNAs (NT1 and NT2) for control are also provided in **Table 4**.

65. Resuspend the top and bottom strand primers to a final concentration of 100 μM. Prepare the following mixture for phosphorylating and annealing the top and bottom strand primers for each validation sgRNA:

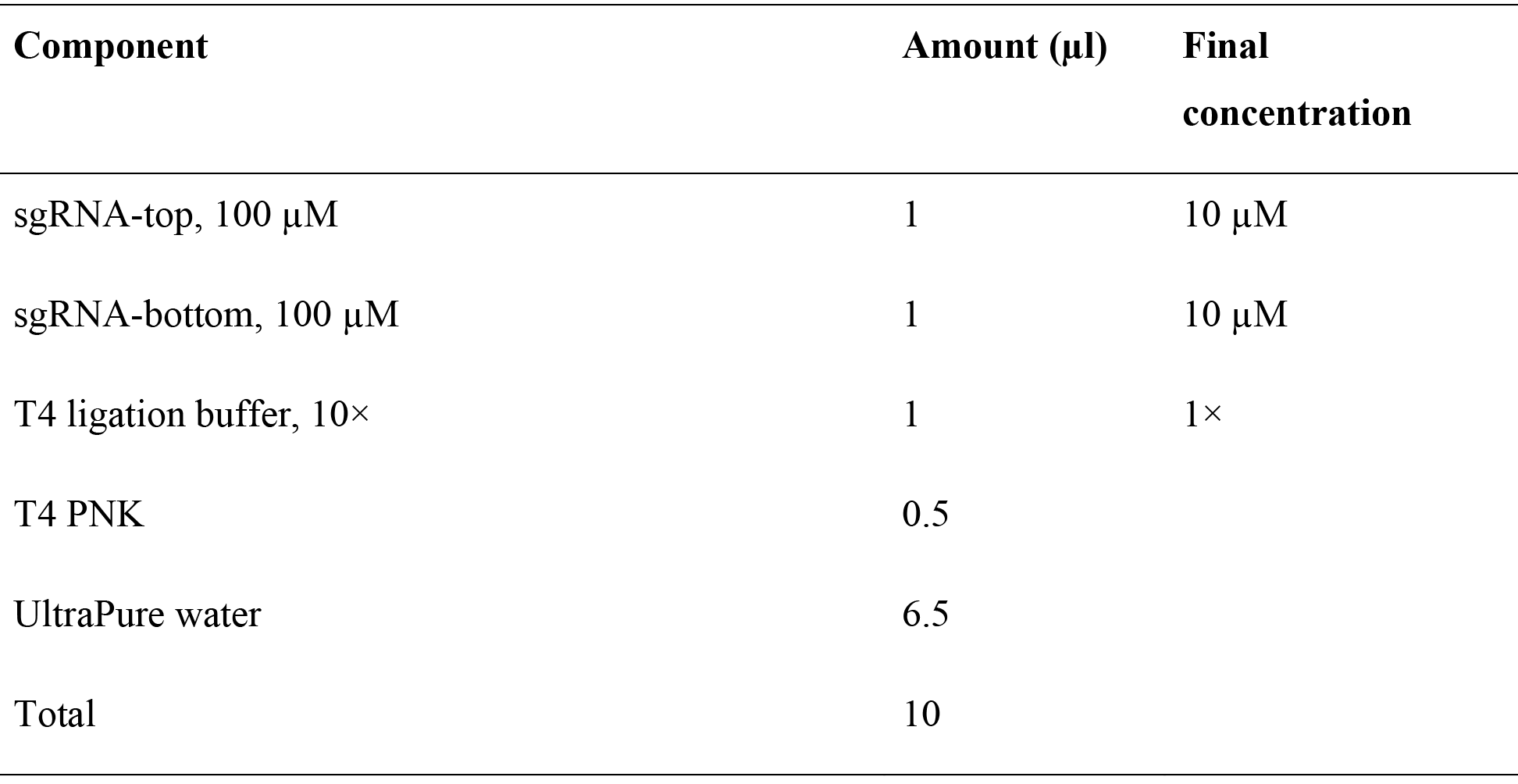

66. Phosphorylate and anneal the primers in a thermocycler by using the following conditions: 37 °C for 30 min; 95 °C for 5 min; ramp down to 25 °C at 5 °C min^-1^.

67. After the annealing reaction is complete, dilute the phosphorylated and annealed oligos 1:10 by adding 90 μl of UltraPure water.

**Pause Point** Annealed oligos can be stored at −20 °C for at least 1 week.

68. Clone the annealed sgRNA inserts into the sgRNA library backbone by setting up a Golden Gate assembly reaction for each sgRNA. We have found that when cloning many sgRNAs, Golden Gate assembly is efficient and offers a high cloning success rate. Mix the following for each sgRNA:

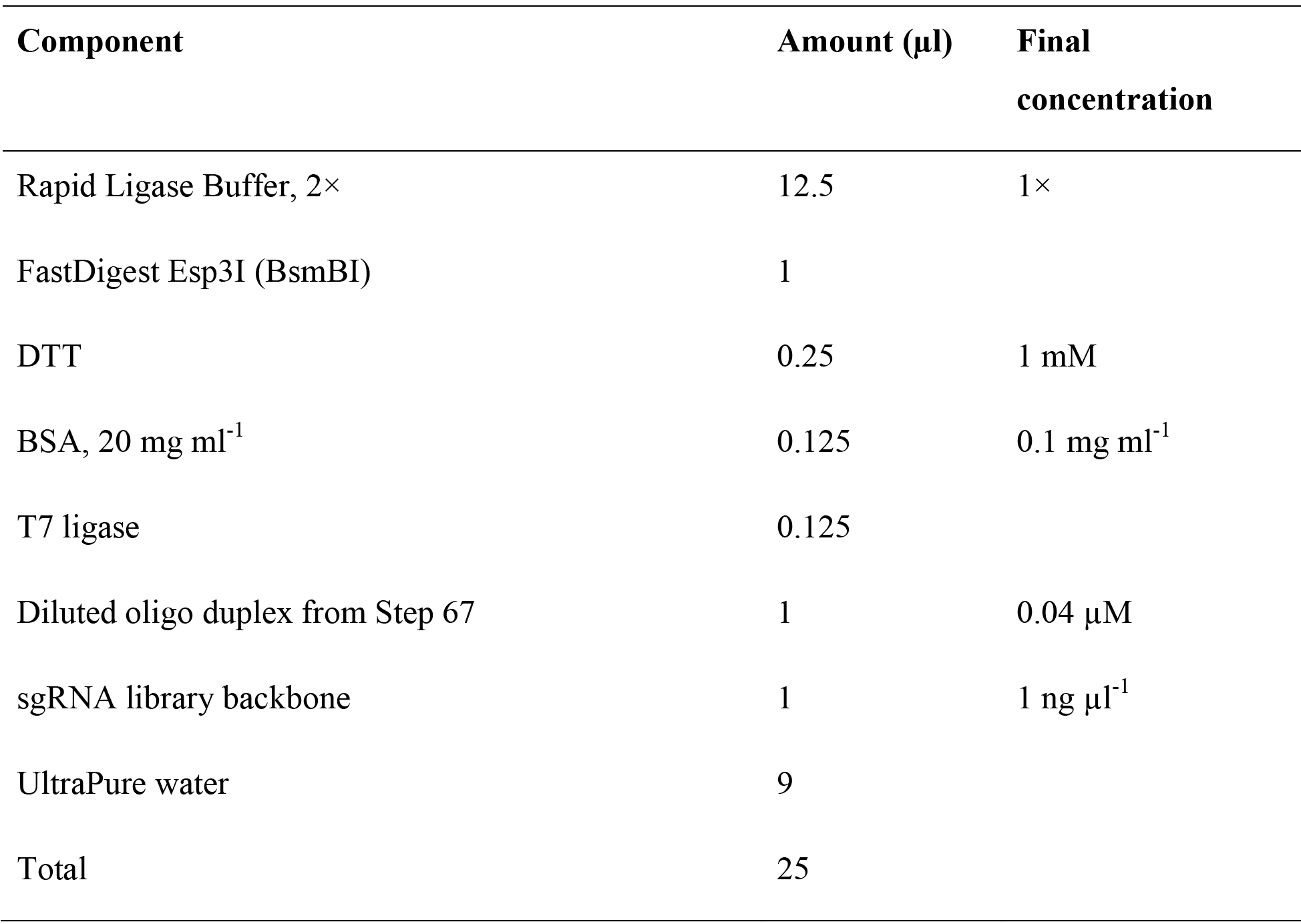

**Critical Step** We recommend using FastDigest BsmBI (Fermentas) as we have had reports of BsmBI from other vendors not working as efficiently in the Golden Gate assembly reaction setup described. It is not necessary to perform a negative control (no insert) Golden Gate assembly reaction as it will always contain colonies and therefore is not a good indicator of cloning success.

69. Perform a Golden Assembly reaction using the following cycling conditions:

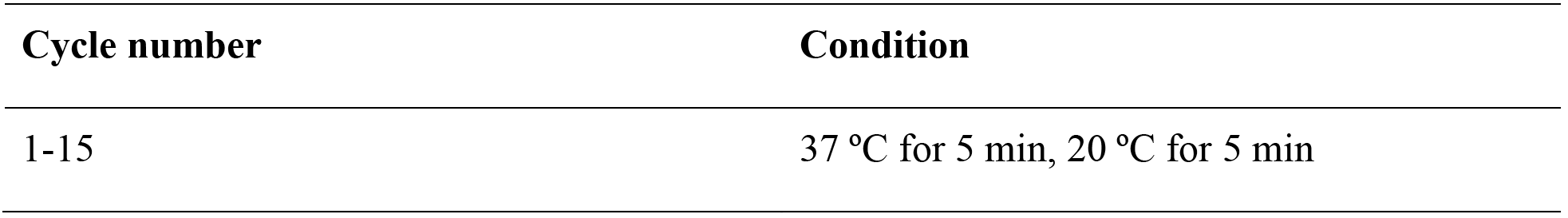

**Pause Point** Completed Golden Gate assembly reactions can be stored at −20 °C for at least 1 week.

*70. Transformation and midiprep*. Transform the Golden Gate assembly reaction into a competent *E. coli* strain, according to the protocol supplied with the cells. We recommend the Stbl3 strain for quick transformation. Thaw the chemically competent Stbl3 cells on ice, add 2 μl of the product from Step 69 into ice-cold Stbl3 cells, and incubate the mixture on ice for 5 min. Heat-shock the mixture at 42 °C for 30 s and return to ice immediately for 2 min. Add 100 μl of SOC medium and plate it onto a standard LB agar plate (100 mm Petri dish, ampicillin). Incubate it overnight at 37 °C.

71. The next day, inspect the plates for colony growth. Typically, there should be tens to hundreds of colonies on each plate.

### Troubleshooting

72. From each plate, pick 1 or 2 colonies for midiprep to check for the correct insertion of sgRNA and for downstream lentivirus production. To prepare a starter culture for midiprep, use a sterile pipette tip to inoculate a single colony into a 3-ml culture of LB medium with 100 μg ml^-1^ ampicillin. Incubate the starter culture and shake it at 37 °C for 4-6 h.

73. Expand each starter culture by transferring the starter culture to 2 separate 25-ml cultures of LB medium with 100 μg ml^-1^ ampicillin in a 50-ml Falcon tube. Remove the cap and seal the top of the tube with AirPore Tape Sheets. Incubate the culture and shake it at 37 °C overnight at >200 rpm.

74. 12-16 h after seeding the starter culture, isolate the plasmids using an endotoxin-free midiprep kit such as the Macherey-Nagel NucleoBond Xtra Midi EF kit according to the manufacturer’s protocol.

**Critical Step** Using an endotoxin-free plasmid purification kit is important for avoiding endotoxicity in virus preparation and mammalian cell culture.

**Pause Point** Midiprepped validation sgRNA constructs can be stored at −20 °C for at least 1 year.

75. *Sequence validation of sgRNA cloning*. Verify the correct insertion of the validation sgRNAs by sequencing from the U6 promoter using the U6-fwd primer. Compare the sequencing results to the sgRNA library plasmid sequence to check that the 20-nt sgRNA target sequence is properly inserted between the U6 promoter and the remainder of the sgRNA scaffold.

### Troubleshooting

76. *Generation of validation cell lines*. Prepare lentivirus for validation by scaling down the lentivirus production in Steps 42-46 to T25 flasks or 2 wells of a 6-well plate. Filter the lentivirus supernatant using 5-ml syringes and 0.45 μm filters.

77. Titer the lentivirus according to Step 47. If preparing multiple validation sgRNA lentivirus in the same plasmid backbone at the same time, titer lentivirus from 2-3 different sgRNAs and extend the average titer to the rest of the lentivirus.

78. Similar to during screening, transduce either naive cells or MS2-p65-HSF1-expressing cells for knockout or activation screening with validation sgRNA lentivirus at an MOI < 0.5 according to Steps 48-50. For knockout validation, select for 7 d to allow for sufficient time for indel saturation.

79. *Validation of candidate genes for screening phenotype*. Once the antibiotic selection for validation cell lines is complete, verify the screening phenotype. In addition, determine the indel rate for knockout screens (Steps 80-96) or fold activation for activation screens (Steps 97-108).

80. *Indel rate analysis for validating a knockout screen*. We describe a two-step PCR for NGS in which the first step uses custom primers to amplify the genomic region of interest and the second step uses universal, barcoded primers for multiplexed sequencing of up to 96 different samples in the same NGS run. For each validation sgRNA, design custom round 1 NGS primers (NGS-indel-R1) that amplify the 100-300bp region centered around the sgRNA cut site according to **Table 5**. It is important to design primers situated at least 50 bp from the target cleavage site to allow for the detection of longer indels. Aim for an annealing temperature of 60 °C and check for potential off-target sites using Primer-BLAST. If necessary, include a 1-10bp staggered region to increase the diversity of the library.

**Table 5.**
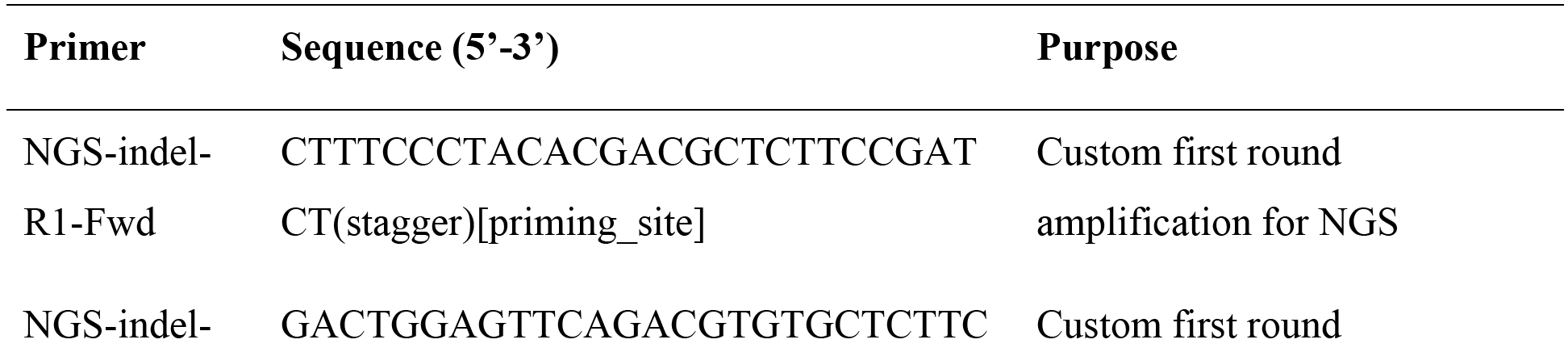
Primers for amplifying target sites to determine percentage of indels by NGS. Custom first round primers amplify the target region and second round universal, barcoded primers amplify the first round products for multiplexed NGS.

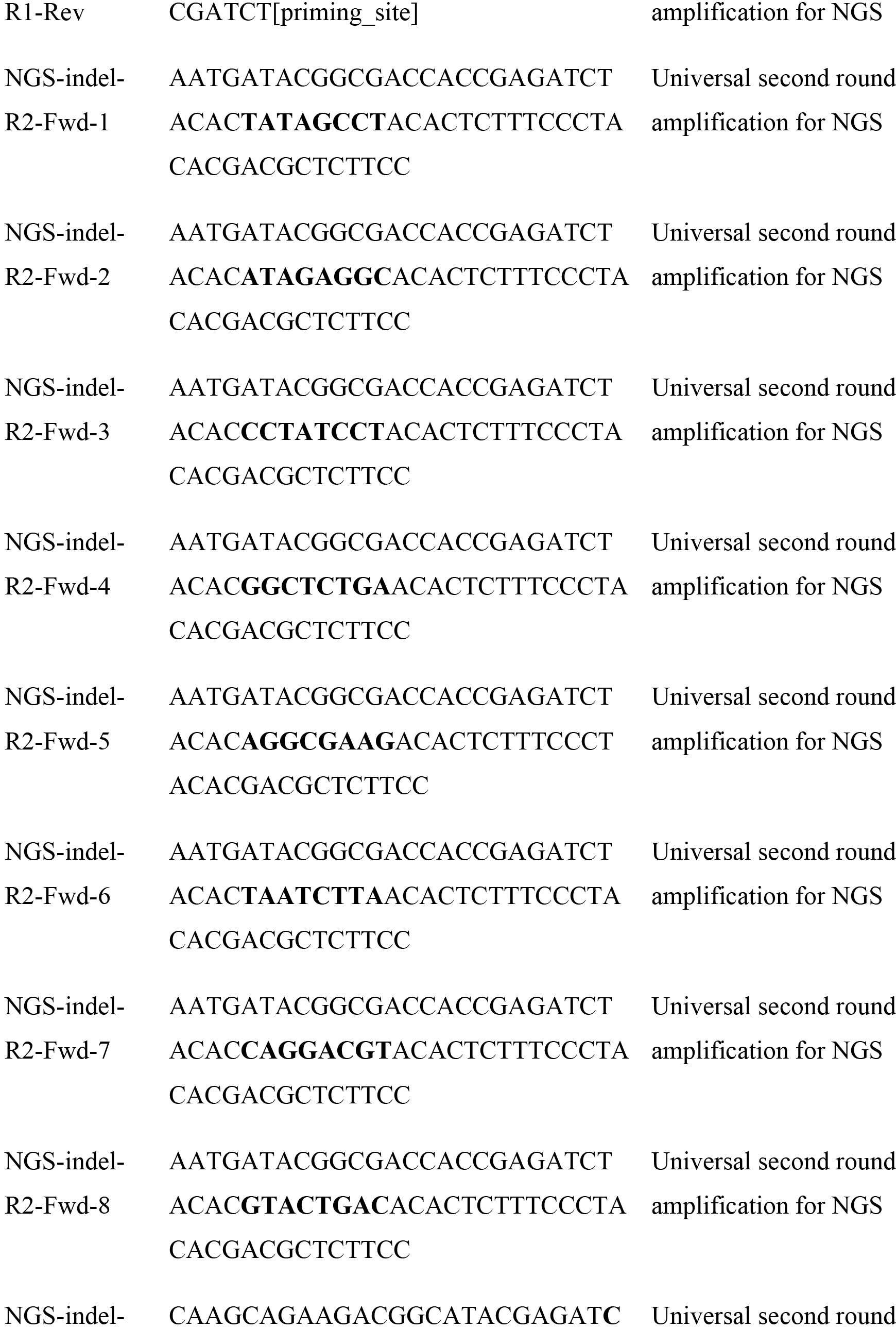

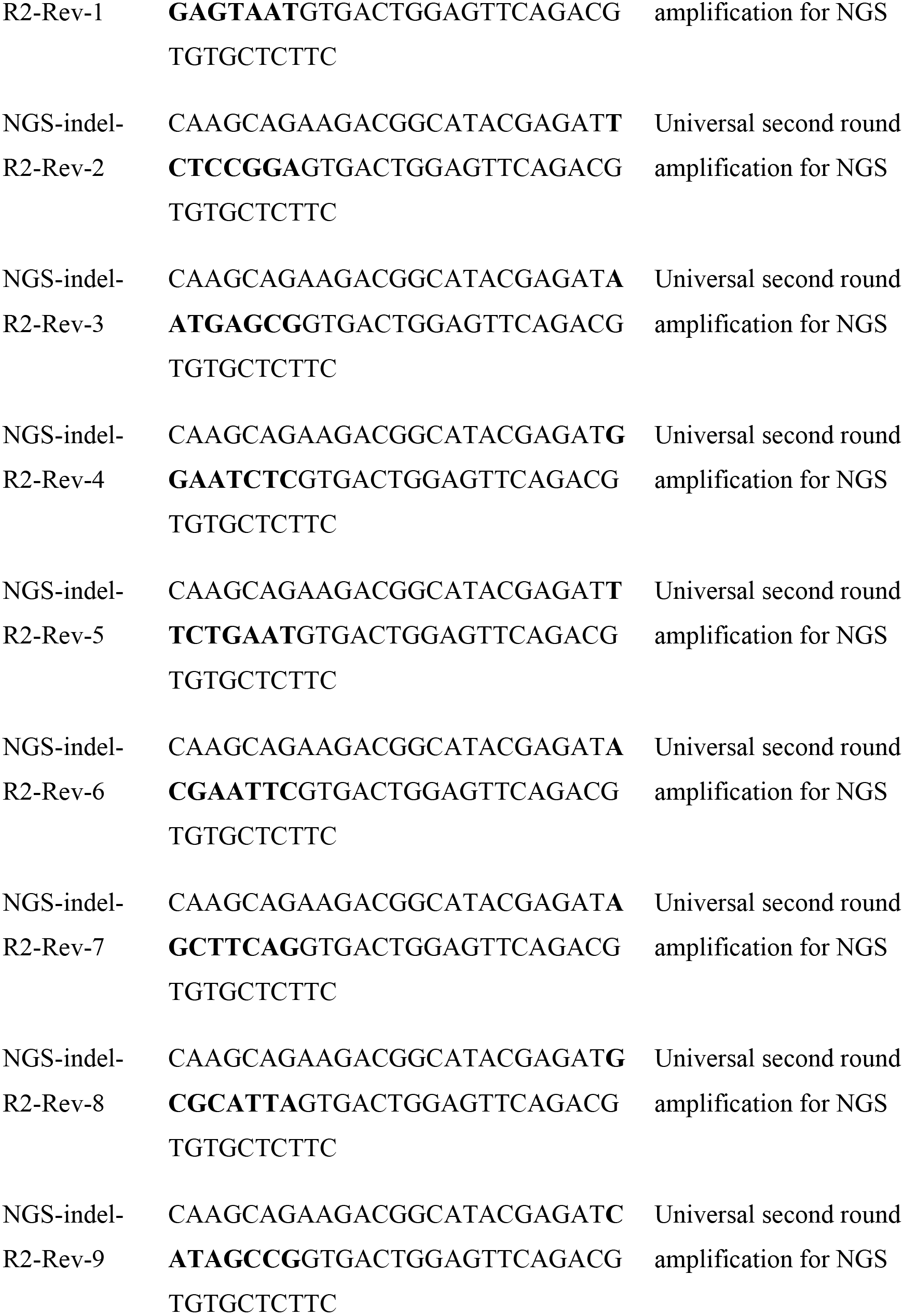

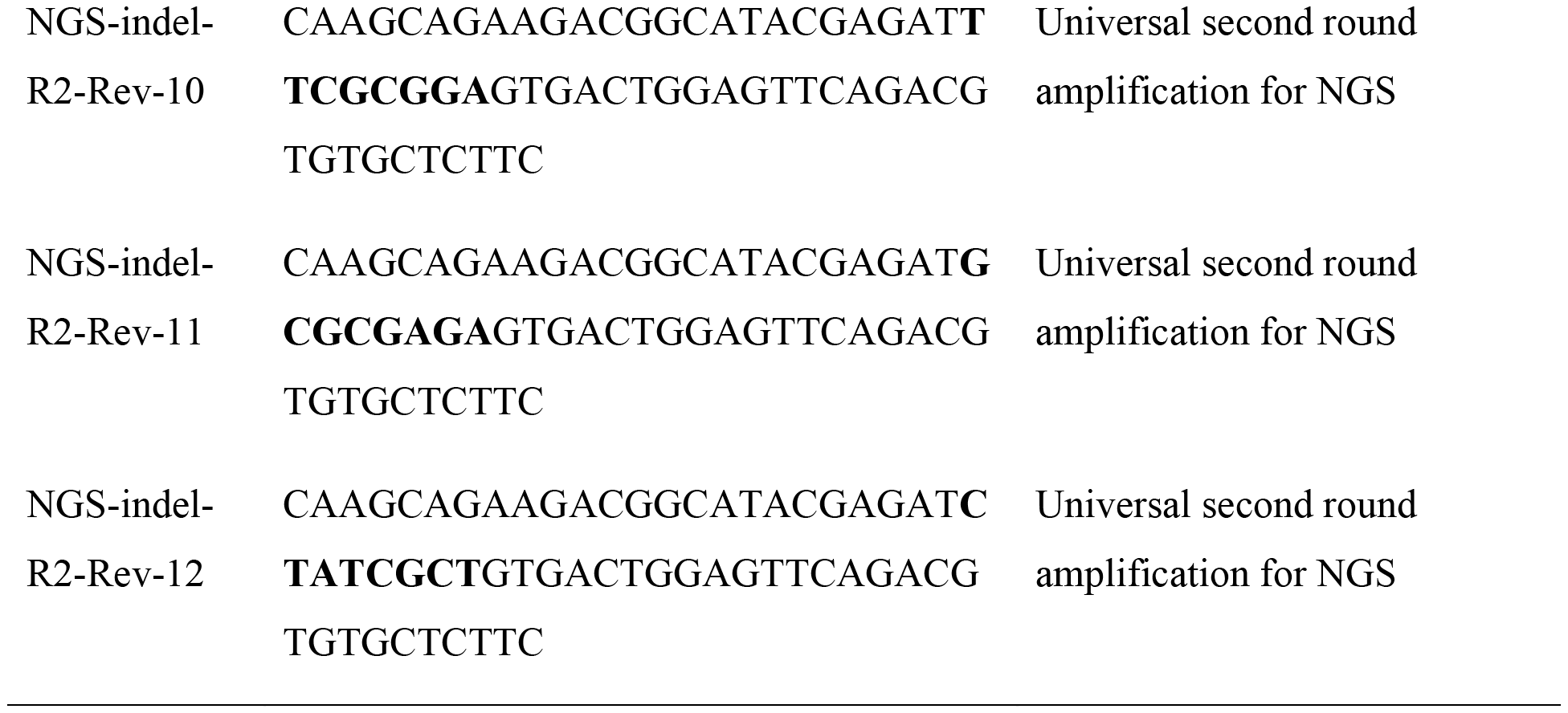

**Table 5.**
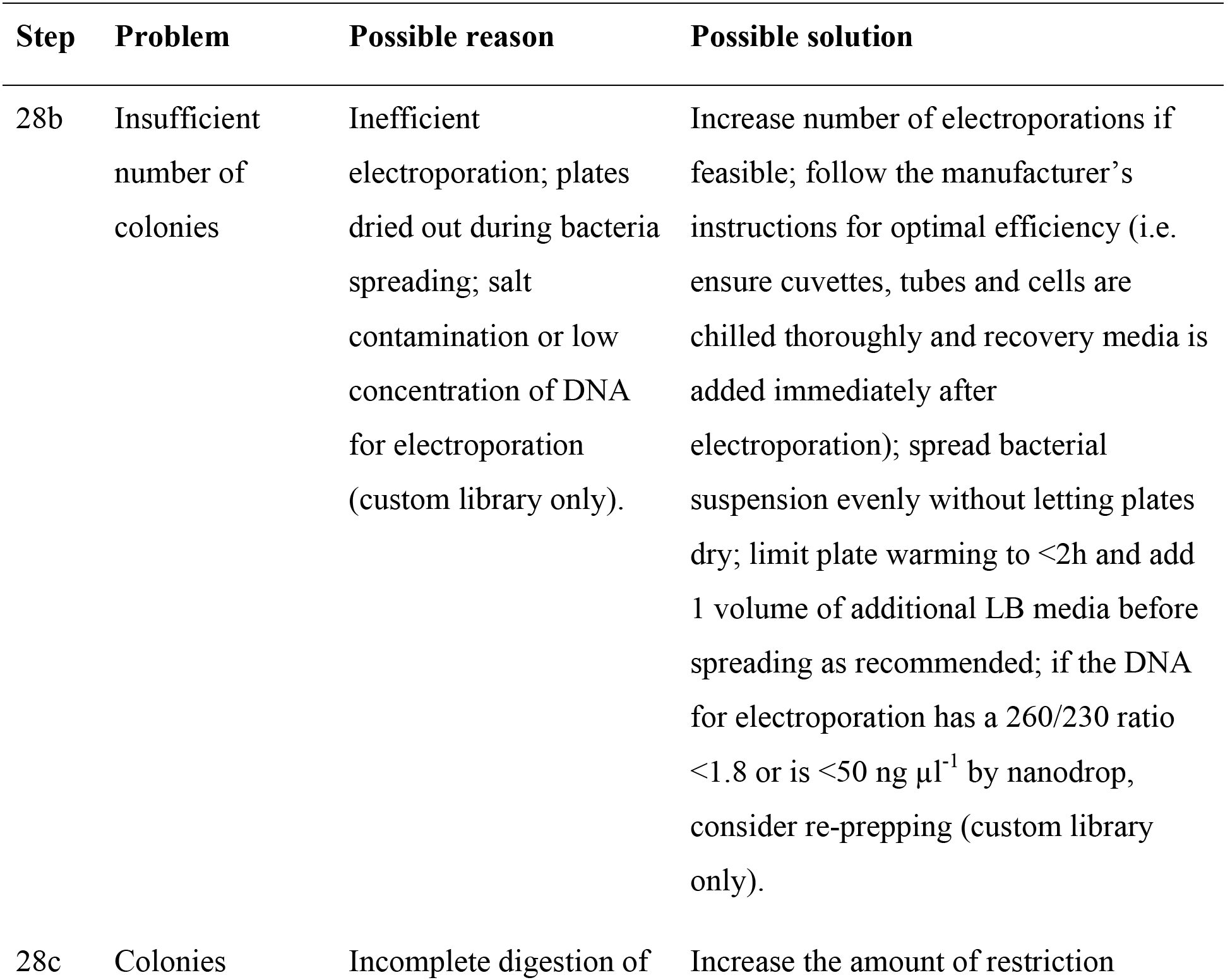
Troubleshooting table.

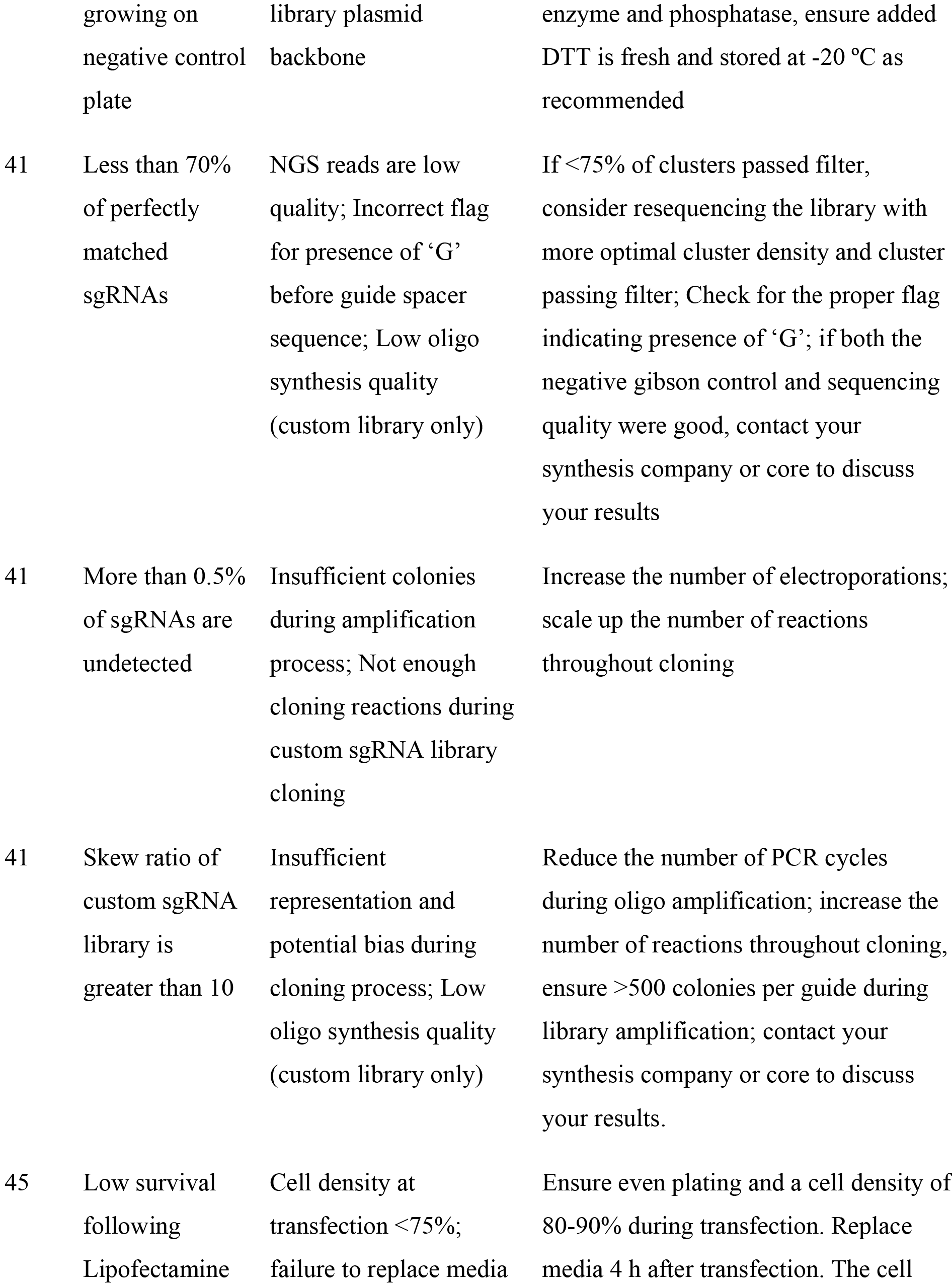

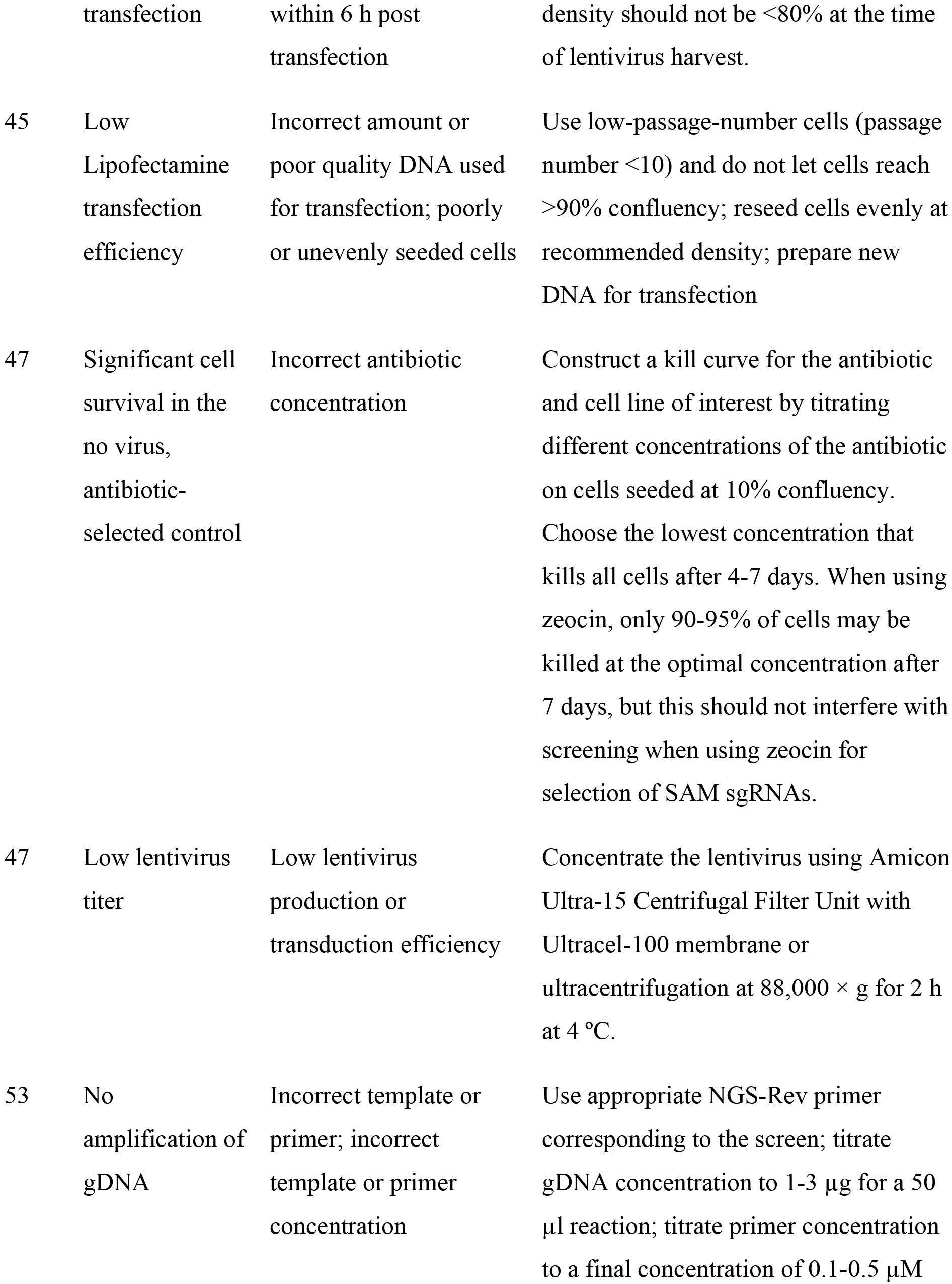

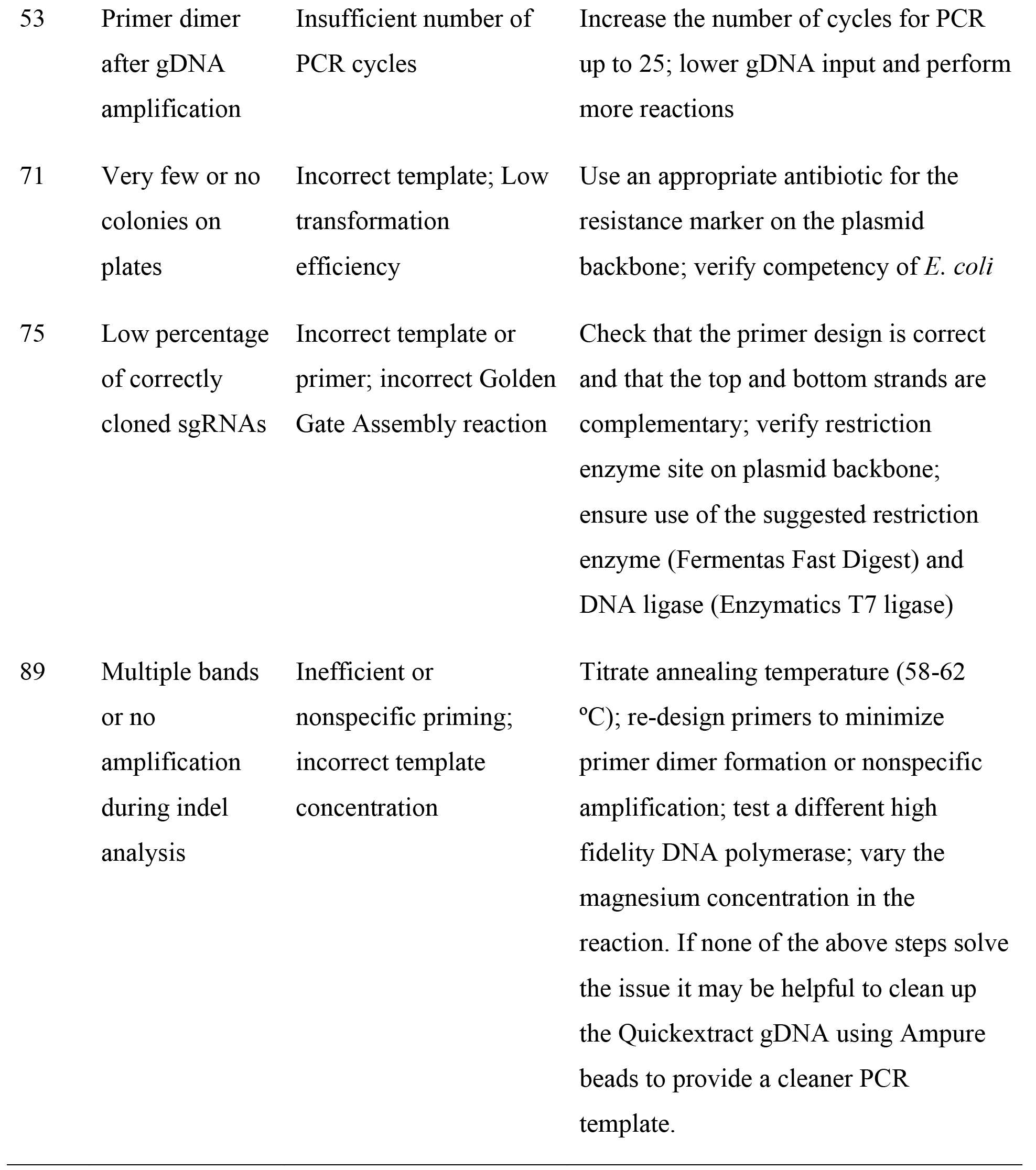

81. *Harvest gDNA from validation cell lines*. Seed the validation cells at a density of 60% confluency with 3 bioreps in a 96-well clear bottom black tissue culture plate.

82. 1 d after seeding, when the cells have reached confluency, aspirate the media and add 50 μl of QuickExtract DNA Extraction Solution. Incubate at room temperature for 2-3 min.

83. Scrape the cells with a pipette tip, mix thoroughly by pipetting up and down, and transfer the mixture to a 96-well PCR plate.

84. Extract the genomic DNA by running the following cycling conditions:

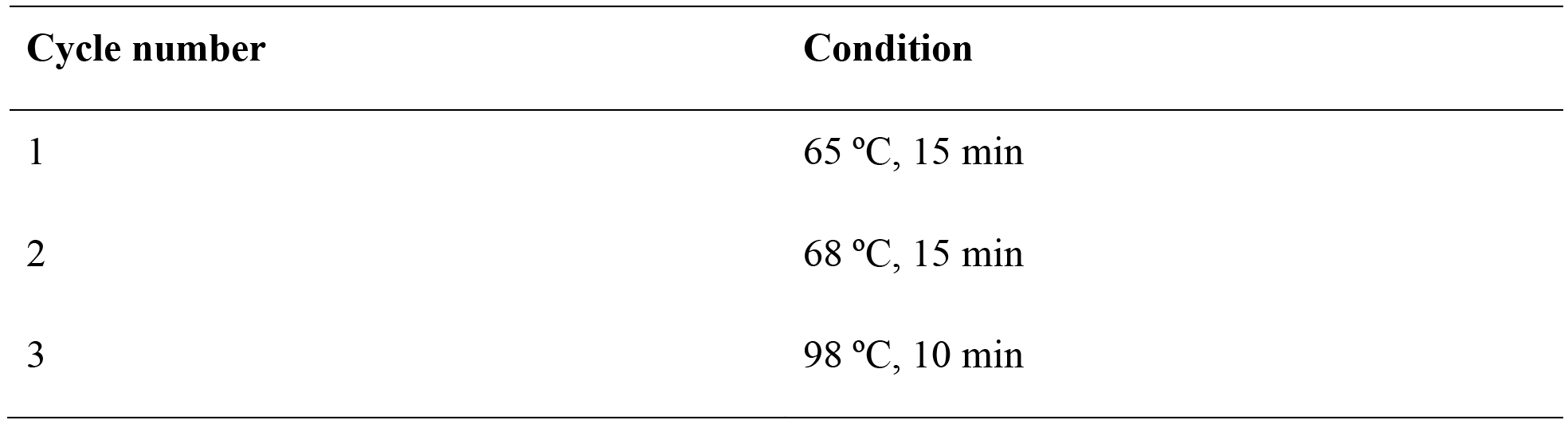

**Pause Point** Extracted genomic DNA can be stored at −20 °C for up to several months.

85. *First round PCR for indel analysis by NGS*. Amplify the respective target regions for each validation and control cell line by using custom NGS-indel-R1 primers (**Table 5**) in the following reaction:

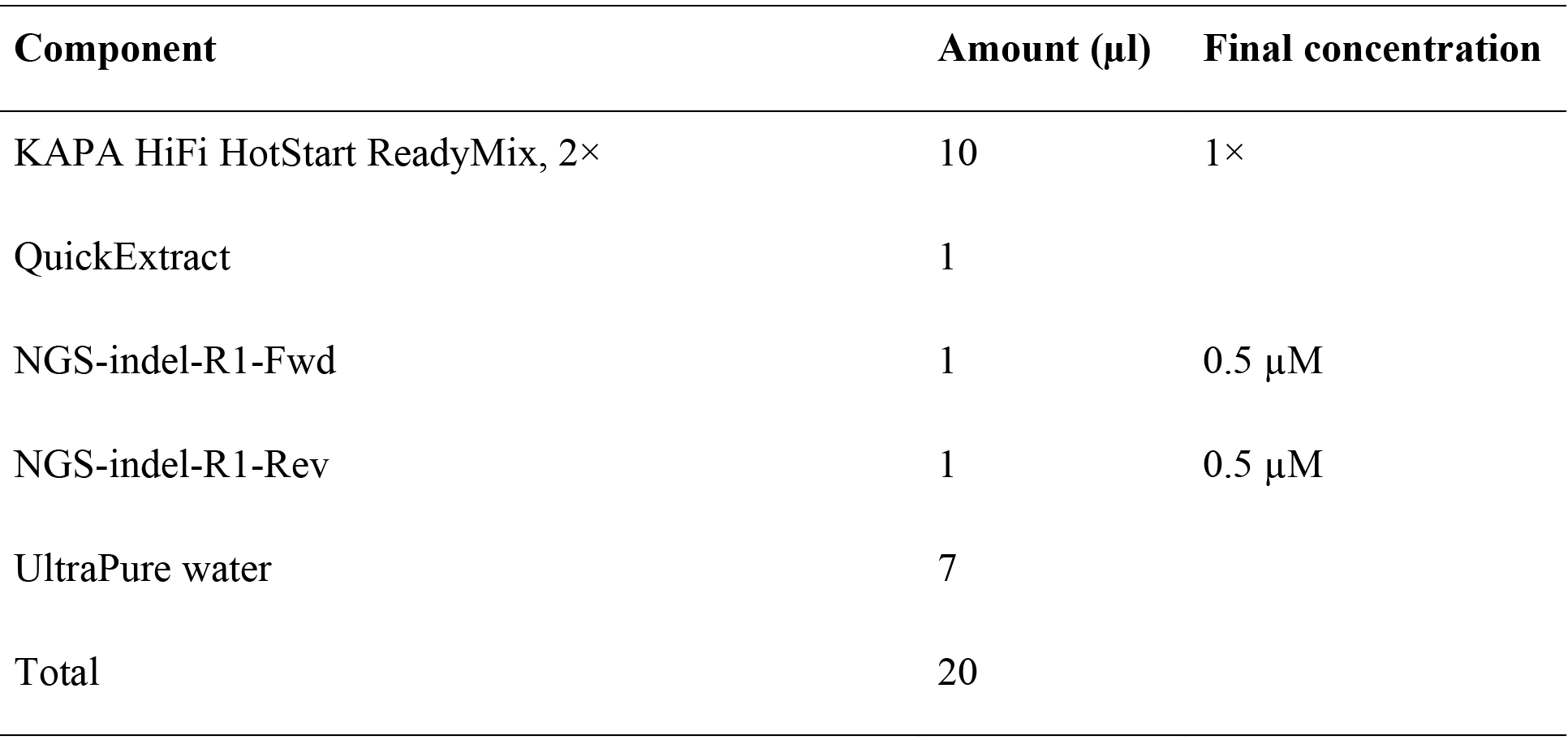

**Critical Step** To minimize error in amplifying sgRNAs, it is important to use a high-fidelity polymerase. Other high-fidelity polymerases, such as PfuUltra II (Agilent) or NEBNext (New England BioLabs), may be used as a substitute.

86. Perform a PCR with the following cycling conditions:

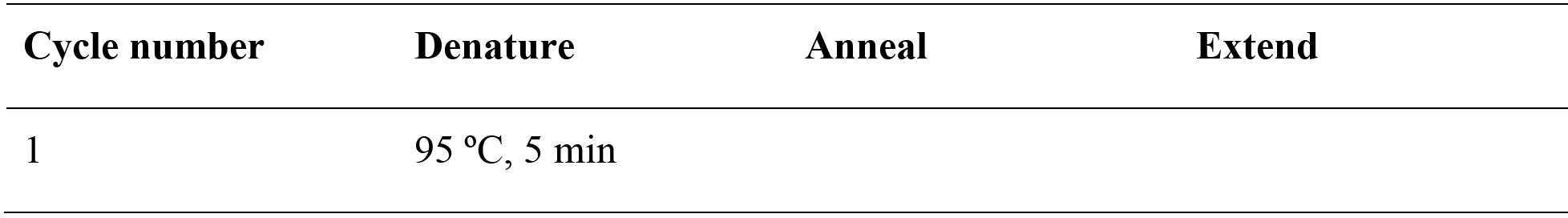

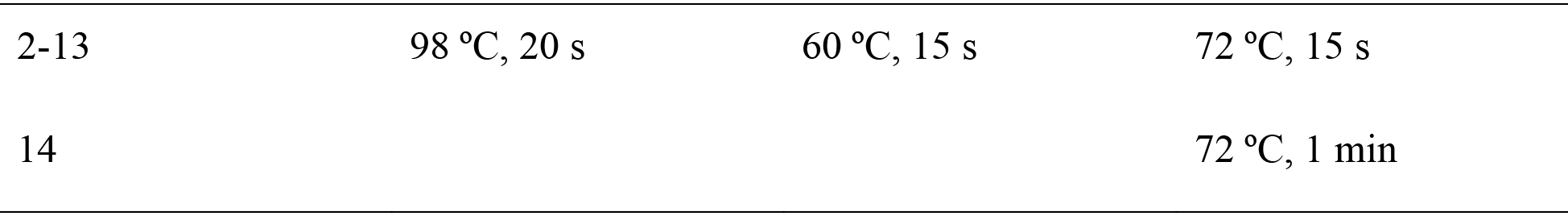

87. *Second round PCR for indel analysis by NGS*. Barcode the first round PCR for NGS by amplifying the product with different NGS-indel-R2 primers (**Table 5**) in the following reaction:

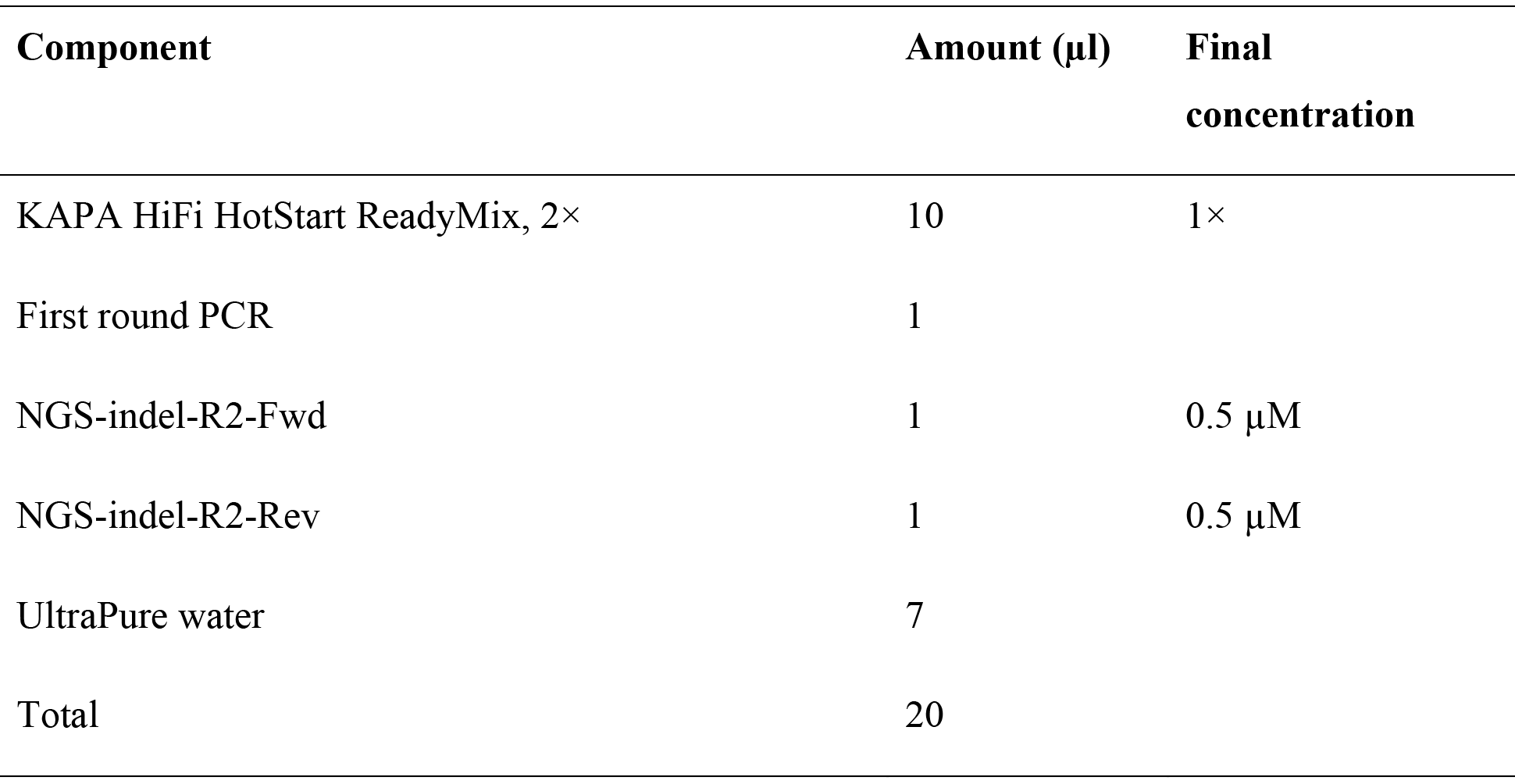

88. Perform a PCR using the same cycling conditions as described in Step 76.

89. After the reaction is complete, run 1 μl of each amplified target on a 2% (wt/vol) agarose gel to verify successful amplification of a single product at the appropriate size.

### Troubleshooting

90. Pool the PCR products and purify the pooled product using the QIAquick PCR purification kit.

91. Gel extract the appropriate sizes using the QIAquick gel extraction kit according to the manufacturer’s directions.

**Pause Point** Gel-extracted product can be stored at −20 °C for several months.

92. Sequence gel-extracted samples on the Illumina MiSeq according to the Illumina user manual with 260 cycles of read 1, 8 cycles of index 1, and 8 cycles of index 2. We recommend aiming for >10,000 reads per sgRNA.

93. *Indel analysis of validation sgRNAs with calculateindel.py*. Install biopython (http://biopython.org/DIST/docs/install/Installation.html) and SciPy (https://www.scipy.org/install.html). Construct a sample sheet with each line corresponding to a separate file. The structure of each line should be: <sample name>,<fasta/fastq file name>,<guide sequence>,<PCR target amplicon>,<Experimental or Control>. The last column is only required when performing the maximum likelihood estimate (MLE) correction. When MLE is performed, control samples which reflect the background indel rate should be labeled “Control” and experimental samples should be labeled “Experimental”.

94. If processing all files with a single command, run python calculate_indel.py, with following optional parameters:

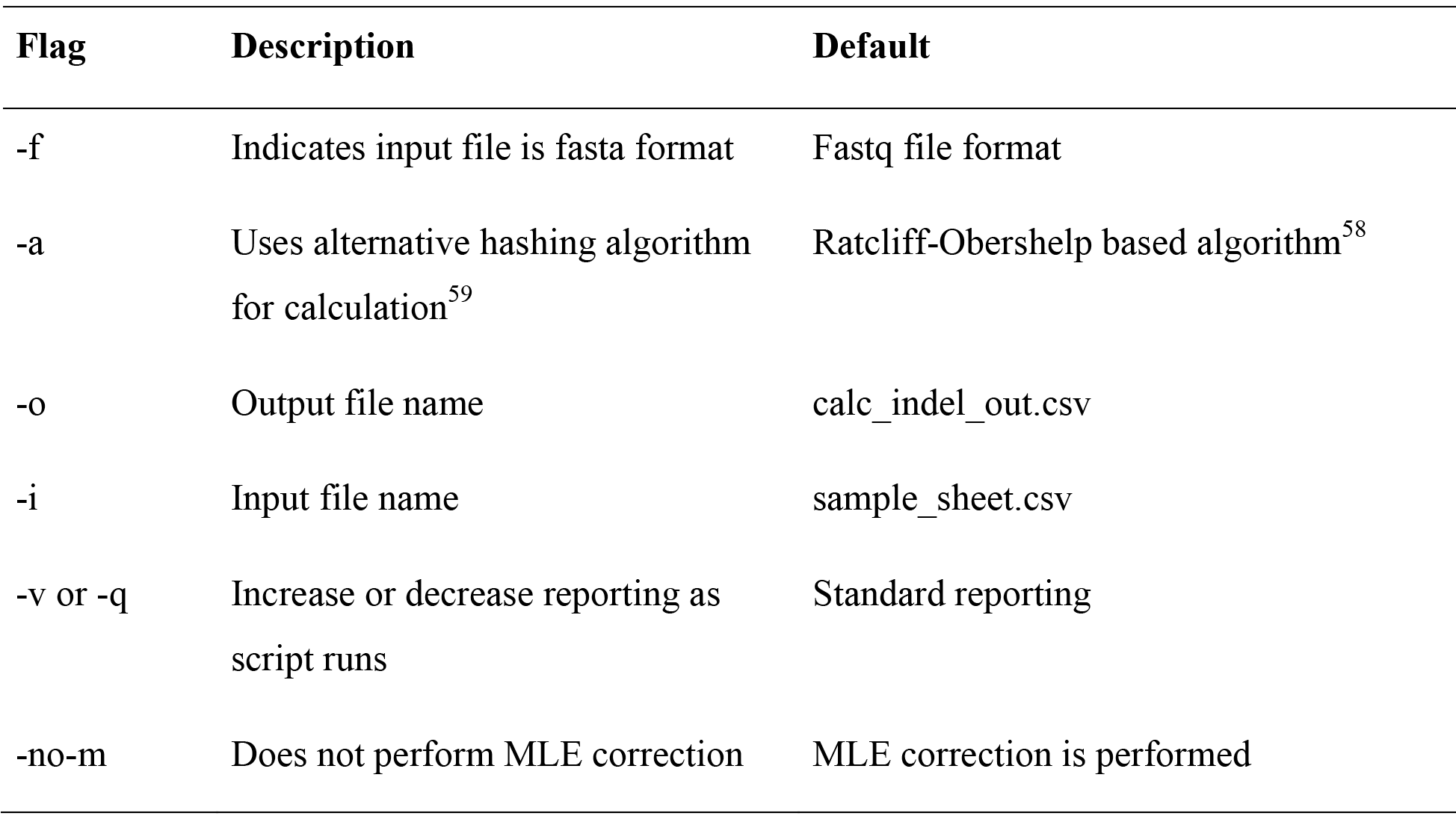

95. Place all files in the same folder before running calculate_indel.py. For processing individual samples, such as in the case of parallelization, run python calculate_indel.py-sample <sample name>. This will produce a file <sample name>_out.csv, which can be combined by calling python calculate_indel.py --combine

96. After running calculate_indel.py, calculated indels will be in the output file, which also contains counts of reads that matched perfectly, failed to align, or were rejected due to quality, or had miscalled bases/replacements. There will also be three columns corresponding to the MLE corrected indel rate, as well as the upper and lower bounds for the 95% confidence interval of indels

97. *Determine the fold activation for validating an activation screen*. Prepare cells by seeding the validation cells at a density of 60% confluency with 4 bioreps for each validation cell line in a 96-well poly-d-lysine coated tissue culture plate.

98. *Reverse transcription to cDNA*. When the cells are confluent approximately 1 d after seeding, prepare the following reagents for each well:

- Complete RNA lysis buffer:

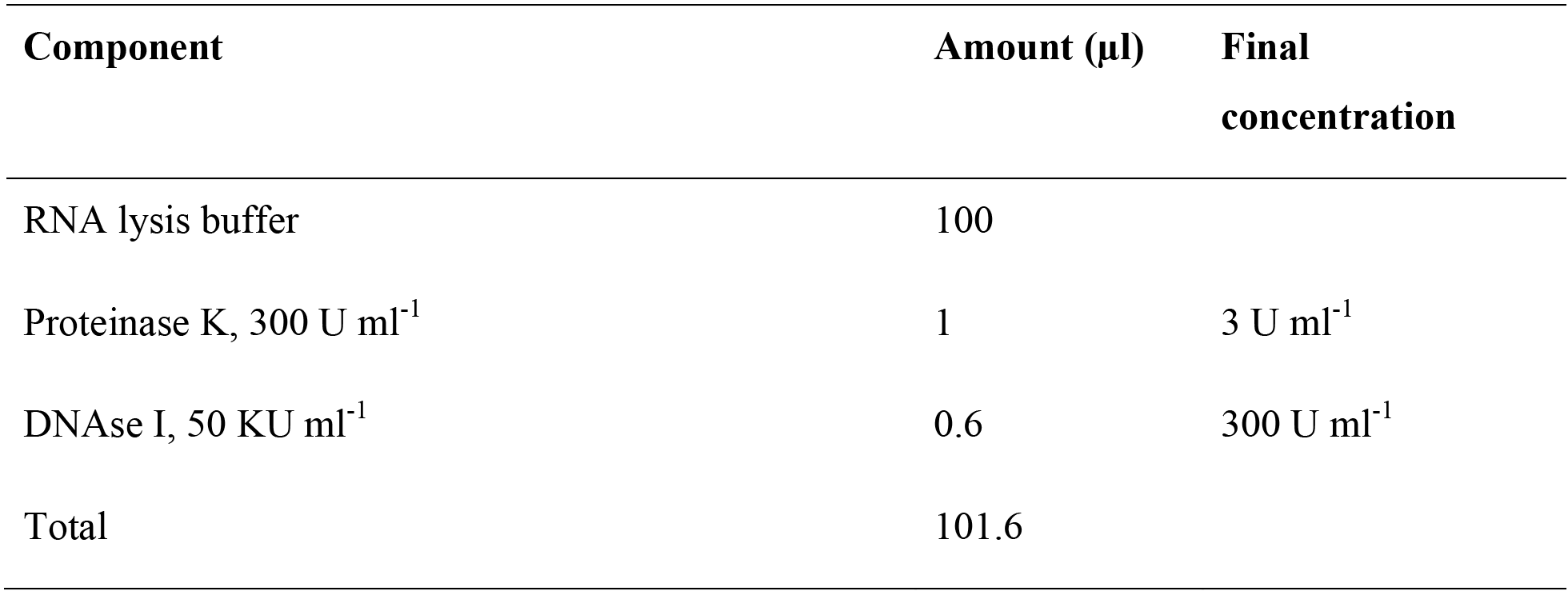
- Reverse Transcription Mix:

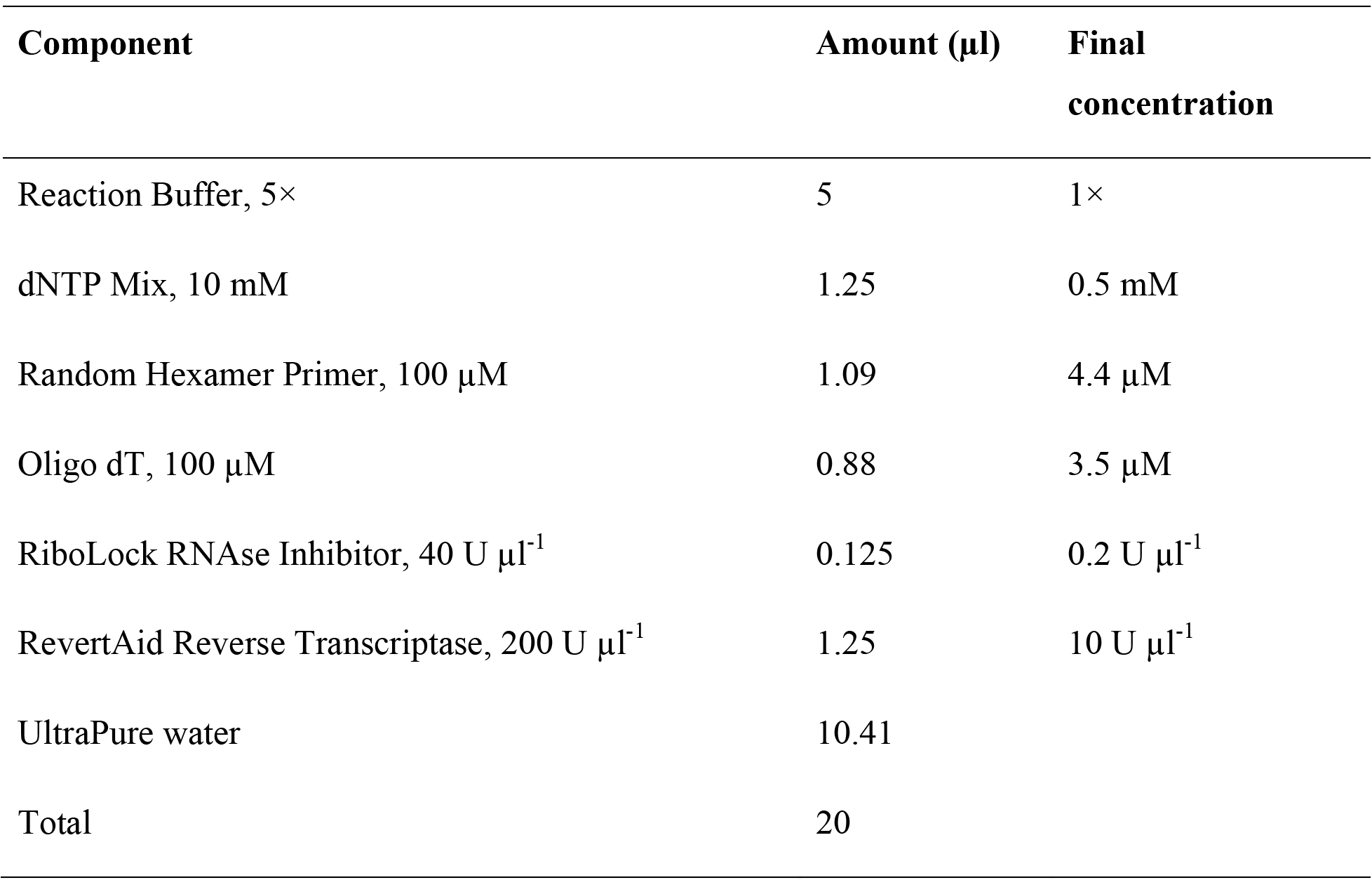

Except for Oligo dT, all components can be found in the Thermo RevertAid RT Reverse Transcription kit.

**Critical Step** Make sure all reagents are RNAse free and take proper precautions when working with RNA.

99. Aliquot 20 μl of the Reverse Transcription Mix into each well of a 96-well PCR plate. Thaw the RNA lysis stop solution, prepare cold DPBS, and keep all reagents except for the complete RNA lysis buffer on ice until needed.

100. Aspirate the media from each well of the 96-well poly-d-lysine tissue culture plate, wash with 100 μl of cold DPBS, and add 100 μl of room temperature complete RNA lysis buffer. Incubate at room temperature while mixing thoroughly for 6-12 min to lyse the cells.

**Critical Step** It is important to limit the lysis time to less than 12 min to prevent RNA degradation.

101. Transfer 30 μl of the cell lysate to a new 96-well PCR plate. Add 3 μl of RNA lysis stop solution to terminate lysis and mix thoroughly. The cell lysate with RNA lysis stop solution can be stored at −20 °C for additional reverse transcription reactions.

102. Then, add 5 μl of the cell lysate with RNA lysis stop solution to the Reverse Transcription Mix for a total volume of 25 μl and mix thoroughly.

103. Reverse transcribe the harvested RNA to cDNA with the following cycling conditions:

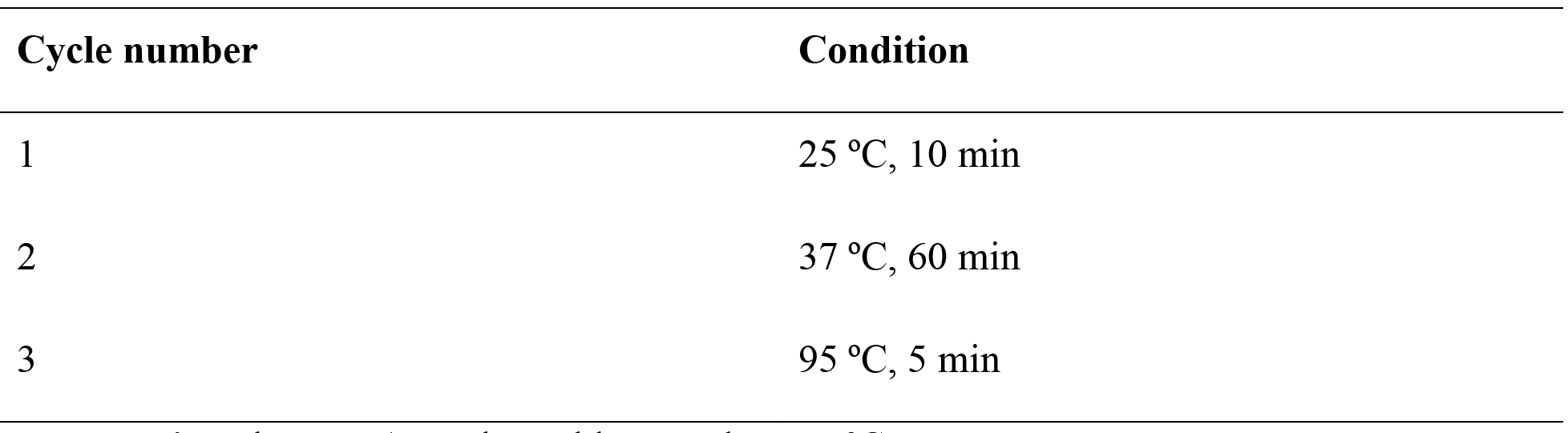

**Pause Point** The cDNA can be stably stored at −20 °C.

104. *Perform a TaqMan qPCR for fold activation analysis*. Thermo Fisher Scientific provides design ready TaqMan Gene Expression Assays for candidate genes as well as for endogenous control genes such as GAPDH or ACTB. Make sure that the experimental and control gene expression assays have different probe dyes (i.e. VIC and FAM dyes) that allow for running in the same reaction.

105. Prepare the following qPCR master mix per reverse transcription reaction. We recommend pre-mixing the TaqMan Fast Advanced Mastermix and gene expression assays for all samples with the same target gene first.

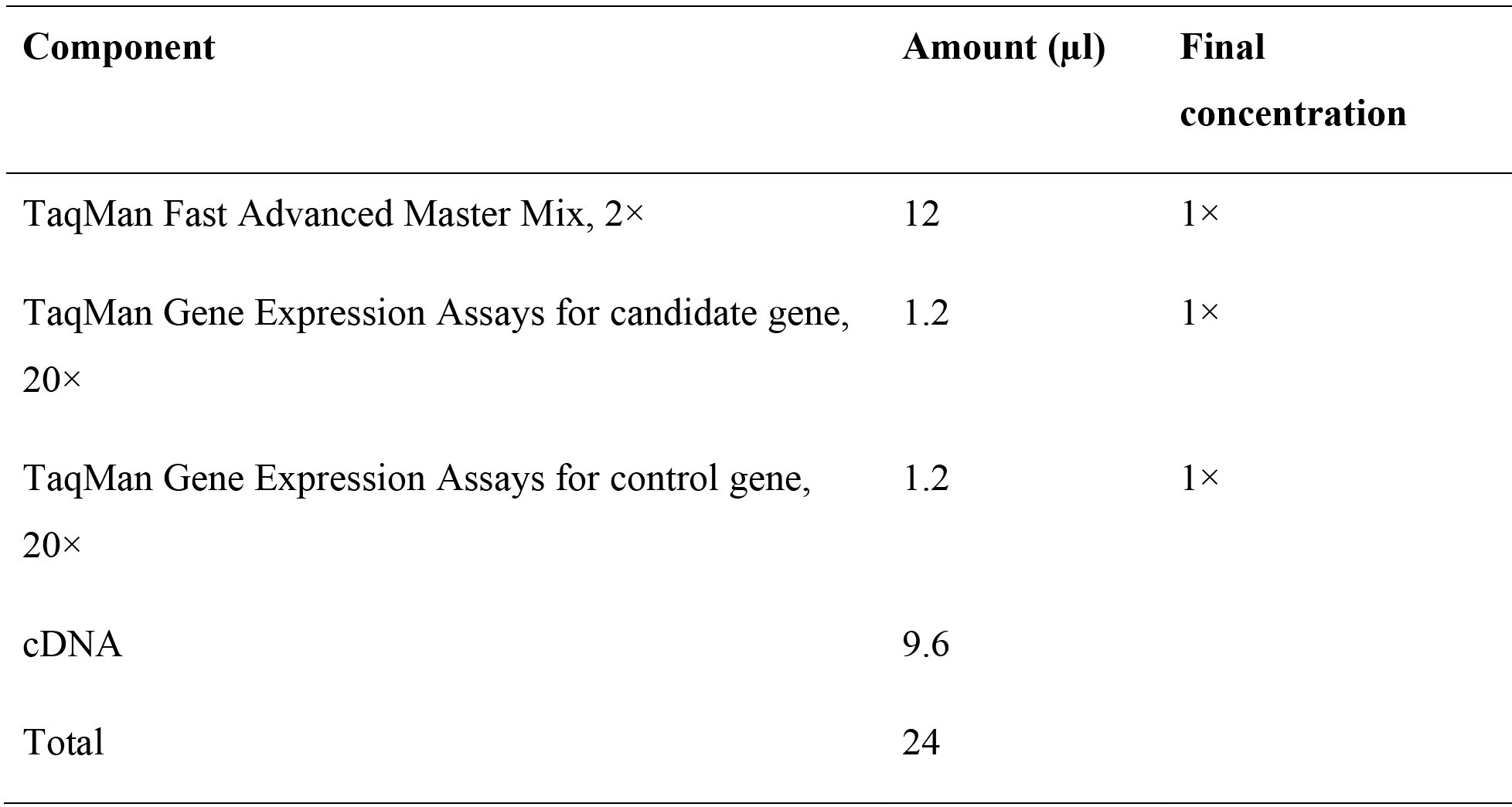

106. Aliquot 4 × 5 μ of the qPCR master mix into a 384-well optical plate for technical replicates.

107. Perform a qPCR with the following cycling conditions:

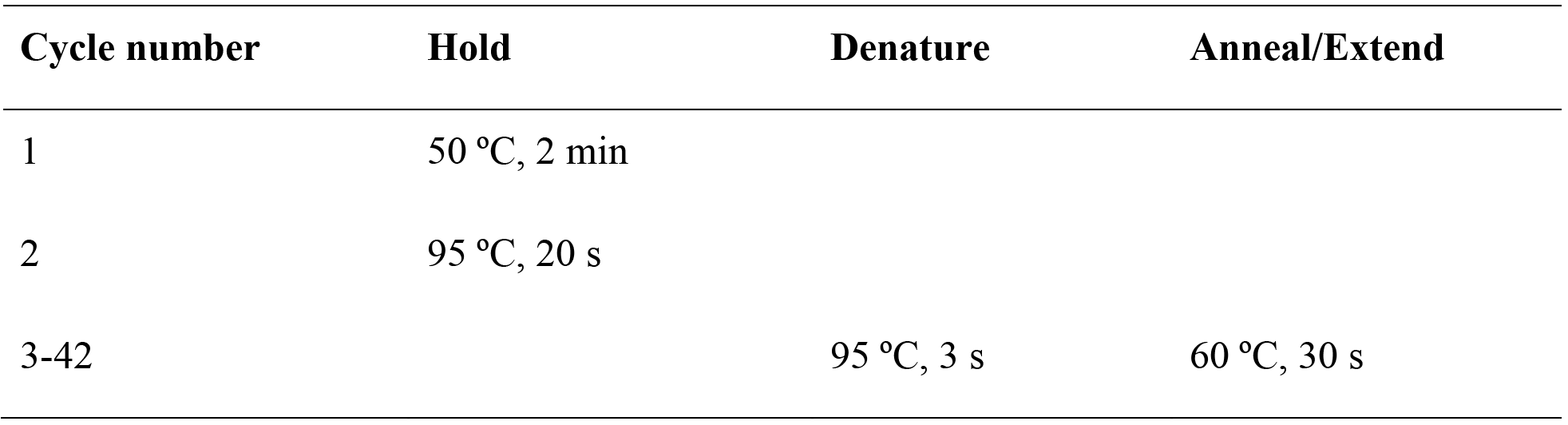

108. Once the qPCR is complete, calculate the candidate gene expression fold change relative to control using the ddCt method according to the instrument manufacturer’s protocol.

109. *Additional steps for combining candidate genes from knockout and activation screens using dead sgRNAs (dRNAs)*. dRNAs are sgRNAs with 14- or 15-nt spacer sequences, which are truncated versions of the standard sgRNAs with 20-nucleotide spacer sequences, that are still capable of binding DNA^60, 61^. dRNAs are considered catalytically ‘dead’ because they can guide wild-type Cas9 without inducing doublestranded breaks. Adding MS2 binding loops to the dRNA backbone allows for wild-type Cas9 to activate transcription without cleavage. To achieve simultaneous knockout and activation, refer to Steps 42-49 to generate a cell line that stably expresses wild-type Cas9 and MS2-p65-HSF1.

110. Transduce the Cas9-and MS2-p65-HSF1-expressing cell line with a standard sgRNA for knocking out a candidate gene and a 14-nt dRNA with MS2 binding loops for activating a second candidate gene according to Steps 64-78.

111. Verify the screening phenotype, indel percentage, and fold activation according to Steps 79-108.

### Troubleshooting

Troubleshooting advice can be found in **Table 5**.

### Timing

Steps 1-20, designing and cloning a targeted screen: 3 d

Steps 21-32, amplification of pooled sgRNA library: 2 d

Steps 33-41, next-generation sequencing of the amplified sgRNA library: 2-3 d

Steps 42-47, lentivirus production and titer: 8-10 d

Steps 48-51, lentiviral transduction and screening: 3-6 w

Steps 52-63, harvest genomic DNA for screening analysis: 3-4 d

Steps 64-111, validation of candidate genes: 3-4 w

## ANTICIPATED RESULTS

As a reference for screening results, we provide data from genome-scale knockout and transcriptional activation screens for genes that confer BRAF inhibitor vemurafenib (PLX) resistance in a BRAF^V600E^ (A375) cell line. After applying vemurafenib selection, the sgRNA library distribution, which is measured by NGS, in the experimental condition is more skewed than the baseline and vehicle control conditions, with some sgRNAs
enriched and others depleted (**Fig. 4a,b**). SgRNAs targeting genes involved in vemurafenib resistance are enriched because they provide a proliferation advantage upon vemurafenib treatment. RIGER analysis of enriched sgRNAs in the vemurafenib condition relative to the control identified several candidate genes responsible for resistance. Each candidate gene has multiple significantly enriched sgRNAs (**Fig. 4c, d**) and *P* values that are significantly lower than the rest of the genes (**Fig. 4e, f**). To clarify potential points of confusion when performing genome-scale screens using CRISPR-Cas9, we have compiled a list of most-frequently asked questions from our web-based CRISPR forum (discuss.genome-engineering.org) (**Box 6**). (**Fig. 4**)

## Acknowledgements

We would like to thank O. Shalem, D. A. Scott, and P. D. Hsu for helpful discussions and insights; R. Belliveau for overall research support; R. Macrae for critical reading of the manuscript; and the entire Zhang laboratory for support and advice. O. A.A. is supported by a Paul and Daisy Soros Fellowship, a Friends of the McGovern Institute Fellowship, and the Poitras Center for Affective Disorders. J.S.G. is supported by a D.O.E. Computational Science Graduate Fellowship. F.Z. is supported by the NIH through NIMH (5DP1-MH100706 and 1R01-MH110049) and NIDDK (5R01DK097768-03), the New York Stem Cell, Simons, Paul G. Allen Family, and Vallee Foundations; and David R. Cheng, Tom Harriman, and B. Metcalfe. F.Z. is a New York Stem Cell Foundation Robertson Investigator. Reagents are available through Addgene; support forums and computational tools are available via the Zhang lab website (http://www.genome-engineering.org).

## Author contributions

J.J., S.K., J.S.G., O.O.A., R.J.P., M.D.B., N.S., and F.Z. designed and performed the experiments. J.J., S.K., and F.Z. wrote the manuscript with help from all authors.

## Boxes

### Box 1: Different types of screening selection

Before setting up the screen, it is important to determine the type of screening selection based on the phenotype of interest and available selection pressures for the screen. Positive selection screens rely on enrichment of sgRNAs for genetic perturbations that produce the screening phenotype as a result of cell proliferation. These typically have the highest signal-to-noise ratio compared to the other types of screens because the number of phenotypically relevant sgRNAs increases relative to the rest of the sgRNAs. On the other hand, negative selection screens involve depletion of sgRNAs that correspond to the phenotype due to cell death. However, for a large number of screens the phenotype of interest will not result in cell proliferation or cell death and thus the phenotypically relevant sgRNAs are not enriched or depleted. For these phenotypes, the screen may be read out by changes in protein expression using either endogenous-tagged fluorescent proteins or a highly specific antibody and FACS. Regardless of the type of screening selection, NGS is used to compare the number of reads for each sgRNA in the perturbed experimental condition relative to a control to identify candidate genes for validation.

### Box 2: Considerations for setting screening parameters

Optimal screening parameters should maximize the difference in sgRNA distribution between the experimental and control conditions. Selection conditions such as drug dosage or FACS bin cutoff should be predetermined, if possible, using positive and negative controls from the literature and set to the level at which the greatest difference is observed. As for determining the duration of the screen, collection of time points throughout the screen helps identify the best time point for harvesting and analyzing the screen. These time points are also informative for assessing whether it is necessary to increase the duration to enhance the difference between experimental and control conditions.

Throughout the screen, it is imperative to maintain sufficient coverage to avoid losing sgRNA representation or bias the screening results. Try to maintain sufficient coverage at >500 cells per sgRNA in the library during library transduction, screening selection, and screening harvest. In addition, we recommend 2-4 infection replicates per screen to account for stochastic noise. Increase the coverage and number of infection replicates if the screening selection is noisy. Finally, consistency of screening conditions such as sgRNA representation and passaging reduces the variability between infection replicates.

### Box 3: Additional considerations for *ex vivo* and *in vivo* pooled screening

*Ex vivo* screening involves removing a primary cell type of interest from a living animal, culturing them in vitro and then performing the screen. For example, Parnas et al. demonstrated this strategy by deriving immune dendritic cells from Cas9 mice, transducing them with a CRISPR knockout library, triggering an immune response with lipopolysaccharide (LPS), and then FACS sorting different populations of cells based on immune response (e.g. TNF expression)^36^. This *ex vivo* screen identified many known as well as novel regulators of LPS response. When performing an *ex vivo* screen, it is necessary to be able to obtain enough cells to maintain library representation, deliver appropriate reagents to the cells, and culture the cells for long enough to perform the screen. In cases where these conditions cannot be met, adapt the screening strategy by, for instance, reducing the library size to capture a subset of genes.

*In vivo* screening is performed with either a) transduction of cells *in vitro* followed by *in vivo* cell transplantation, or b) direct transduction of tissues *in vivo*. The first strategy was demonstrated by Chen et al., whereby a cancer cell line was transduced with a CRISPR knockout library and injected subcutaneously in immunocompromised mice^34^. NGS analysis of harvested tumors identified known and novel tumor suppressors associated with tumor growth and metastasis. The main challenge of this approach is engrafting cells *in vivo*. Special care must be taken to ensure that the library is not only maintained upon infection of cells *in vitro* but also after engraftment of cells *in vivo*. While it is not required to maintain library representation on a per animal basis, a sufficient number of animals should be used such that library representation is maintained for each experimental cohort. Because the engraftment efficiency and time of engraftment can change for each application it is necessary to sequence the library at several time points after injection of cells in vivo. The optimal time point is one where engraftment is complete and selection (i.e. proliferation, death, or migration) has not yet occurred. Identifying this time point is critical as it is used as a reference to identify enriched and/or depleted perturbations.

For the second method of *in vivo* screening, special considerations will vary widely depending on the specific animal model, tissue, cell type, developmental time point, or biological question. Thus, each screen should be uniquely designed. In addition to the screening considerations outlined previously, the additional challenge for this strategy is the delivery of reagents in a complex environment while maintaining library representation and also infecting cells at a low MOI. Beyond specific circumstances, it may not be feasible to achieve appropriate cell numbers suitable for a genome-scale library. In these cases it is recommended to design smaller, targeted libraries with a specific hypothesis in mind. The complexity of the in vivo environment makes it difficult to meet the critical requirements for performing an informative screen. In assessing whether a direct in vivo screening strategy is feasible for any particular application, consider these guiding questions: 1) Is there a delivery strategy for infecting the target cells at low MOI? 2) Can enough of the target population be infected and purified to maintain library representation? 3) Can a reference population be identified before the guide RNA abundance changes?

### Box 4: Designing and analyzing a saturated mutagenesis screen

Although most pooled CRISPR screens to date have focused on knockout or activation of protein-coding genes, CRISPR screens can also be used to identify functional elements in noncoding regions of the genome such as enhancers or repressors. These functional elements are often inferred using biochemical hallmarks associated with function (e.g. chromatin accessibility, transcription factor binding sites, or post-translational histone modifications). In contrast, CRISPR screens enable direct testing of how mutagenesis at a specific noncoding site impacts phenotype.

Several strategies can be used to design libraries to target noncoding regions. For understanding regulation of a particular gene, tiling mutagenesis libraries were designed to include many or all possible target sites within a noncoding region near a gene^62^. This allows unbiased identification of all regulatory elements in regions near a gene that has already been established to be important for the screening phenotype (see illustration). Another approach is to design the library to target all instances across the genome of a specific biochemical hallmark, such as all binding sites of a transcription factor like p5 3 ^63^. With this kind of library, it is possible to identify specific binding sites or regulatory elements associated with a phenotype of interest.

As with screening the coding genome, a key factor in assessing the performance of the screen is to find multiple sgRNAs targeting the same element that are enriched or depleted together. In coding regions, this is straightforward, as the library is designed to have multiple sgRNAs that target the same gene. In noncoding regions, the same principle of consistent enrichment or depletion can be applied to multiple sgRNAs that target neighboring regions, as the indels are of variable length. Once a functional element is validated using multiple sgRNAs with adjacent target sites, expression of nearby genes and potential mechanisms such as alterations in transcription factor binding at the site can be used to gain further insight into biological mechanisms.

### Box 5: Lentivirus production with Polyethylenimine

1. Prepare polyethylenimine (PEI) transfection reagent:

a. Dissolve 50 mg PEI Max in 45 ml UltraPure Water.
b. Adjust pH to 7.1. First add 10 M NaOH dropwise until the pH approaches 6 and then add 1 M NaOH dropwise until final pH reaches 7.1.
c. Adjust final volume to 50 ml with UltraPure Water.
d. Sterilize using Millipore’s 0.45 μm Steriflip filter.
e. Prepare 50 × 1 ml aliquots and store at −20 °C until use. Note: PEI is stable for up to one year and can undergo 5 freeze-thaw cycles without a drop in transfection efficiency.
2. Prepare HEK293FT cells for lentivirus transfection as described in Steps 41-44.

a. For each lentiviral target, combine the following lentiviral target mix in a 50-ml Falcon tube and scale up accordingly:

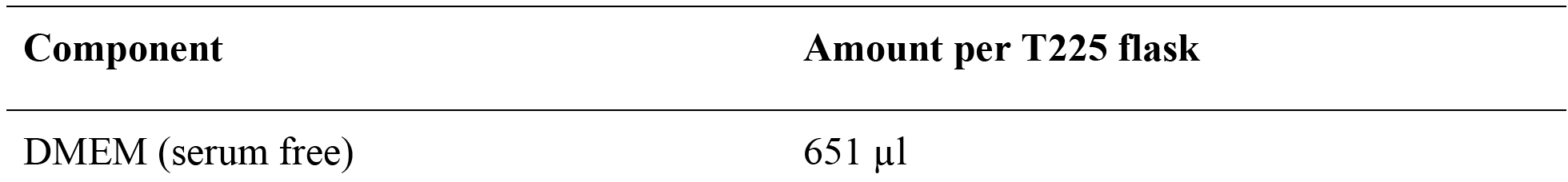

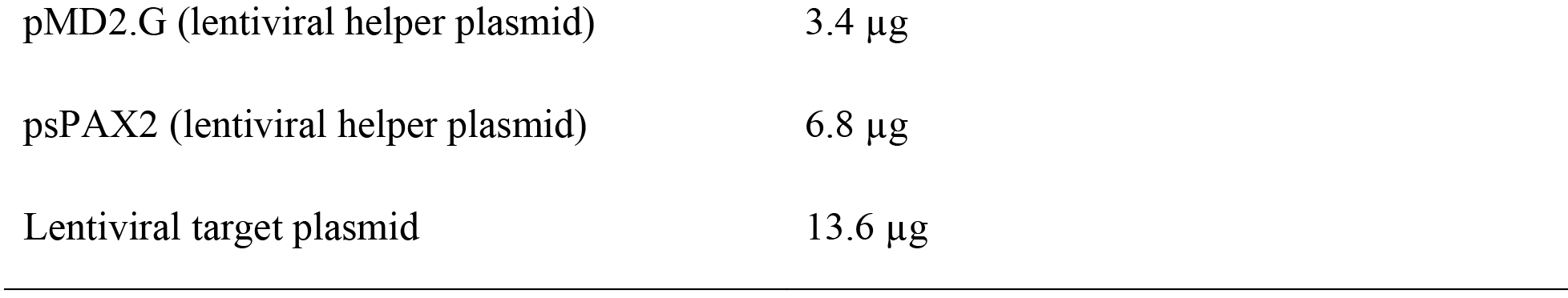
b. Add 195 μl PEI transfection reagent, vortex, and incubate at room temperature for 10 min.
c. Add 25 ml D10 media to the transfection reagent mixture.
d. Aspirate old media from cells, gently add new media containing the transfection reagent mixture and shake gently to mix. Return the T225 flask to the incubator.
3. 2 d after transfection, harvest and store lentivirus as described in Step 46.

### Box 6: Frequently asked screening questions from the CRISPR Forum

The following questions are selected from the CRISPR Discussion Forum (discuss.genome-engineering.org).

Q1: Can I use liquid culture amplification of the library rather than solid plates?

We recommend using plates because liquid culture can generate more bias in the plasmid library. Beta-lactamase, the enzyme responsible for ampicillin resistance, is secreted and eventually in liquid culture the selective pressure on the plasmid is decreased causing bias. Additionally, it is more difficult for certain clones to predominate on solid plates because they are spatially limited in growth and each clone is spatially separated to prevent potential intercolony competition. However, it is important to note that some studies have had success with liquid culture amplification^38^.

Q2: Is there a difference between using HEK293FT vs HEK293T cells for lentivirus production?

Yes, HEK293FT cells are generally more ideal for lentivirus production. HEK293T cells are a cell line stably expressing the SV40 large T antigen, which helps to boost protein production off expression constructs containing the SV40 enhancer element. HEK293FT cells are a fast-growing, highly transfectable clonal derivation of HEK293T cells that yield higher lentivirus titer than the HEK293T line.

Q3: For activation, how do I design guides relative to the TSS of the transcript? Additionally, can I expect these guides to work with transient transfection of dCas9-VP64 and MS2-VP65-HSF1 plasmids?

The TSS is the first base of the transcript, i.e. beginning of the 5’ UTR. The UCSC table browser is a good resource for TSS annotations. We have observed the most robust transcriptional upregulation when sgRNAs are designed to target the 200bp region upstream of the TSS. We have created a web tool using these parameters to simplify activation sgRNA design for human and mouse genes (http://sam.genome-engineering.org/database/). SAM is highly robust and should yield significant activation levels even in the case of transient transfection^50^.

Q4: What are important considerations for NGS PCR amplification?

When designing primers, it is important to include stagger between the primer binding site and the Illumina adapter sequence such that the sequencing regions of different amplicons are offset, improving the sequence diversity and quality. For genomic DNA amplification, it can be helpful to optimize the DNA input for the sequencing readout PCR step. Generally, it is recommended for any given instance of the screen to titrate the DNA input and use the highest possible input without a decrease in the target band intensity. It is critical to minimize amplification bias. The optimal cycle number should always be determined by doing a series of different cycle numbers (e.g. 5, 10, and 15) and identifying the lowest cycle number that generates a visible band by gel electrophoresis. Avoid conditions that yield additional bands at higher cycle numbers.

Q5: My screening design requires too many cells. Can I reduce the coverage?

We recommend screening at a coverage of >500 cells per sgRNA. Because there is always variability in the copy number of each sgRNA in a given library, it is important to have high coverage to overcome any bias. If it is impossible to screen at this coverage (e.g. insufficient primary cells or cells are difficult to transduce), consider screening with a smaller, targeted library.

Q6: How do you measure the quality of a cloned plasmid library?

While there are many methods for determining the quality of a library, we typically use the following measures for a sequencing depth of >100 reads per sgRNA:

1. Overall representation: <0.5% of sgRNAs have dropped out with no reads.
2. Library uniformity: <10-fold difference between the 90th percentile and the 10th percentile.

**Table S1.**
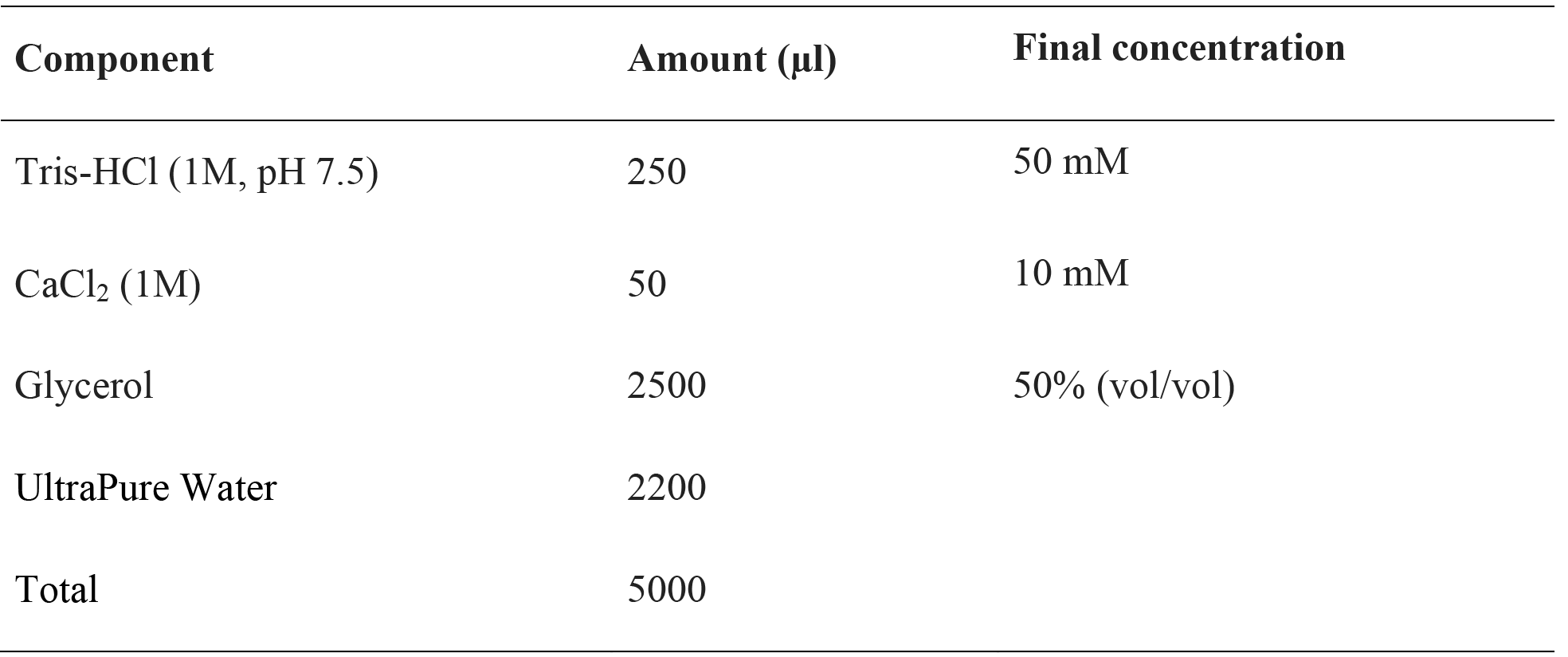
Deoxyribonuclease I storage solution. Solution for resuspending and storing deoxyribonuclease I for fold activation analysis during validation. Prepared solution can be stored at −20 °C for up to 2 years.

**Table S2.**
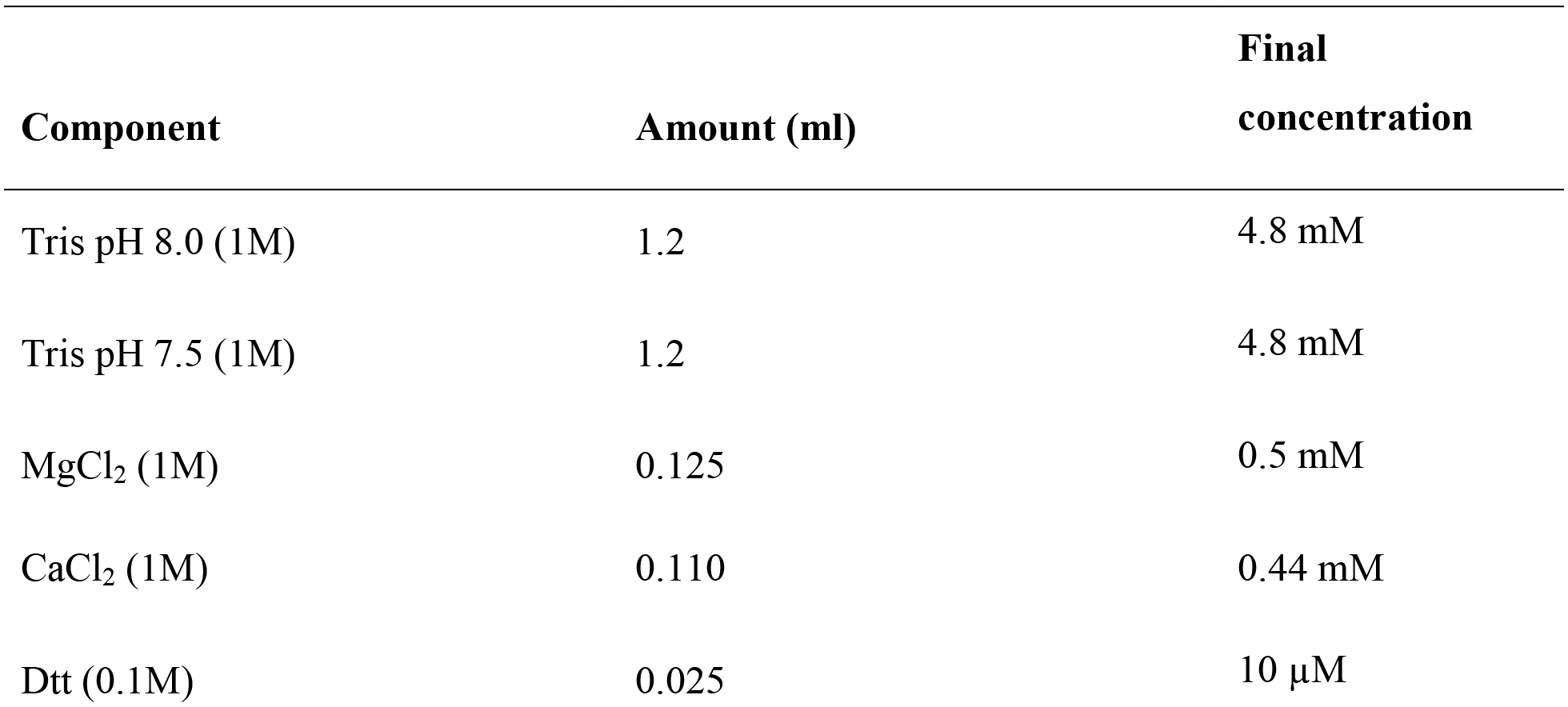
RNA Lysis Buffer Setup. Buffer for lysing and harvesting RNA from cells for fold activation analysis during validation. Final pH of the solution should be approximately 7.8

**Table S2.**
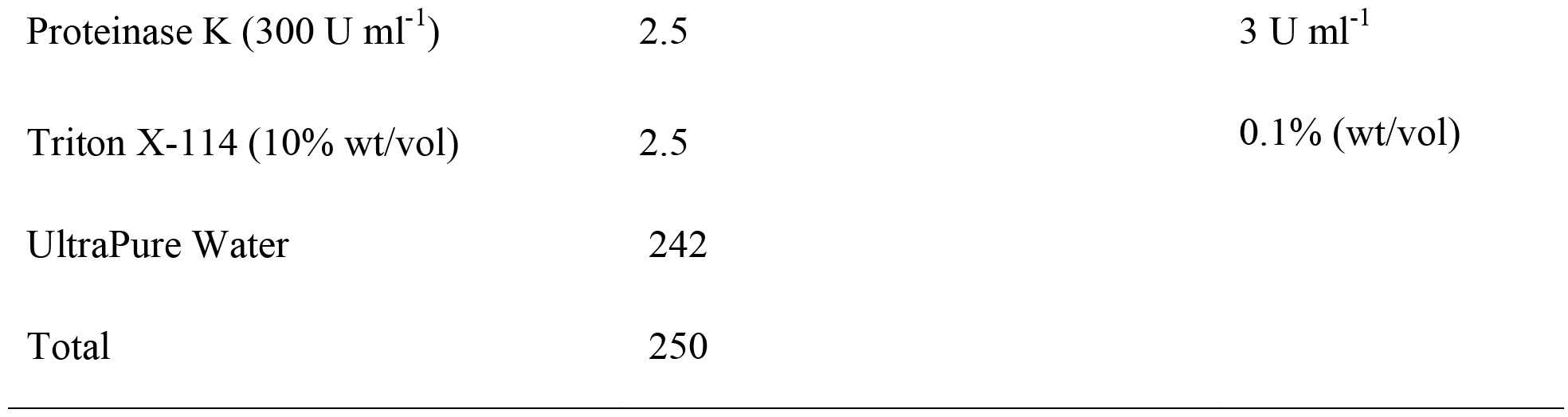

**Table S3.**
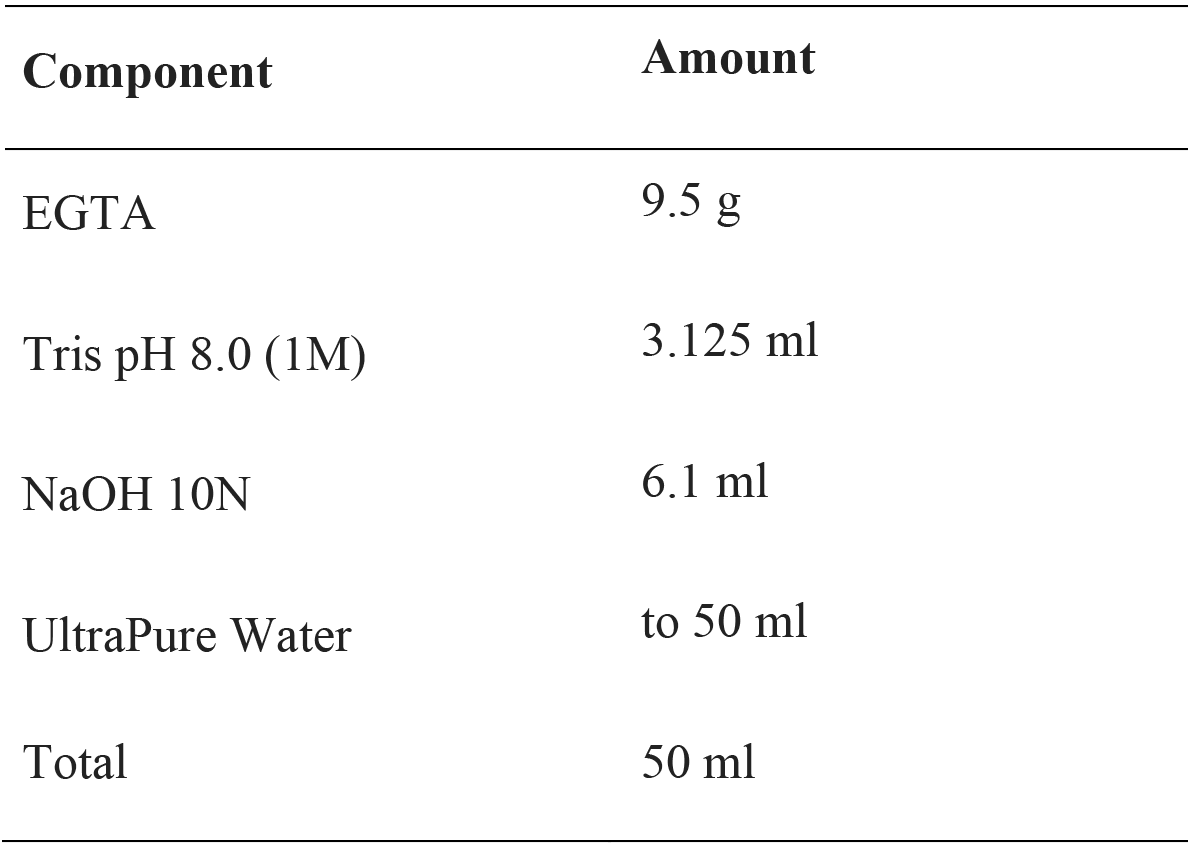
EGTA (0.5M, pH 8.3) stock. EGTA stock solution used to prepare RNA lysis stop solution for fold activation analysis during validation. Adjust with NaOH to a final pH of 8.3 if necessary. Take aliquots to measure pH in order to keep main stock from being RNAse-contaminated by the pH probe.

**Table S4.**
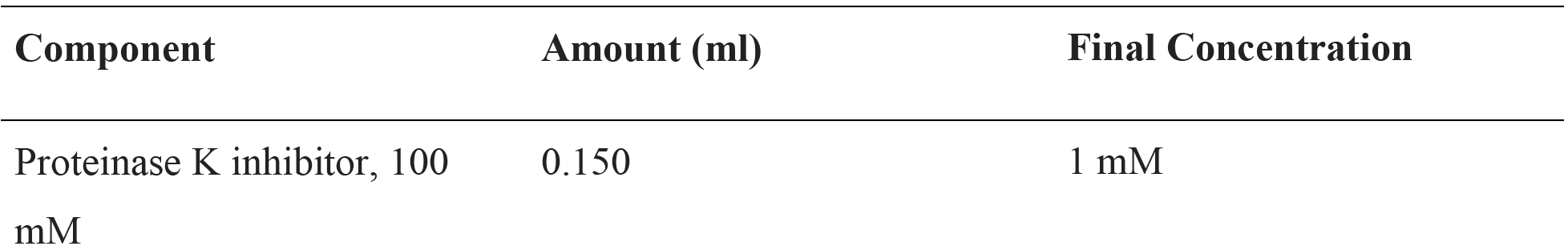
RNA lysis stop solution. Solution for terminating RNA lysis for fold activation analysis during validation. Prepare using EGTA (0.5M, pH 8.3) from **Table S3**.

**Table S4.**
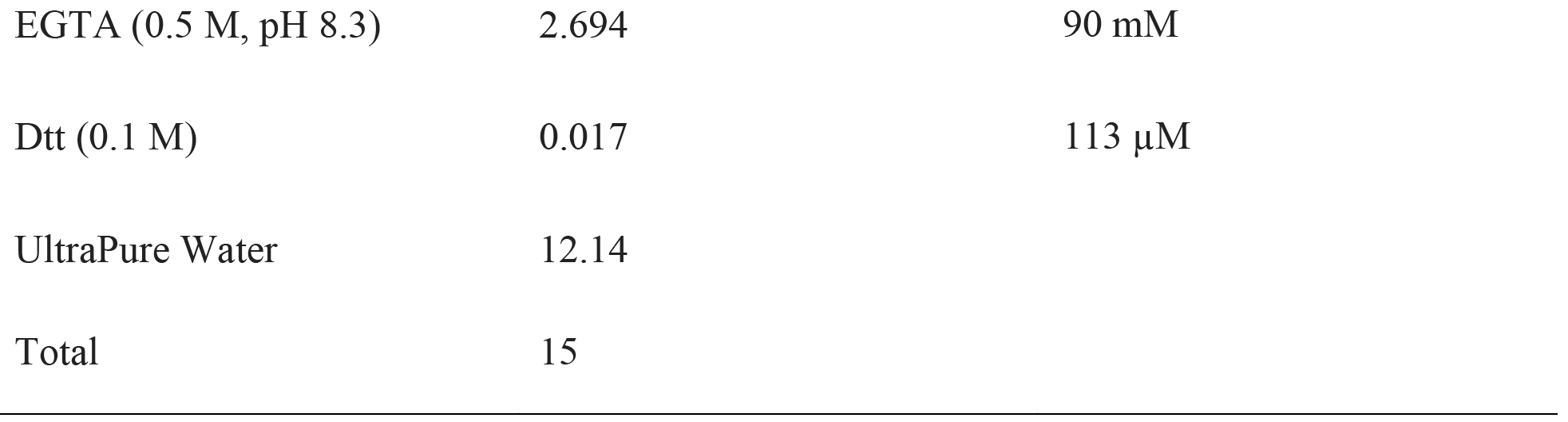

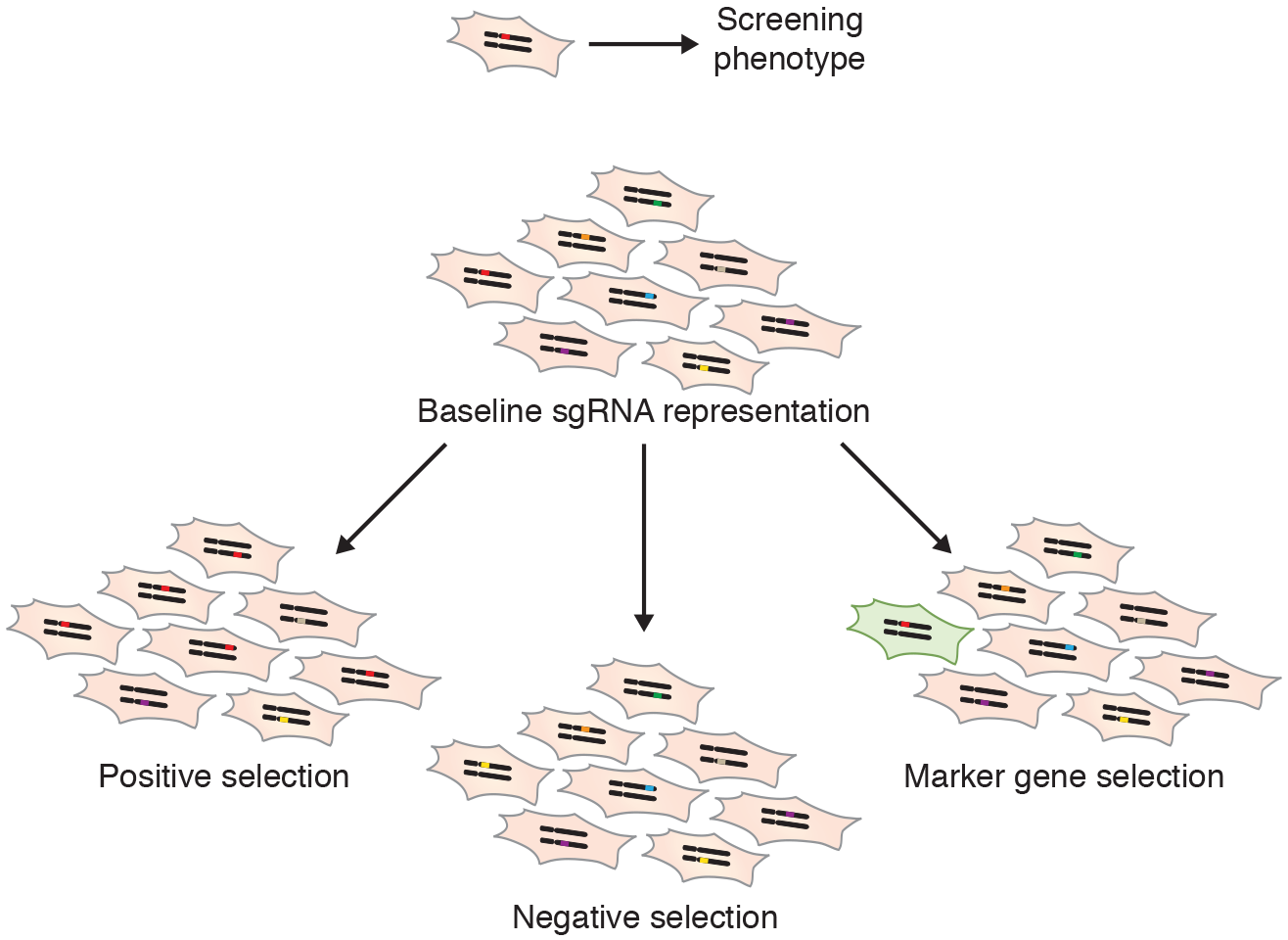

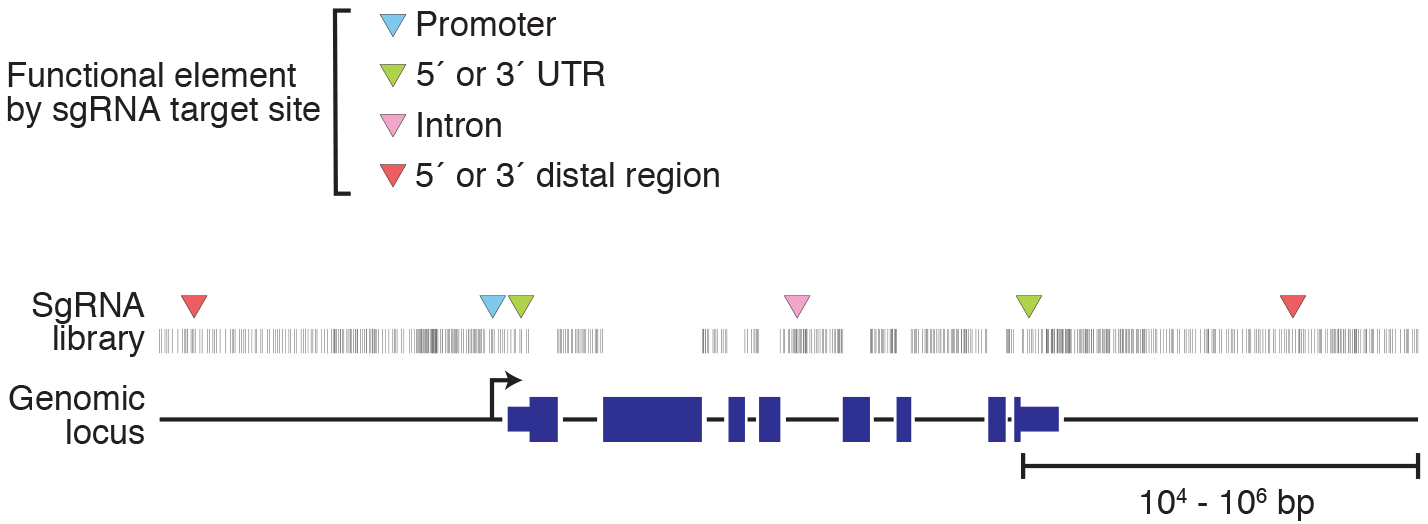

